# Intact rDNA arrays of *Potentilla*-origin detected in *Erythronium* nucleus suggest recent eudicot-to-monocot horizontal transfer

**DOI:** 10.1101/2021.12.15.472635

**Authors:** László Bartha, Terezie Mandáková, Aleš Kovařík, Paul-Adrian Bulzu, Nathalie Rodde, Václav Mahelka, Martin A. Lysak, Margaux-Alison Fustier, Jan Šafář, Petr Cápal, Lujza Keresztes, Horia L. Banciu

## Abstract

The occurrence of horizontal gene transfer (HGT) in Eukarya is increasingly gaining recognition. Nuclear-to-nuclear jump of DNA between plant species at high phylogenetic distance and devoid of intimate association (e.g., parasitism) is still scarcely reported. Within eukaryotes, components of ribosomal DNA (rDNA) multigene family have been found to be horizontally transferred in protists, fungi and grasses. However, in neither case HGT occurred between phylogenetic families, nor the transferred rDNA remained tandemly arrayed and transcriptionally active in the recipient organism. This study aimed to characterize an alien eudicot-type of 45S nuclear rDNA, assumingly transferred horizontally to the genome of monocot European *Erythronium* (Liliaceae). Genome skimming coupled by PacBio HiFi sequencing of a BAC clone were applied to determine DNA sequence of the alien rDNA. A clear phylogenetic signal traced the origin of the alien rDNA of *Erythronium* back to the Argentea clade of *Potentilla* (Rosaceae) and deemed the transfer to have occurred in the common ancestor of *E. dens-canis* and *E. caucasicum.* Though being discontinuous, transferred rDNA preserved its general tandemly arrayed feature in the host organism. Southern blotting, molecular cytogenetics, and sequencing of a BAC clone derived from flow-sorted nuclei indicated integration of the alien rDNA into the recipient’s nuclear genome. Unprecedently, dicot-type alien rDNA was found to be transcribed in the monocot *Erythronium* albeit much less efficiently than the native counterpart. This study adds a new example to the growing list of naturally transgenic plants while holding the scientific community continually in suspense about the mode of DNA transfer.

**Significance Statement:** Ribosomal DNA is an essential component of all cellular genomes. In plants, accidental movement of rDNA via horizontal gene transfer has only been reported in sexually incompatible grasses (monocots) where it involved non-functional rDNA units. In this study, we propose that evolutionary trajectories of eudicots and monocots were bypassed by the jump of rDNA from a *Potentilla* species (Rosaceae) to a common ancestor of *Erythronium dens-canis* and *E. caucasicum* (Liliaceae). The alien eudicot-type rDNA appeared relatively well conserved in the examined host *Erythronium* genome, being able to be expressed while preserving its general tandemly repeated feature, evidences that have no match in earlier literature.

## Introduction

Horizontal gene transfer (or alternatively, lateral gene transfer, LGT) is an evolutionary mechanism surmounting reproductive barriers between species. It refers to the movement of genetic material between organisms through routes other than parent-to-offspring. The phenomenon is widespread in prokaryotes and its significance and prevalence are also increasingly recognized in eukaryotes (1, 2). Successful gene transfers can result in evolutionary innovations, like adaptation to a new environment (3), improvement of parasitic lifestyle (4), or a shift in metabolism resulting in C4 biochemistry (5).

Concerning plant-to-plant gene transfers, the very first examples of HGT referred mostly to the jump of mitochondrial DNAs within host-parasite systems [see review (6)]. Later, reporting HGT of nuclear genes and their expression in parasitic plants also became mainstream (7). Not surprisingly, the host-parasite interface provides a suitable opportunity for gene movement via the intimate physical connections between donor and recipient organisms. More recently, HGT was shown to frequently occur in certain groups of plants like grasses where it was evidenced that phylogenetic proximity and rhizomatous nature increased the frequency of HGT (8).

Nucleus-to-nucleus jump of DNA among angiosperms that are found at high phylogenetic distance (e.g., those belonging to different families) and not showing some kind of biological promiscuity (e.g., parasitism) are still rarely reported. Indeed, transfer of a nuclear-encoded transposable element (TE) between the dicot grapevine and the monocot palm tree has been reported (9). These genomic components are thought to be particularly prone to HGT thanks to their propensity for self-replication, extrachromosomal stage from within their transposition cycle, and the capacity to integrate themselves into host chromosomes (10–12).

Ribosomal DNA constitutes an essential component of all cellular genomes. It encodes ribosomal RNA (rRNA) molecules that together with ribosomal proteins make up ribosomes, factories for protein synthesis. In the majority of angiosperms, three of the RNA genes [18S, 5.8S, and 25S (26S)] are physically linked in a conserved cluster referred to as 45S rDNA. The above genes are separated by two internal transcribed spacers (ITS1 and ITS2) and altogether form a single transcription unit (13, 14). The units are arrayed tandemly and are separated by the so-called intergenic spacer (IGS) region, the 3’-end of which is commonly referred to as external transcribed spacer (ETS) (14).

In plants, intra-individual heterogeneity of rDNA is a well-documented feature. Withing a genome, different rDNA units (e.g., those resulting from hybridization or pseudogenization) can be homogenized by concerted evolution via unequal crossing-over and gene conversion (15, 16). The propensity to maintain or erase intra-individual rDNA heterogeneity/paralogy seems to vary across genomic and taxonomic contexts. As such, rDNA can be particularly polymorph in ancient plant lineages (17, 18) but in the same time, it can be subjected to rapid unidirectional gene conversion in recently formed allopolyploids (19, 20).

According to the ‘complexity hypothesis’ (21), members of complexes involved in the transcriptional and translational processes (a.k.a. ‘informational genes’) are less prone to HGT when compared to the ‘operational genes’ (i.e. ‘metabolic housekeeping genes’). Indeed, albeit HGT of rDNA has been repeatedly documented in prokaryotes (22–24), it was shown to occur at a low rate and generally was limited to take place between closely related taxa upon a survey of 2143 prokaryotic genomes (24). Within eukaryotes, HGT of 45S rDNA elements has been revealed among protists (25), fungi (26) and monocot plants (grasses) (27). Published evidences of horizontally transferred eukaryotic rDNA concern only jumps at the intrafamily level and, if tested, in none case the transferred rDNA was shown to be transcriptionally active in the recipient organism. Moreover, the general tandemly arrayed nature of eukaryotic 45S rDNA (28) was not yet confirmed in any host organism after its transfer.

The genus *Erythronium* belongs to the monocot family Liliaceae and includes ca. 30 species of bulbous geophytes confined mainly to temperate forest and damp meadow habitats of the northern hemisphere (29, 30). Species of the genus cluster in three well supported and biogeographically meaningful (eastern North American, western North American and Eurasian) lineages (29, 30). During a phylogenetic study (31) of Eurasian *Erythronium*, Sanger sequencing of the ITS region of 45S nuclear rDNA in *Erythronium dens-canis* L. did not provide sequences of satisfactory quality. This led to the necessity of cloning of the ITS amplicons. Surprisingly, upon BLAST searches, a subset of the sequenced clones showed 97 percent identity with ITS sequences of the eudicot (hereafter for simplicity named dicot) genus *Potentilla* (Rosaceae). This finding hinted at either a possible artefact, or the actual presence of *Potentilla*-type DNA within the genome of *Erythronium.* A series of experiments conducted subsequently in different laboratories supported the latter scenario, providing evidence for sharing genetic material between plant lineages which separated ca. 152 Mya (http://www.timetree.org/) (32). Hence, in this study, we summarize the results of analyses aiming to characterize the nature of *Potentilla*-specific rDNA within the genome of the monocot plant *Erythronium.* Particular challenges of documenting alien rDNA in the host genome were raised by the general repetitive nature of rDNA and by the large genome of *Erythronium* combined with the relative low amount of alien rDNA therein. A previous genome size measurement of the species estimated a monoploid size (1Cx) to approx. 12.2 gigabase pairs (Gbp) (33), which was more than twice the mean angiosperm 1C value reported recently (34).

## Results and Discussion

### rDNA Jumped Between Plant Families

Throughout the study, for consistency purposes an effort was made to conduct key analyses on specimens from a single population of *E. dens-canis*. For this purpose, we selected the so-called ‘Feleacu’ population from NW Romania (Table S1). Endeavors to differentially PCR-amplify *Potentilla*-specific ITS2 in selected New and Old World *Erythronium* taxa resulted in the expected product only in case of *E. dens-canis* and *E. caucasicum* Woronow (Fig. S1). Besides the initial cloning of ITS in *E. dens-canis*, additional strategies for retrieving and tracing back the origin of the alien rDNA included *de novo* and reference-guided assemblies of ITS and partial ETS from genomic Illumina reads. Furthermore, a low-coverage bacterial artificial chromosome (BAC) library was constructed from flow-sorted nuclei of *E. dens-canis* and screened for *Potentilla*- and *Erythronium*-specific ITS2. The two sequenced BAC clones, 1J24 and 18I01, contained alien and native rDNA, respectively, from which ITS and partial ETS sequences were extracted for phylogeny purposes. Alien-type ITS sequences of *Erythronium* showed 97% identity with *Potentilla* in BLAST search, pointing to this genus as a likely source of the sequences. A follow-up ML phylogenetic analysis placed the sequences with maximal statistical support within the Argentea clade of the genus (Fig. 1).

**Figure 1.**
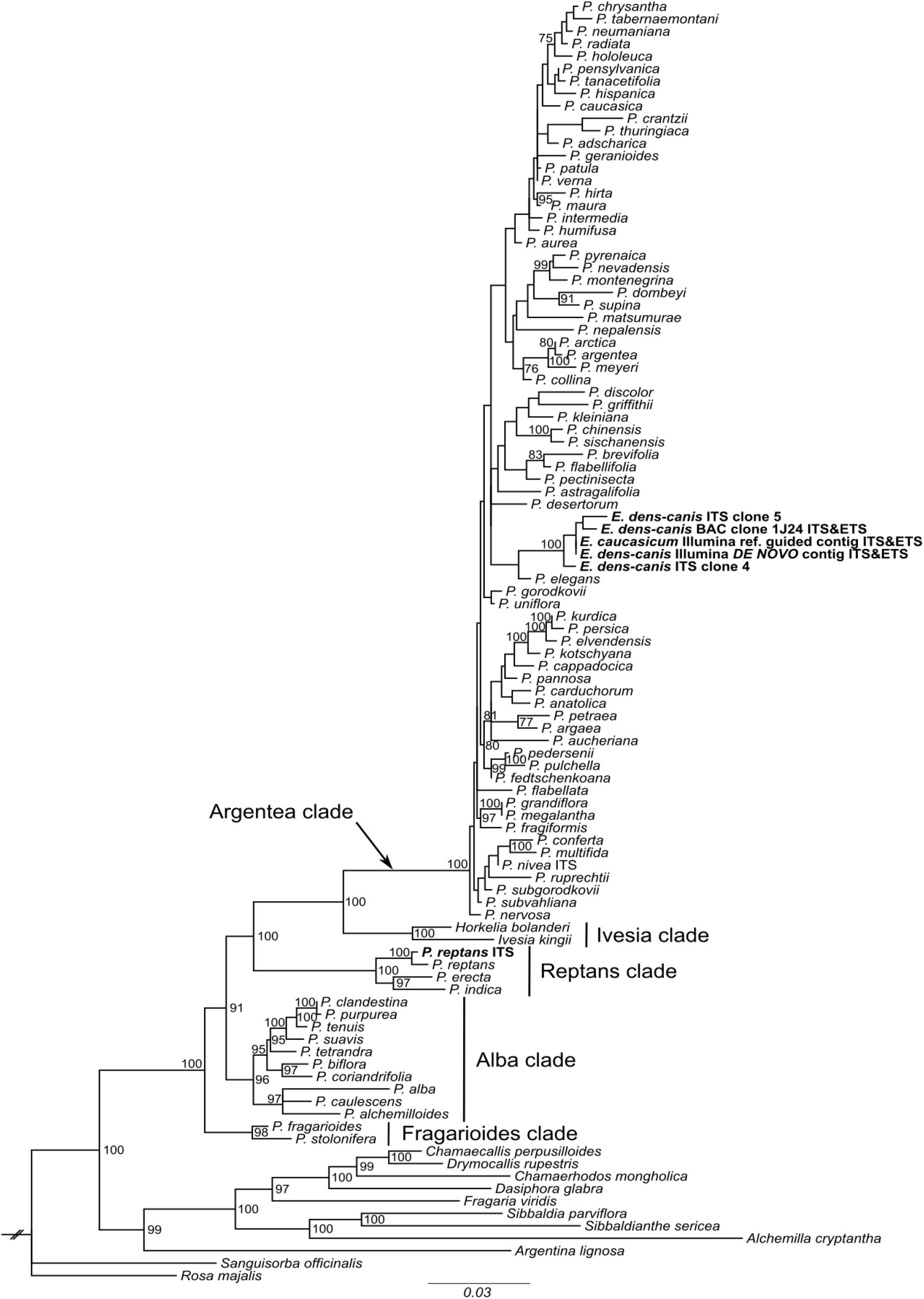
IQ-TREE ML phylogram based on *Potentilla* and *Potentilla*-specific *Erythronium* ITS and partial ETS sequences. Terminal names with newly generated sequences are in bold. Nodal support percentages (not shown below 75) were derived from 1000 ultrafast bootstrap [UFBoot (72)] replicates.

The native *E. dens-canis* ITS from BAC clone 18I01 fitted well the previously published Eurasian *Erythronium* phylogeny (31), serving as a general control (Fig. S2). In nuclear ITS and plastid DNA-based phylogenies of the genus, *E. dens-canis* and *E. caucasicum* were consistently resolved as sister taxa (29–31). Additionally, populations of *E. dens-canis* subdivide into the so-called ‘Transylvanian’ and ‘non-Transylvanian’ phylogeographic lineages based on plastid DNA (35). The ‘Feleacu’ population belongs to the ‘Transylvanian lineage’ of the species. In order to enlarge the coverage area of the study, four *E. dens-canis* specimens from different western and central European populations [all belonging to the ‘non-Transylvanian lineage’ (35)] were additionally spot-checked for *Potentilla*-type ITS2. In all cases the presence of alien ITS2 was confirmed by direct sequencing (Table S1). The phylogenetically meaningful presence of *Potentilla*-specific rDNA in European *Erythronium* can be best interpreted as a HGT event preceding the split of *E. dens-canis* and *E. caucasicum.* One can speculate that alien rDNA is present in most of the populations of *E. dens-canis* across its latitudinally narrow and longitudinally wide European range. Testing this assumption, however, was beyond the scope of the present study.

The invoked ML phylogeny confidently traced the origin of alien rDNA back to the donor organism at the generic level. *Potentilla* is a species-rich genus comprising ~400 typically yellow-flowered representatives in the Northern Hemisphere (36). Most of the members (~300 spp.) belong to the informally named Argentea clade (37), in which species relationships are generally unresolved [(38), Fig. 1]. Illumina consensus sequences of alien ITS from *E. dens-canis* and *E. caucasicum* are identical, pointing to their recent acquisition from *Potentilla* and conserved nature in *Erythronium*. Since the sequences do not cluster with statistical significance with any current *Potentilla* species included into the analysis, the donor remains unknown. Revealing the donor’s identity or narrowing relationships of the donor organism are challenging tasks, that can potentially be addressed in the future with the aid of more variable genetic markers and/or by increasing sampling of ITS sequences from the genus of *Potentilla.*

### Alien rDNA is integrated in the nuclear genome of *Erythronium*

Importantly, a series of experiments was conducted to unambiguously prove the insertion of alien rDNA in the host *Erythronium* genome. The experiments led to the elimination of both possible contamination of DNA or samples, and potential involvement of organellar genomes as the locations of alien DNA.

A Southern blot technique was employed using *Apo*I-digested genomic DNAs of samples of *E. dens-canis* as the host genome, complemented by *Erythronium sibiricum* subsp. *altaicum* Rukšāns (later for simplicity mentioned as *E. sibiricum*) used as a negative control and *Potentilla reptans* L. used as a positive control. Digested DNA of *E. dens-canis* produced a distinct band of ~0.9 kilobase pairs (kbp) when hybridized to either *Potentilla*-specific ITS1-5.8S-ITS2 or ITS2 probes. This result fitted expectations based on the restriction map (Fig. 2). This band of *Potentilla*-origin was missing in *E. sibiricum*. At the applied default stringency recommended by the labeling kit producer, probes were not able to discriminate between the native and alien rDNA fragments resulting from the digestion. Specificity was therefore guaranteed by differences in band sizes. Due to the high sequence conservation of 5.8S regions between *Erythronium* and *Potentilla*, binding of the longer probe also to the ~2.4 kbp long native rDNA fragment of *Erythronium* was expected. On the other hand, hybridization of *Potentilla* ITS2 probe to this fragment was somehow surprising. It seems that the 51% sequence identity (over 260 bp) between *Erythronium* and *Potentilla* ITS2 hindered the design of a fully specific probe. At least the bands are more clear-cut in the case of the shorter probe, indicating an improvement in specificity. Smears in case of *E. sibiricum* may indicate that its rDNA is not homogenous and does not conform entirely to the predicted digestion pattern.

**Figure 2.**
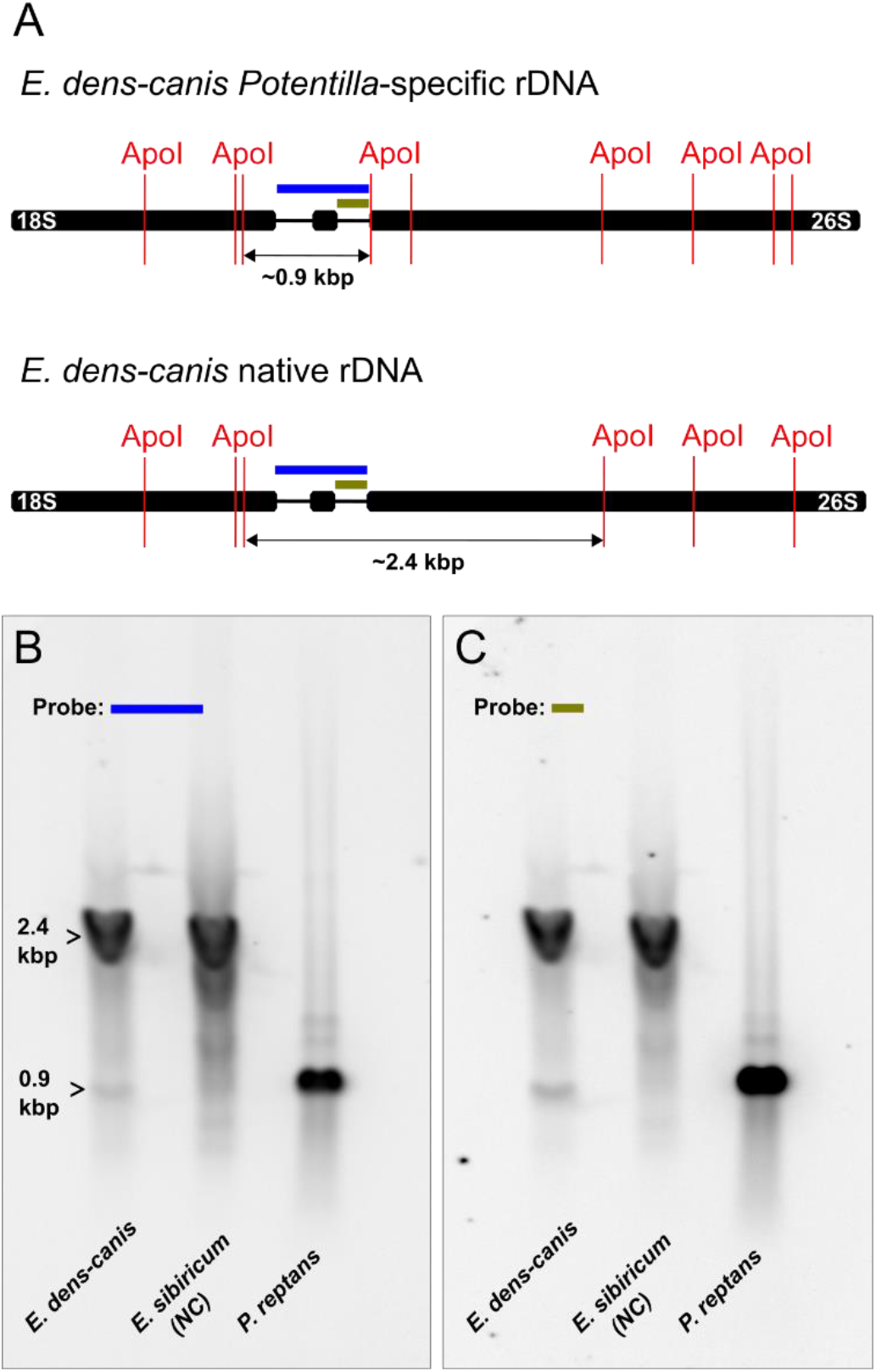
Southern blot analysis of rDNA fragments produced by *Apo*I-HF from gDNA of *E. dens-canis, E. sibiricum* and *P. reptans.* (A). Restriction map of *Apo*I-HF targeting the native and alien rDNA of *E. dens-canis.* Probes consisted of *P. reptans* ITS region (B) and *Potentilla-* type ITS2 cloned from *E. dens-canis* (C). Arrows indicate *Potentilla-* (~0.9 kbp) and *Erythronium*-specific (~ 2.4 kbp) rDNA fragments. NC, negative control.

Additional evidence of nuclear insertion of alien rDNA was provided by the alien rDNA-containing BAC clone 1J24. BAC library of *E. dens-canis* was constructed from flow-sorted nuclei, eliminating the possibility of *Potentilla*-rDNA to represent any kind of artefact.

Our efforts to directly visualize parts of alien rDNA using fluorescence *in situ* hybridization (FISH) technique failed. Probes either cross-hybridized with native loci (in case of ITS1 and ETS) or provided no signals (in case of 5’-end IGS). We therefore resorted to an indirect approach targeting native ETS and, additionally, native-alien 26S gene on *E. dens-canis* and *E. sibiricum* chromosomes. The 26S probe identified two minor terminal loci on two different, non-homologous chromosomes of *E. dens-canis* that were extraneous to the native ETS loci (Fig. 3). These can be interpreted as alien rDNA. Such additional loci were not observed in *E. sibiricum* (Fig. S3). We are aware of the fact that chromosomal locations in which native and alien rDNA co-localize could not be identified by this approach therefore the real number of alien rDNA loci could be higher.

**Figure 3.**
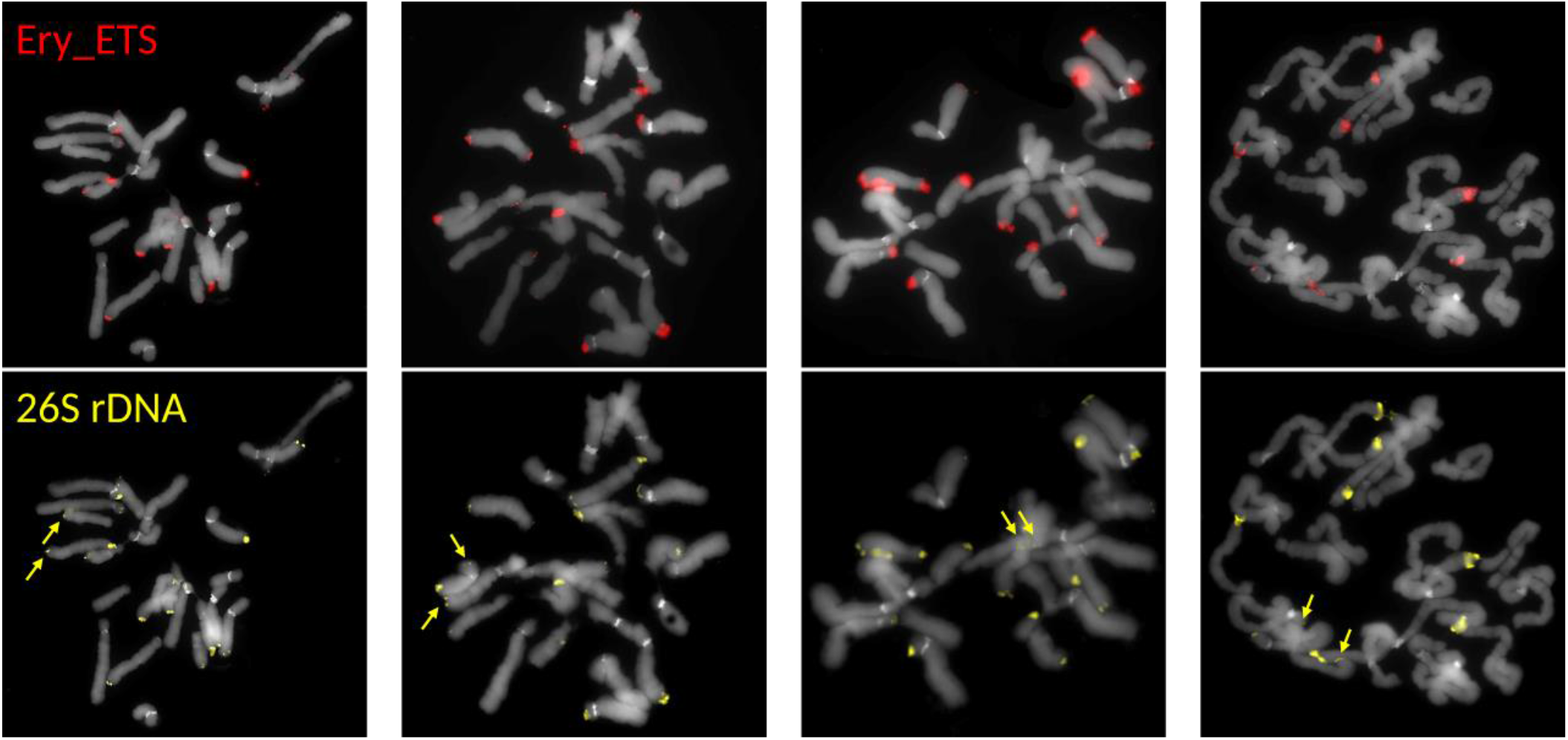
Chromosomal localization of rDNA in *Erythronium dens-canis*. Four mitotic chromosome complements were hybridized with *Erythronium* ETS (Ery_ETS, red signals) and *Nicotiana* 26S rDNA probe (yellow signals). Extraneous sites identified by the 26S probe (marked by arrows) are likely representing the sites of alien rDNA. Chromosomes were counterstained with DAPI.

### Alien Dicot rDNA Is Organized Tandemly in the Host Monocot Genome

Physical map of BAC clone 1J24 provides unprecedented insight into the organization of a dicot rDNA within the genome of a monocot plant (Fig. 4). BAC clone 1J24 contained eight transcription units of *Potentilla*-derived rDNA. Five out of the eight copies of 26S rRNA gene are disrupted by potentially coding sequences (CDSs) of opposite orientation. In one additional case, the 26S rDNA gene is interrupted by an insertion of sequence of unknown origin. The CDSs are not identical and all contain the domain of unknown function (DUF) 4283. Interestingly, similar hypothetical CDSs interleaving 26S rRNA gene were also found within a considerably longer array of the native rDNA of *E. dens-canis* found within the 125.9 kbp long BAC clone 18I01 (Fig. S4).

**Figure 4.**
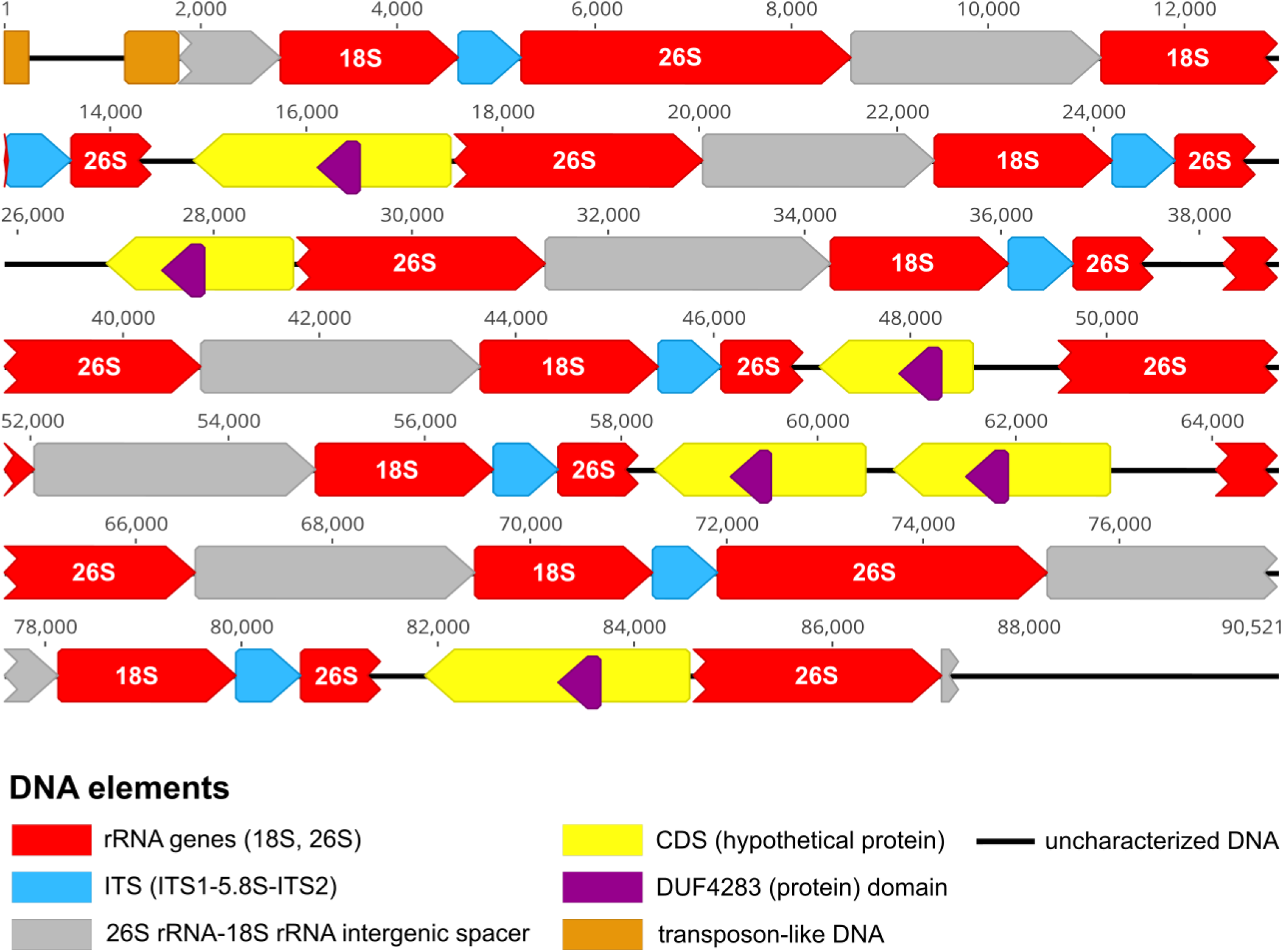
Physical map of *E. dens-canis* BAC clone 1J24 containing rDNA of exclusively *Potentilla*-origin.

A Neighbor-Joining tree was generated based on the alignment of translated DUF4283 sequences of the two BAC clones supplemented with publicly available similar protein sequences (Fig. S5). Sequences belonging to BAC 1J24 formed a clade to which the domain sequence of BAC 18I01 was resolved as sister. This suggests their homology. Unfortunately, the phylogeny is not robust enough to establish the phylogenetic origin (dicot versus monocot) of the BAC CDSs. Most of the participating sequences are hypothetical and/or uncharacterized, leaving us with a conundrum about the possible roles of these CDSs in the *Erythronium* genome.

In addition to potential CDSs, short (up to 531 bp long) transposon-like fragments were also identified in BAC clone 1J24 (Table S2). These, however, did not assemble (at least in this clone) into complete transposable elements. TEs have been the subject of HGT in plants relatively frequently and are viewed as notably prone to HGT (9–12, 39). Whether this relies on the fact that they exist as extrachromosomal elements during their transposition cycle, remains to be clarified (1). TEs are thought to be able to insert themselves into vector DNA and, subsequently, into host chromosomes (10). Once integrated into the host genome, they can undergo various processes, like mutation, epigenetic silencing, relocation or amplification (transpositional burst) (12). As such, they can reshuffle genomic regions (40). The fact that TEs often accompany foreign DNA in the host genomes (27, 41, 42) suggests that they could play a role in the DNA transfer. Moreover, DNA fragments similar to the *Potentilla*-sister *Fragaria* DNA transposon CACTA elements found within BAC clone 1J24 (Table S2) suggest transposable elements might have also been participating in the actual transfer.

Simultaneous presence of native and alien rDNA was not detected in the sequenced BAC clones although a chimeric sequence read containing parts of both native and alien rDNA (Fig. S6 and Dataset S1) was found within the Nanopore whole genome shotgun (WGS) dataset (Tables S3, S4). The fragmentation of native rDNA observed in this sequence is possibly result of pseudogenization. Nevertheless, given the rarity of this sequence type, one cannot rule out its origin as a sequencing artefact. Presumed pseudogenization of alien rDNA at a larger scale was caught in action in the longest Nanopore read containing *Potentilla*-specific rDNA (Fig. S7 and Dataset S2). Here, disruption of rDNA subunits has a more or less symmetrical pattern within a palindromic structure that renders little likelihood for the artefact origin. Numbers of polymorphic nucleotide sites were also assessed in alignments of native and alien 18S and 26S sequences from within the two analyzed BAC clones. Pseudogenization of alien rDNA is further supported by the fact that number of polymorphic nucleotide sites is several folds higher in alien 18S and 26S sequences when compared to their native homologues (Table S5). Still, conclusions in this respect must be taken cautiously since nucleotide variation in two BAC clones cannot truly reflect variation of target regions across the whole-genome level.

Albeit the presence of horizontally-transferred rDNA was demonstrated previously in eukaryotes (25, 27, 43), in none of the cases the detected alien DNA was arrayed like shown in the present study.

### Alien rDNA Is Expressed in *Erythronium*

The presence of tandemly arrayed alien rDNA units in the *Erythronium* genome raised the possibility of their transcription. To test this supposition, RNA was extracted from actively growing roots of a cultivated *E. dens-canis* plant from the ‘Feleacu’ population, reverse-transcribed, and subjected to a regular PCR by using *Potentilla* ITS2-specific primers (Table S6). The experiment showed that the alien rDNA was transcriptionally active in *Erythronium* roots (Fig. 5). This finding, however, does not necessarily mean that expressed *Potentilla*-specific rRNA is ‘biologically active’ in the sense of taking part in the protein synthesis machinery of the host plant. Alien and native 18S and 26S RNAs might be too diverged (95.7 and 90.5 percent identities, respectively) to be interchangeable within a fully functional ribosome. Ribosomal heterogeneity is a well-documented concept and although it can be defined by rRNA sequence variation, the exhaustive list of previously published examples covers mainly heterogeneity determined by ribosomal protein paralogy (13). No precedent in the literature can be found about 18S or 26S rRNA variations leading to deviation of canonical ribosomal structure or differences in translation efficiencies between the uses of different rRNAs by ribosomes of the same plant. In any case, transcription of assumedly horizontally transferred rDNA found within the current study is unprecedented in the literature.

**Figure 5.**
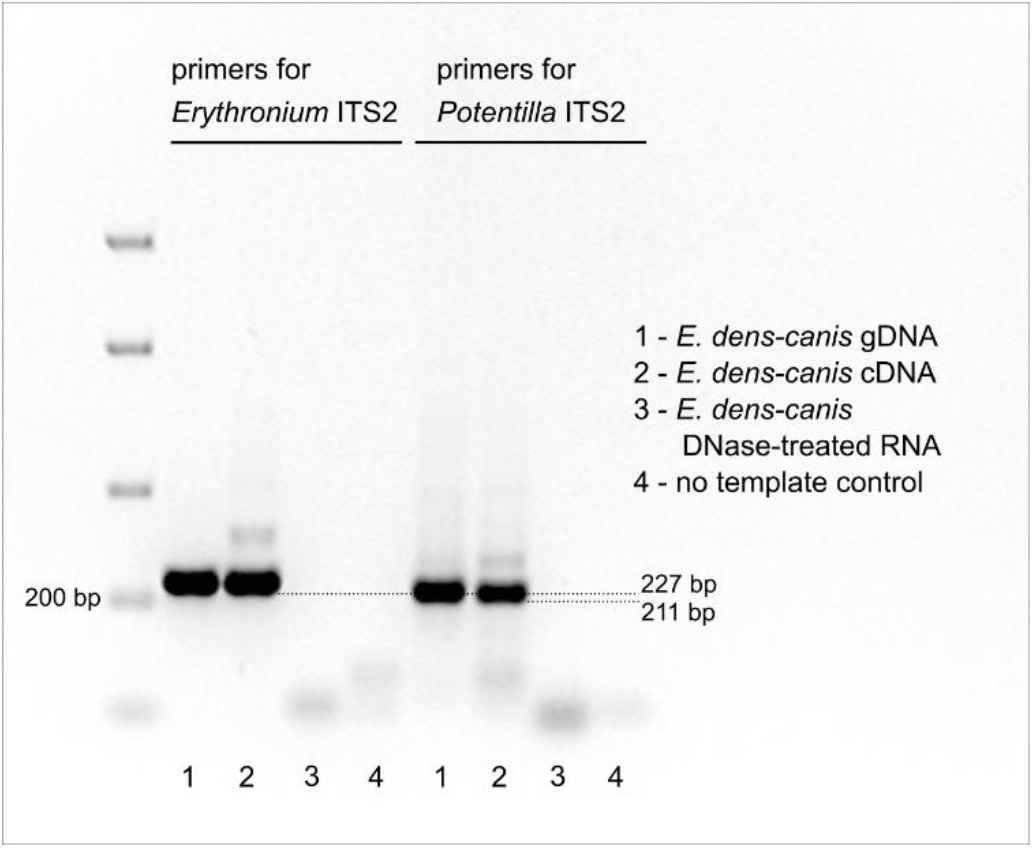
Evidence of transcription of alien rDNA in *E. dens-canis* based on the amplification of *Potentilla*-specific ITS2 from cDNA. PCR products were visualized according to ethidium bromide-UV detection and were based on the templates and primers indicated in the body of figure. ITS2 fragment sizes include lengths of primers.

Another issue that was intriguing to explore was the amount of the alien rDNA in the host genome. Specifically, we measured relative fold difference (RFD) between the native and its alien homologue of rDNA within the host genome and transcriptome. A semi-quantitative PCR (SQ-PCR) approach was carried out to perform pilot estimates of this ratio in *E. caucasicum* gDNA, as well as in *E. dens-canis* gDNA and cDNA. Within gDNA, highest values of RFD (553) were found in *E. caucasicum* and in one accession of *E. dens-canis* from the eastern part of the species’ range (i.e., the ‘Feleacu’ population) (Tables S1, S7). RFD tended to decrease towards the western end of species’ range, reaching values of 195 and 88 in the analyzed Spanish and French samples, respectively, while being somewhat intermediate (259) in the Croatian sample. Ideally, a future population phylogenomic study should establish whether *E. dens-canis* had an east-west spread across Southern Europe. In such a case, estimates of RFD accompanied by genome size measurements on a larger number of populations should be performed to test whether RFD between the monocot and dicot rDNA changed in the genome of *E. dens-canis* across a longitudinal geographic gradient. Such study may provide novel insights into the dynamics of evolution of rDNA in plant populations across space and time.

Within cDNA of one sample of the core *E. dens-canis* population analyzed, there appeared to be approximately 3000 times less alien ITS2 when compared to its native counterpart (Table S7). This finding – in combination with quantity estimates based on gDNA – signals that transcription efficiency of dicot rDNA in *Erythronium* is several folds lower than that of the native loci. Predicting factors that contributed to this transcription inefficiency (beyond the already mentioned pseudogenization) would be mere speculation. The ongoing Iso-Seq analysis of *E. dens-canis* RNA will potentially shed further light on the transcription efficiency of presumably horizontally transferred rDNA in plants.

### Mode of DNA Transfer Remains Elusive

An intriguing question that generally arises in the context of HGT concerns the mode of the transfer. In this study, total length of the alien DNA of *Potentilla* origin is not known. Nevertheless, the presence of stretches of alien rDNA spanning at least 85.4 kbp in case of a BAC clone and 67.8 kbp in case of Nanopore reads suggests that alien DNA may amount to hundreds of kilobases. Transfers of such pieces of DNA are not exceptional (27, 41, 44). Given the potential large size of transferred DNA, one tends to favor a direct transfer via, for example, a sap-sucking insect over a delivery via a vector organism with relatively small genome size (like bacteria or virus) into whose genome incorporation of alien DNA should have occurred first. Due to this reason a direct transfer seems to be more parsimonious. Parasitic association was shown to increase the chance of HGT (4, 39, 44–47). To our best knowledge, no insect parasite shared by *Erythronium* and *Potentilla* is known to date. It is well documented that *Erythronium* is parasitized by the rust fungus *Uromyces erythronii* (DC.) Pass. in Eurasia (48, 49). A survey of European species belonging to Rosaceae identified 39 *Potentilla* species as hosts of rust fungi (50). A surprisingly large haploid genome size of 2489 Mbp has been reported in another *Uromyces* species (51) and genome sizes are generally high in rust fungi (Pucciniales) (52). Taken together, transfer of alien DNA into the *Erythronium* genome by a rust fungus cannot be excluded though it contradicts the direct transfer hypothesis favored above.

*Erythronium* is a textbook example of myrmecochory (dispersal of elaiosome-bearing seeds by ants) and *Potentilla* is one of the two genera of Rosaceae identified so far in which myrmecochory also was reported (53). If myrmecochory somehow contributed to the HGT event, integration of alien DNA into the *Erythronium* genome could not occur prior to seed ripening. Recently, within grasses, a statistical increase of the number of HGTs was found in rhizomatous species (8). Perennial plants able to regenerate from vegetative parts (like rhizomes or bulbs) may have an increased opportunity to fix alien DNA in their germline which was previously transferred e.g., via cell-cell contact (8). *Erythronium* species are perennials and vegetative propagation of bulbs is common in the genus. Therefore, alien DNA potentially captured during e.g., germination stage could be transferred to a whole new plant via bulb formation. Such scenario also fits the weak-link model of HGT (54).

Along with direct transfers based on cell-cell contact, genetic material exchange following non-standard (illegitimate) pollination has been postulated in grasses (55). This scenario presumes growth of *Potentilla* pollen tube on *Erythronium* stigma to enable exchange of DNA between chromosomes. Such process would require certain degree of compatibility. So far, regular wide hybridization involving *Potentilla* pollen was documented with *Fragaria* (Rosaceae), leading to their commonly reported intergeneric hybridization (56). Due to the phylogenetic distance of the monocot and dicot lineages, the above scenario of HGT via pollination seems to be unlikely. The current study adds a new example to the growing list of naturally transgenic plants and essentially suggests that ‘in plants anything is possible’.

### Conclusions and Perspectives

In this article, authors invoked the horizontal gene transfer phenomenon to explain the presence of tandemly arrayed and transcriptionally active dicot-type rDNA in the nuclear genome of a monocot plant. Working with different parts (leaves, bulbs, root tips) of the host *Erythronium* plant in different laboratories refuted an artefact origin of the alien rDNA. *Potentilla*-type ITS and ETS sequences perfectly fitting a clear phylogenetic framework of the genus *Potentilla*, coupled by >150 My separate evolution of dicot and monocot lineages represent indirect but strong arguments in support of HGT scenario.

The underlying evolutionary accident between a dicot and monocot plant might had consequences for the *Erythronium* genome reaching far beyond the simple hosting of an alien-type rDNA. *Erythronium* genome will have to be carefully tiptoed in search for non-rDNA sequences of *Potentilla* origin. During this, particular attention should be paid to transposable elements and their role in the transfer, either as passive hitchhikers, or active agents, carriers, or rebuilders. Should any coding sequences be discovered, investigating their processing as well as functionality at the background of monocot genomic apparatus would be unprecedented. Ramping availability of genomic resources will provide ample opportunity to determine the extent of HGT with all of its consequences to the *Erythronium* genome.

## Materials and Methods

### Plant Materials and DNA Extraction

The list of *Erythronium* samples analyzed in the study (including geographical origin, voucher information and specific molecular analyses used) is shown in Table S1. DNA was preferentially extracted from fresh bulbs using DNeasy Plant Mini Kit (Qiagen, Hilden, Germany). Pilot molecular cytogenetic analyses confirmed specimens of the main analyzed *E. dens-canis* (‘Feleacu’) population to be diploid (2n=24), according to the common ploidy level reported for the species (57). For certain techniques where inclusion of a positive control was necessary, fresh leaves of *Potentilla reptans* L. were processed for DNA extraction.

### PCR Amplifications and Sanger Sequencing

MyTaq™ Red Mix (Bioline, London, UK) was utilized for PCR amplifications throughout the study. Primers ITS5 and ITS4 (58) were used for routine amplification of rDNA ITS. For differential amplification of *Potentilla*-specific ITS2 in *Erythronium*, new primers were designed (Table S6). The single instance of cloning of *E. dens-canis* ITS region was performed as described previously (31). *Potentilla reptans* ITS, *Erythronium* ITS clones and *Potentilla* ITS2-specific PCR products of *Erythronium* were Sanger sequenced at Macrogen Europe B.V. (Amsterdam, The Netherlands).

### Phylogenetic Analyses

*Potentilla*-type ITS and partial ETS sequences retrieved from BAC clones and Illumina datasets of *Erythronium* were added into a recent backbone phylogeny of *Potentilla* (59). This phylogeny was supplemented with additional *Potentilla* taxa having publicly available ITS sequences that were potentially closely related to the alien ITS of *Erythronium.* A maximum likelihood (ML) phylogeny was computed with the IQ-TREE (60) web server (http://iqtree.cibiv.univie.ac.at/). The strategy for public *Potentilla* ITS sequence gathering as well as details of their aligning and parameters of the applied ML phylogeny are provided in *SI Materials and Methods.* GenBank accession numbers for newly generated and previously published *Potentilla* and *Erythronium* sequences included into the study are listed in Tables S1, S8 and S9, respectively.

### Southern Blot Analysis

In addition to constructing a BAC library from flow-sorted nuclei (see further), Southern blotting technique was used to verify integration of dicot rDNA into the genome of *E. dens-canis.* Based on the BAC clone sequences of *E. dens-canis*, it was predicted the *Apo*I restriction enzyme to produce an ~878 bp long fragment selectively from the dicot-type rDNA. This comprised 3’-end of 18S rRNA gene, ITS1, 5.8S rRNA gene and ITS2. In contrast, a 2468 bp long fragment was expected to arise from the native 45S rDNA of *Erythronium* that also contained complete ITS region (Fig. 2a). Forty-five micrograms of *Erythronium* and five micrograms of *Potentilla* gDNAs (concentrated by ethanol precipitation to 0.5 μg/μl) were digested with *Apo*I-HF enzyme (New England Biolabs, Hertfordshire, UK), loaded onto 1% agarose gel, and electrophoretically separated overnight. Size-selected DNA fragments were transferred onto Amersham Hybond-N^+^ nylon membrane (GE Healthcare, Buckinghamshire, UK) by high salt transfer with 20×SSC, followed by UV-crosslinking (0.120 J/cm^2^). Amersham Alkphos Direct labeling reagents (GE Healthcare, Buckinghamshire, UK) were used for probe labeling and subsequent hybridization. The blot was probed twice. At first, gel-purified ITS region of *P. reptans* amplified from gDNA was used as probe. For the second time, *Potentilla*-specific ITS2 amplified from a dicot-type ITS clone of *E. dens-canis* which served as probe. Conditions for hybridization and post hybridization washes were the same as in Amersham protocol. Hybridization lasted overnight in a 30 cm long hybridization tube containing 30 ml hybridization buffer and 300 ng of probe DNA. Chemiluminescence detection was achieved with CDP-*Star* in a ChemiDoc MP imaging system (BioRad, Hercules, CA, USA).

### Chromosome Preparation

Mitotic chromosome spreads were prepared from root tips as previously described (61). Briefly, root tips of *E. dens-canis* and *E. sibiricum* were harvested from seedlings and bulbs of cultivated plants, respectively, pre-treated with ice-cold water for 16 h, fixed in ethanol/acetic acid (3:1) fixative for 24 h at 4°C and stored at −20°C until further use. Selected root tips were rinsed in distilled water (twice for 5 min) and citrate buffer (10 mM sodium citrate, pH 4.8; twice for 5 min), and digested in 0.3% cellulase, cytohelicase and pectolyase (all Sigma-Aldrich, St. Louis, MO, USA) in citrate buffer at 37°C for 3 h. After digestion, individual root tips were dissected on a microscope slide in 20 μl acetic acid and spread on the slide placed on a metal hot plate (50°C) for c. 30 s. Then, the preparation was fixed in freshly prepared ethanol/acetic acid (3:1) fixative by dropping the fixative around the drop of acetic acid and into it. The preparation was dried using a hair dryer and staged using a phase contrast microscope. Chromosome preparations were treated with 100 μg/ml RNase in 2× sodium saline citrate (SSC; 20× SSC: 3 M sodium chloride, 300 mM trisodium citrate, pH 7.0) for 60 min and with 0.1 mg/ml pepsin in 0.01 M HCl at 37°C for 5 min; then postfixed in 4% formaldehyde in 2× SSC for 10 min, washed in 2× SSC twice for 5 min, and dehydrated in an ethanol series (70%, 90%, and 100%, 2 min each).

### Fluorescence *in situ* Hybridization

The 821 bp ETS from *Erythronium* and 220 bp 26S rDNA from *Nicotiana* (62) were used for *in situ* localization of rDNA loci in *E. dens*-*canis* and *E. sibiricum.* The two rDNA probes were labelled with biotin-dUTP or digoxigenin-dUTP by nick translation as described in (63). 200 ng of each rDNA probe were pooled together, ethanol precipitated, dissolved in a 20 μl mixture containing 50% formamide, 10% dextran sulfate and 2× SSC, and pipetted onto microscopic slides. The slides were heated at 80°C for 2 min and incubated at 37°C overnight. Hybridized probes were visualized through fluorescently-labelled antibodies against biotin-dUTP and digoxigenin-dUTP as in (63). Chromosomes were counterstained with 4’,6-diamidino-2-phenylindole (DAPI, 2 μg/ml) in Vectashield antifade. Fluorescence signals were analyzed and photographed using a Zeiss Axioimager epifluorescence microscope and a CoolCube camera (MetaSystems, Altlussheim, Germany). Individual images were merged and processed using the Photoshop CS software (Adobe Systems, San Jose, CA, USA).

### Construction and Screening of a BAC Library from *E. dens-canis*

HMW DNA was prepared according to a protocol modified from that in ref (64). The method takes advantage of purification and sorting of nuclei using flow cytometry and embedding them into agarose plugs. Briefly, 30 g of fresh leaf tissue was fixed in 2% formaldehyde for 20 minutes and homogenized. Nuclei were concentrated from the resulting suspension by passing it through a FACSAria device (Becton Dickinson, San José, CA, USA) and were embedded in 20 μl agarose plugs (each plug harbored around 2 x 105 nuclei). The plugs were solidified on ice and incubated in lysis buffer (0.5 M EDTA pH 9.0, 1% lauroylsarcosine, 0.2 mg/ml proteinase K) at 37°C for 24 hours twice to remove residual proteins. HindIII restriction fragments of the obtained HMW DNA were size-selected by pulse field gel electrophoresis (PFGE) and ligated into pAGIBAC vector. *E. coli* DH10B-T1R strain was used as host for the cloning. Unique clones (173 610 in number and having average insert size of 115 kbp) were arranged in pools of 90 and tested. Primers for dicot and monocot ITS2 (Table S6) were used to screen the low-coverage (~0.81×) library by quantitative PCR (qPCR). Colony picking was carried out on the positive pools by a high throughput automated station QPix2 XT (Molecular Devices, San José, CA, USA). Individual BAC clones were validated by qPCR. Insert size was estimated using NotI enzymatic digestion and PFGE electrophoresis. BAC-ends sequences (BESs) were performed using Sanger sequencing. Construction and screening of the BAC library as well as its long-term storage (under the name of Ede-B-3544H) was achieved at the Plant Genomic Center (INRAE-CNRGV), Toulouse (France).

### BAC Clone Sequence Assembly and Analysis

Upon screening, multiple BAC clones tested positive for the monocot marker and a single clone (1J24) was found to contain the dicot marker. BAC clones 1J24 and monocot marker-specific 18I01 were tagged and pooled before performing sequencing on a PacBio Sequel II system with the CCS mode (chemistry 3.0). Raw reads were corrected and demultiplexed. The HiFi reads were then filtered by eliminating the residual *E. coli* and vector sequences and assembled with Canu-1.9 (1J24) and Hifiasm-0.12 (18I01). A quality control of the assembly was conducted. First, BESs obtained by the Sanger method were aligned to the ends of the assembled contigs. Second, the coverage of assemblies and the length of the obtained contigs were checked. The assemblies of BAC clones 1J24 and 18I01 had 593× and 976× coverages and 125 kbp and 90 kbp contig lengths, respectively. Each assembly resulted in one single contig. Annotation and physical map creation of obtained BAC clone sequences was performed in Geneious Prime v.2019.2.1. Searches for conserved domains within BAC clone sequences were performed at the National Center for Biotechnology Information website (https://www.ncbi.nlm.nih.gov/Structure/cdd/wrpsb.cgi). The online version of *LTR_FINDER* (65) was used to check for retrotransposon LTRs. BAC clones were further explored for transposon-like sequences with Censor (https://www.girinst.org/censor/index.php) (66) by restricting the search to *Viridiplantae.* Hits not containing simple repeats and those of at least 150 bp in length were considered.

### Test for Dicot rDNA Transcription in *Erythronium* and Assessing the Relative Fold Difference Between the Two rDNA Types

In order to test for the transcription of dicot-type rDNA in *Erythronium*, total RNA was isolated with the RNEasy Plant Mini Kit (Qiagen, Hilden, Germany) from actively growing (in October) roots of an *E. dens-canis* bulb. RNA was treated with DNase, and processed for cDNA synthesis with RevertAid First Strand cDNA Synthesis Kit (Thermo Scientific, Vilnius, Lithuania). Complementary DNA and gDNA (for control) were then used as templates in the upcoming regular PCR using primers for dicot and monocot ITS2 (Table S6). A SQ-PCR approach was employed using Maxima SYBR Green qPCR Master Mix (Thermo Scientific, Waltham, MA, USA) and primers for dicot and monocot ITS2. The experiment was extended with gDNAs of one *E. caucasicum* sample and four additional *E. dens-canis* samples from distant localities of the species’ range (Table S1). Each sample-primer pair combination was run in triplicate. Raw fluorescence data provided by a Rotor-Gene 6000 instrument (Corbett Research, Mortlake, Australia) were analyzed in the LinRegPCR v.11.0 software. Relative quantification was achieved with the RFD = *E*^-delta Cq^ formula. Aliquots of the completed SQ-PCR reactions were ultimately verified by agarose-gel electrophoresis and ethidium bromide-UV detection.

### Nanopore Long-Read Sequencing and Data Analysis

Oxford Nanopore Technologies’ (ONT) long-read sequencing service was ordered from Deep Seq (University of Nottingham, Nottingham, UK). Bulb-extracted gDNA of ~25 kbp in size was used as input for the preparation of the standard SQK-LSK109 library as per ONT’s protocol (ONT, Oxford, UK). The library was run over one PromethION Flow Cell. Base calling was completed in PromethION’s sequencing software MinKnow by using the default quality threshold of 7. Resulting 12.6 million reads (~68.2 Gigabases (Gb) of data) had N50 read length of 9.9 kbp. A total of 64998 contigs with minimum, maximum and average lengths of 1 kbp, 1.38 Mbp and 5.69 kbp, respectively were assembled using CANU v 1.8 (67) from the raw Nanopore data. Polishing of the CANU assembly was attempted using Pilon (68) (https://github.com/broadinstitute/pilon) with default parameters, using a preprocessed paired-end Illumina dataset (see below). Both the raw reads and the contigs were screened for dicot/monocot ITS1/ITS2 fragments (Table S3) in Geneious Prime. In terms of the number and the length of Nanopore reads containing *Potentilla*-specific rDNA sequences, the assembly did not turn out to be superior to the original read dataset (Table S4). Raw Nanopore sequences that were analyzed and presented in the current study were included in the supplementary dataset (Datasets S1 and S2).

### Illumina Whole Genome Shotgun (WGS) Sequencing and Read Analysis

A TruSeq shotgun library (of 350 bp insert-size) was prepared from gDNA of *E. dens-canis* and paired-end (150 bp) sequenced via a HiSeqX platform at Macrogen B.V. Fifty gigabases of raw reads were preprocessed to remove low-quality reads and controls and to trim adapter sequences by using a combination of tools provided in the BBMap project (69, 70). A *de novo* assembly of Illumina reads was performed in SPAdes (71). *Potentilla*-specific ITS and partial-ETS sequences were extracted from contigs following a screen in Geneious Prime with dicot ITS1/ITS2 sequences.

In case of *E. caucasicum* 5 Gb of WGS data was obtained. A shotgun gDNA library of 350 bp insert size was prepared and paired-end (150 bp) sequenced at Novogene (Cambridge, UK) using the Novaseq6000 platform. Raw reads were imported into Geneious Prime, paired, and end-trimmed using the error probability limit of 0.05. Paired reads were then mapped to dicot ITS1/ITS2 reference sequences. Mapped reads were assembled into contigs and consensuses of the contigs were generated by using the built-in Geneious assembler.

## Acknowledgments

The authors would like to thank Adela Jurković (University of Zagreb) for providing guidance to LB in regard with the Southern blotting technique. Chemiluminescence detection was achieved with the assistance of László-Csaba Bencze and Alina Filip (Babeş-Bolyai University) whose help is also acknowledged. The authors thank Kunigunda Macalik, Jānis Rukšāns and the Rare Bulb Nursery (Latvia), Andrey Dedov, Sergey Banketov, Julianna Lingvay, Paul-Marian Szatmari, Gábor Sramkó, Boštjan Surina, Polina A. Volkova, Nikolay V. Stepanov, Alexander L. Ivanov for their help in the acquisition of plant material. This work was supported by a grant of the Romanian Ministry of Research and Innovation, CNCS - UEFISCDI, project number PN-III-P1-1.1-PD-2016-0919, within PNCDI III. PAB was supported by the research grant 20-12496X (Grant Agency of the Czech Republic). The authors would like to acknowledge support of the GENTYANE platform of Clermont-Ferrand INRAE Center (http://gentyane.clermont.inra.fr/) for providing assistance in NGS sequencing and of the Genotoul bioinformatics platform Toulouse Midi-Pyrenees (Bioinfo Genotoul, doi: 10.15454/1.5572369328961167E12, http://bioinfo.genotoul.fr) for providing computing resources.

