## Supplementary Information for "Intact rDNA arrays of *Potentilla*-origin detected in *Erythronium* nucleus suggest recent eudicot-to-monocot horizontal transfer"

**This file includes:**

Supplementary text

Figures S1 to S7

Tables S1 to S9

Legend for Dataset S1 to S2

SI References

**Other supplementary materials for this manuscript include the following:**

Dataset S1. Raw sequence of Nanopore read 202d7c58-6078-4612-8e8a-bf45fa7b7b04 containing parts of both native and *Potentilla*-type rDNA.

Dataset S2. Raw sequence of the longest Nanopore read (e16c7cdb-900b-4c49-8019-e4f837b9606f) containing rDNA of *Potentilla*-origin.

SI Materials and Methods

**Acquisition of Publicly Available Sequences, Sequence Alignment and Phylogenetic Analyses.**

Selected *Potentilla* ITS and partial ETS sequences were downloaded from GenBank in order to reconstruct one of the most recent backbone phylogenies of the genus (1). Sequences of one representative of each of the *Potentilla*-related genera *Alchemilla, Chamaecallis, Chamaerhodos, Dasiphora, Drymocallis, Fragaria, Sibbaldia, and Sibbaldianthe* were also downloaded in order to increases the robustness of the subsequent phylogenetic analysis.

Newly generated *Potentilla*-specific sequences of *Erythronium* used for phylogeny reconstructions included two conventional ITS clones, ITS and partial ETS sequences extracted from BAC clone 1J24 as well as *DE NOVO* and reference-guided Illumina contig consensus sequences of ITS and partial ETS. These were embedded into the backbone phylogeny of *Potentilla* as an effort to establish their phylogenetic placement.

ITS and partial ETS sequences were concatenated and aligned in MEGA v.7 (2) using Muscle. A ***first pilot*** maximum likelihood (ML) phylogenetic analysis conducted in raxmlGUI v.2.0 (3) confirmed the recovery of the main clades of *Potentilla* (Fragarioides, Alba, Reptans, Ivesia and Argentea) as well as nesting of the *Potentilla*-type *Erythronium* sequences in the Argentea clade. As a next step, GenBank was searched for *Potentilla* ITS sequences with the term “Potentilla[Organism] AND 5.8S” in August 9, 2021. The recovered 647 sequences were pooled with the *Potentilla*-type ITS sequences of *Erythronium* and were aligned with the MAFFT online alignment tool v.7 (<https://mafft.cbrc.jp/alignment/server/>). The alignment was visually checked and submitted for a ***second pilot*** phylogenetic analysis using the IQ-TREE (4) web server (<http://iqtree.cibiv.univie.ac.at/>) with default settings. Based on this phylogeny, a large, well-supported clade (BS=87%) was identified in which *Potentilla*-type ITS sequences of *Erythronium* were deeply nested. Based on this clade, a number of 55 taxa were identified as potentially closely related to the *Erythronium* sequences in addition to a few taxa already figuring in the backbone phylogeny. The newly selected ITS sequences (1 sequence/taxon) were inserted into the alignment of the *Potentilla* backbone phylogeny. In 39 out of the 55 cases, ETS sequences were also available from the same accessions (voucher) from which ITS sequences were generated. These were also embedded into the final alignment. Nucleotides represented by *IUPAC* codes were changed to missing data in the alignment. As a conservative measure, an unambiguously aligned 19 bp long portion of the alignment from within ITS2 was deleted. The resulting final alignment, having 103 sequences with 1088 columns, 314 parsimony-informative, 173 singleton and 601 constant sites, was submitted for the ***ultimate*** ML analysis in the IQ-TREE web server. ModelFinder (5) selected the TNe+I+G4 and TIM3e+G4 evolutionary models as the ones best fitting the ITS and ETS datasets, respectively. Statistical robustness of branches was tested with ultrafast bootstrapping (UFBoot) (6) implying 1000 replicates.

Native ITS sequence of *E. dens-canis* from BAC clone 18I01 was inserted into a previously published phylogeny of Eurasian *Erythronium* (7) for general control. The respective phylogenetic analysis relied on the ML criterion implemented in raxmlGUI, with 1000 bootstrap replicates, under the GTR + G model of nucleotide evolution.

Translated potentially coding sequences (CDS’) of BAC clones 1J24 and 18I01 were highly divergent and, apart from their DUF4283 domain parts, could not be unambiguously aligned. Therefore, the 146 amino acid-long DUF4283 sequence of BAC clone 18I01 was blast-searched with the blastp program against the non-redundant protein database. Similar sequences were selected for phylogeny purposes in a way to represent all the taxa among the first 100 blastp hits. BAC clone and publicly downloaded DUF4283-like sequences were aligned in MEGA and subjected to a Neighbor-Joining analysis. Statistical robustness of tree branches was assessed with the Bootstrap test using 1000 replicates.

All resulting phylogenetic trees were edited in FigTree v.1.4.2 (8) and the Inkscape v.0.92.4 software ([www.inkscape.org](http://www.inkscape.org)).

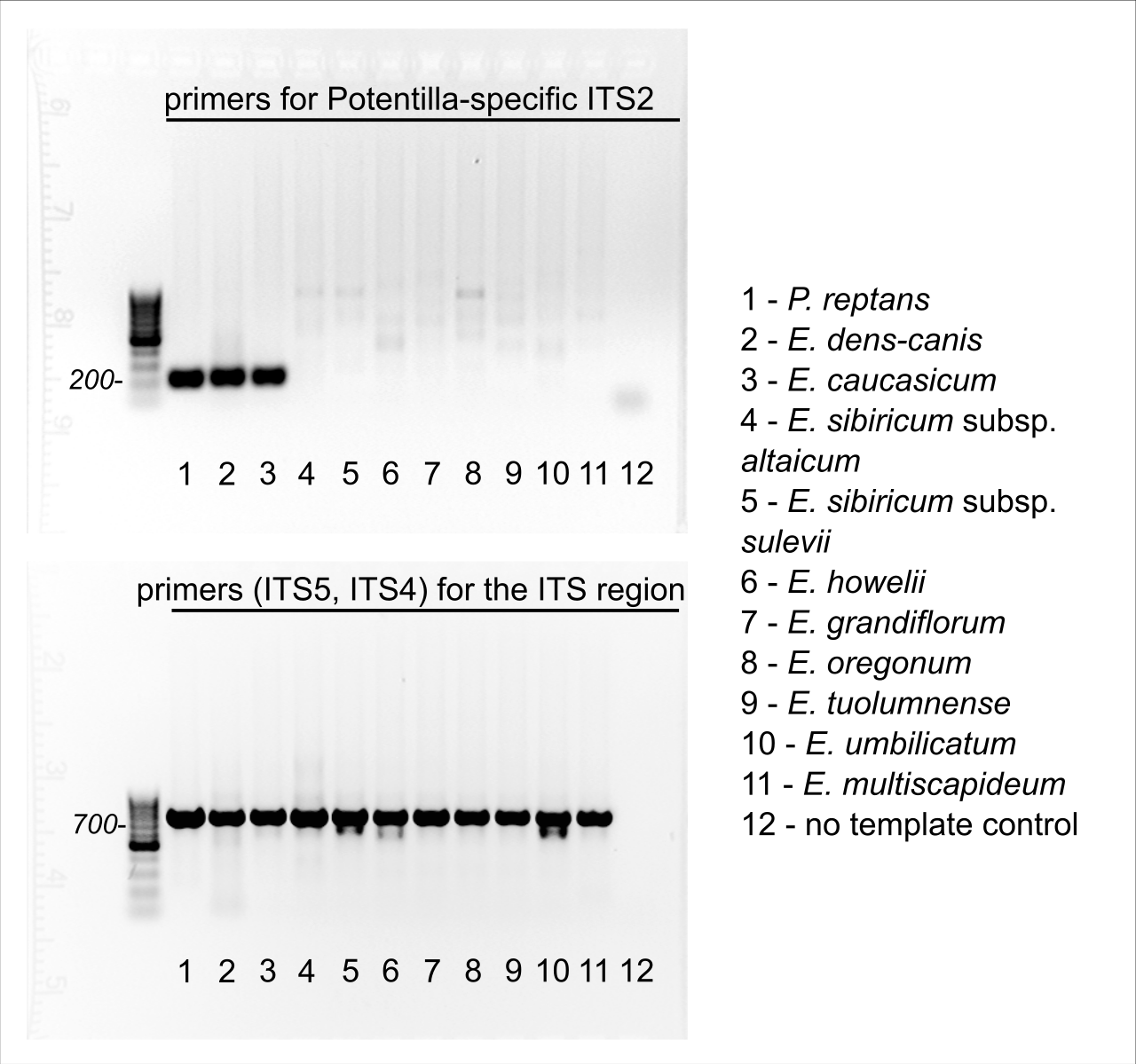

Fig S1. Test for the presence of *Potentilla*-specific ITS2 in the genomes of selected New- and Old World *Erythronium* taxa. PCR products were visualized upon ethidium bromide-UV detection and were based on the primers and gDNA templates shown in the figure body. Amplification of the whole ITS region served as general control.

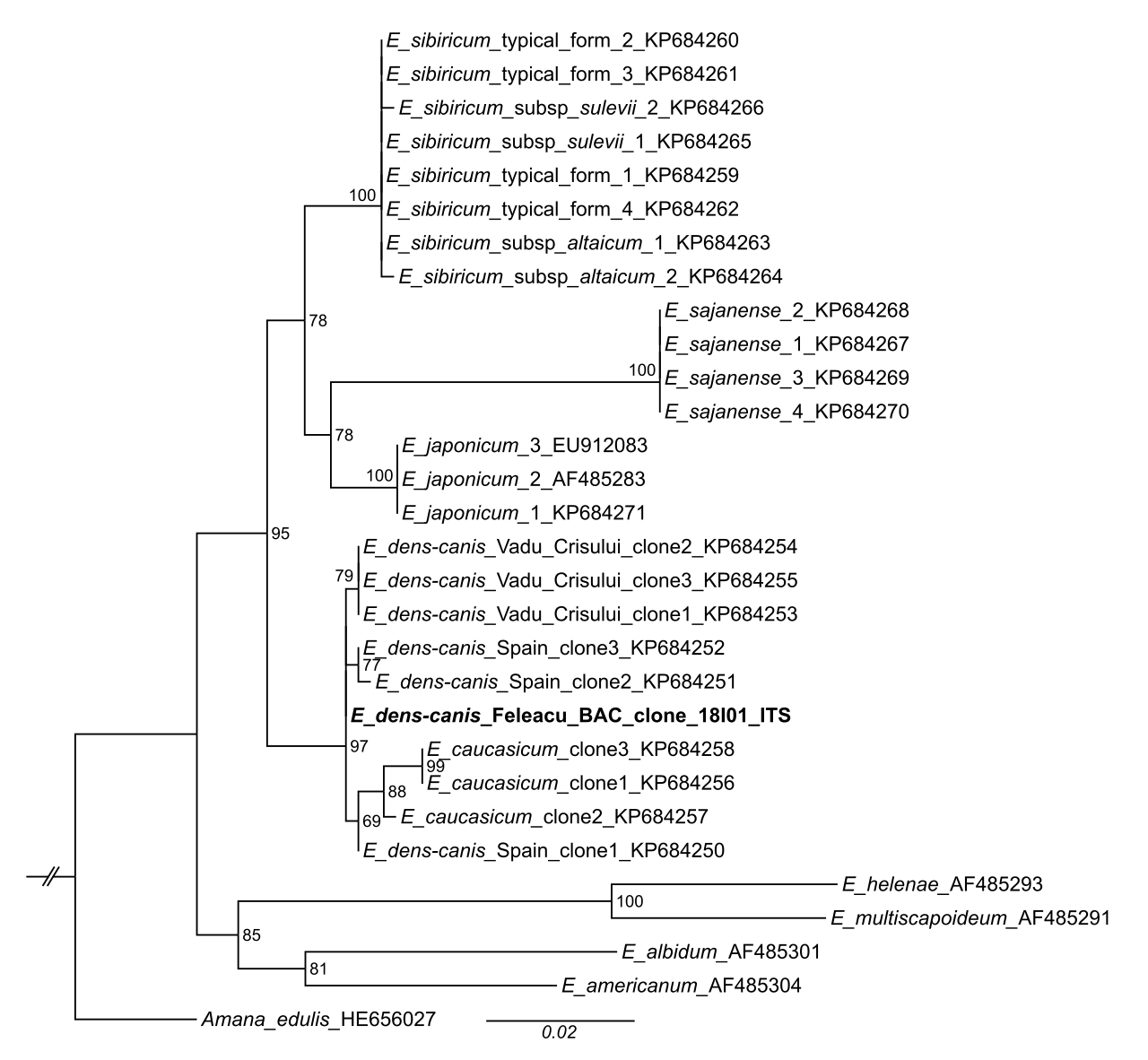

Fig S2. RAxML best ML phylogram of publicly downloaded and newly generated (in bold) *Erythronium* ITS sequences. Nodal support values were derived from 1000 rapid bootstrap replicates. ITS sequence of BAC clone 18I01 represents the common sequence type of the 12 repeat units.

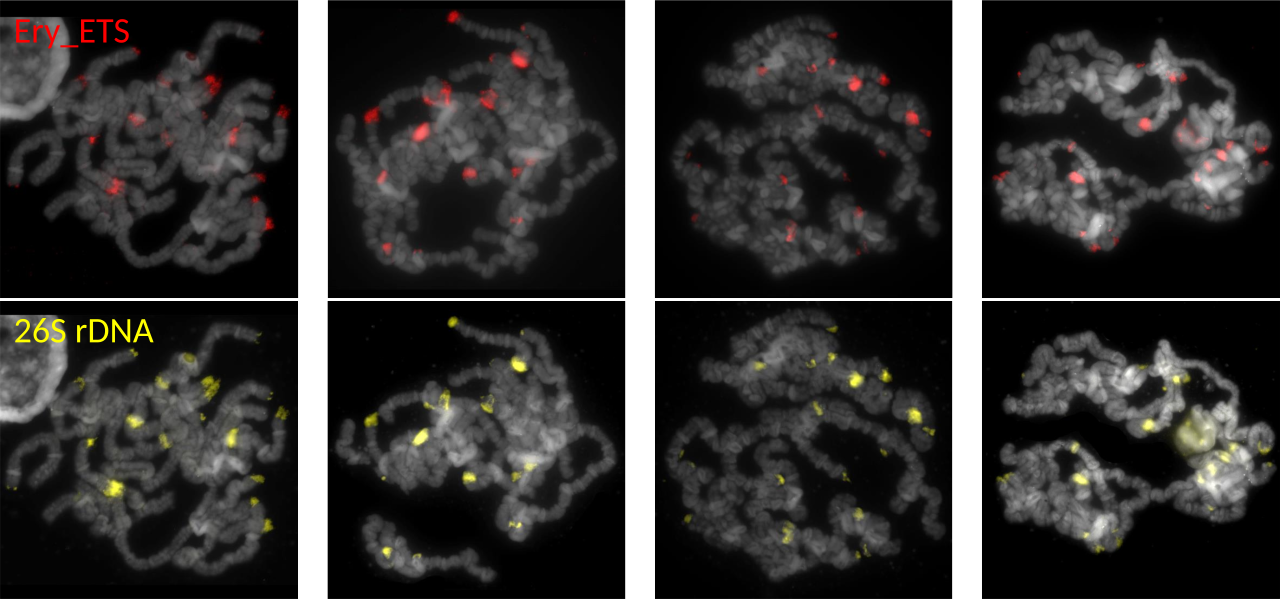

Fig S3. Chromosome localization of rDNA in *Erythronium sibiricum*. Four mitotic chromosome complements were hybridized with *Erythronium* ETS (Ery_ETS, red signals) and *Nicotiana* 26S rDNA probe (yellow signals). Chromosomes were counterstained with DAPI.

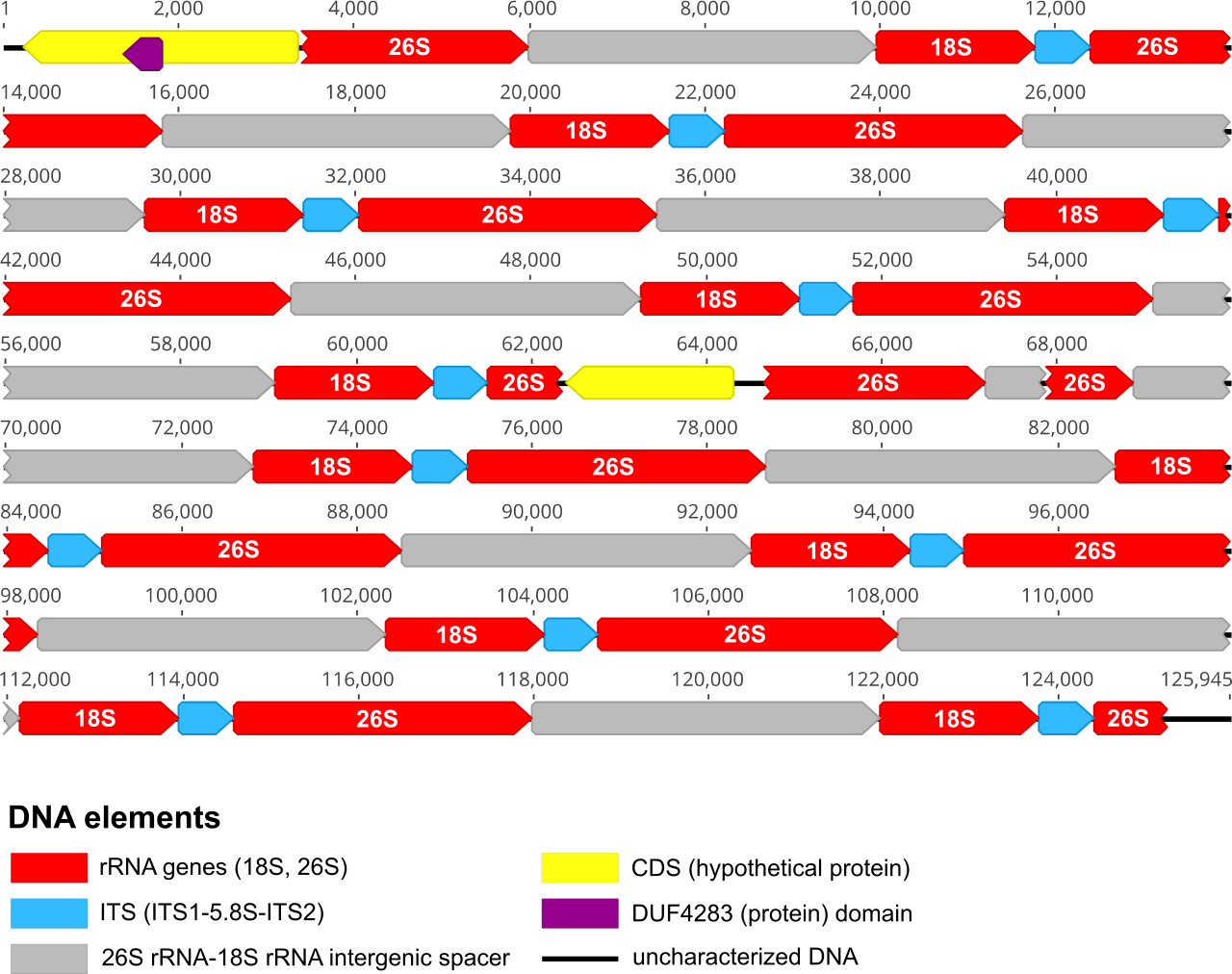

Fig S4. Physical map of *E. dens-canis* BAC clone 18I01 containing native rDNA.

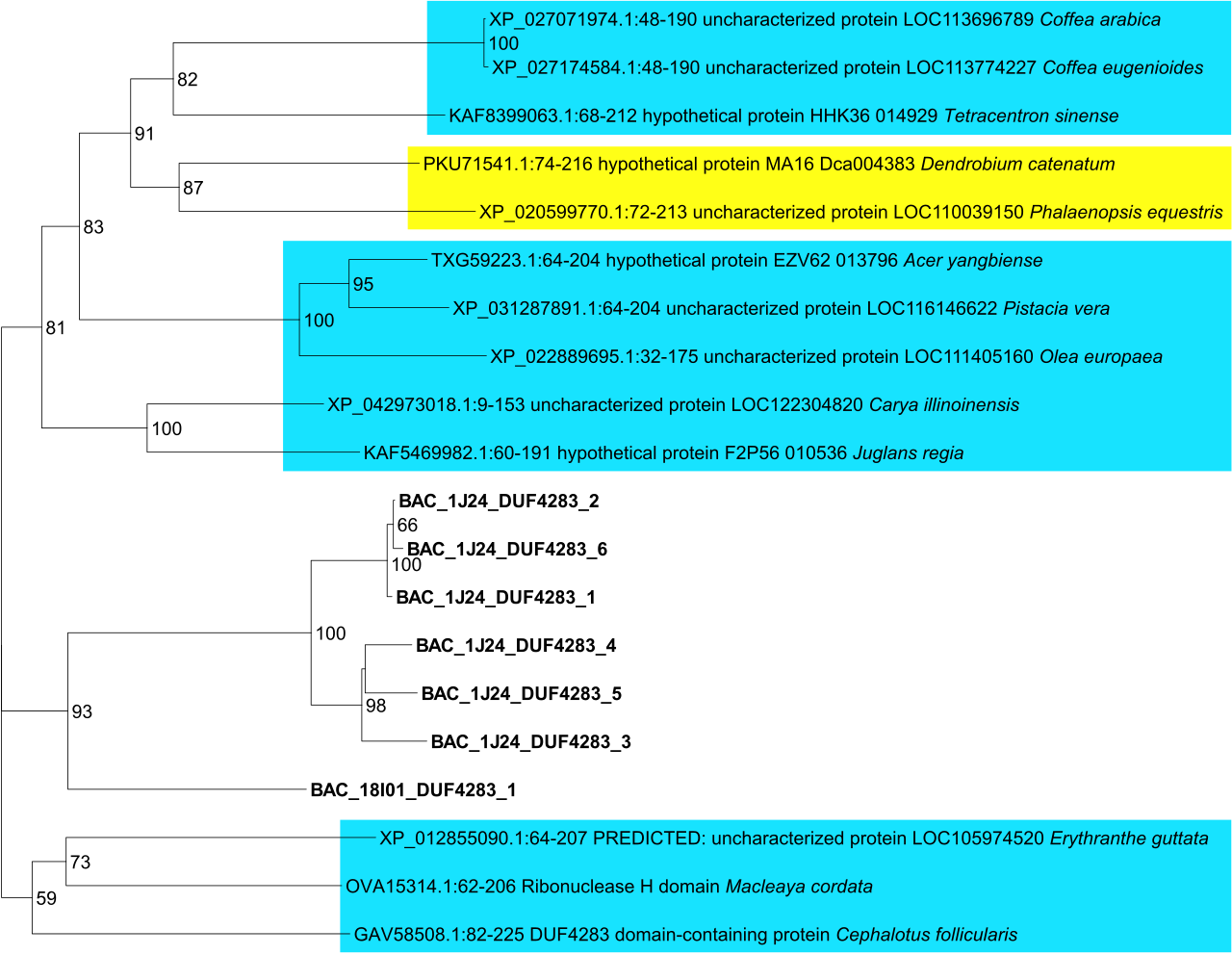

Fig S5. Neighbor-Joining phylogeny of translated DUF4283 domain sequences of BAC clones 1J24 and 18I01 of *E. dens-canis* supplemented with publicly available similar protein fragments. Numbers (1-6) from within BAC clone 1J24 DUF4283 sequence names refer to their order in the physical map of the clone. Eudicot and monocot taxa are shaded in blue and yellow, respectively.

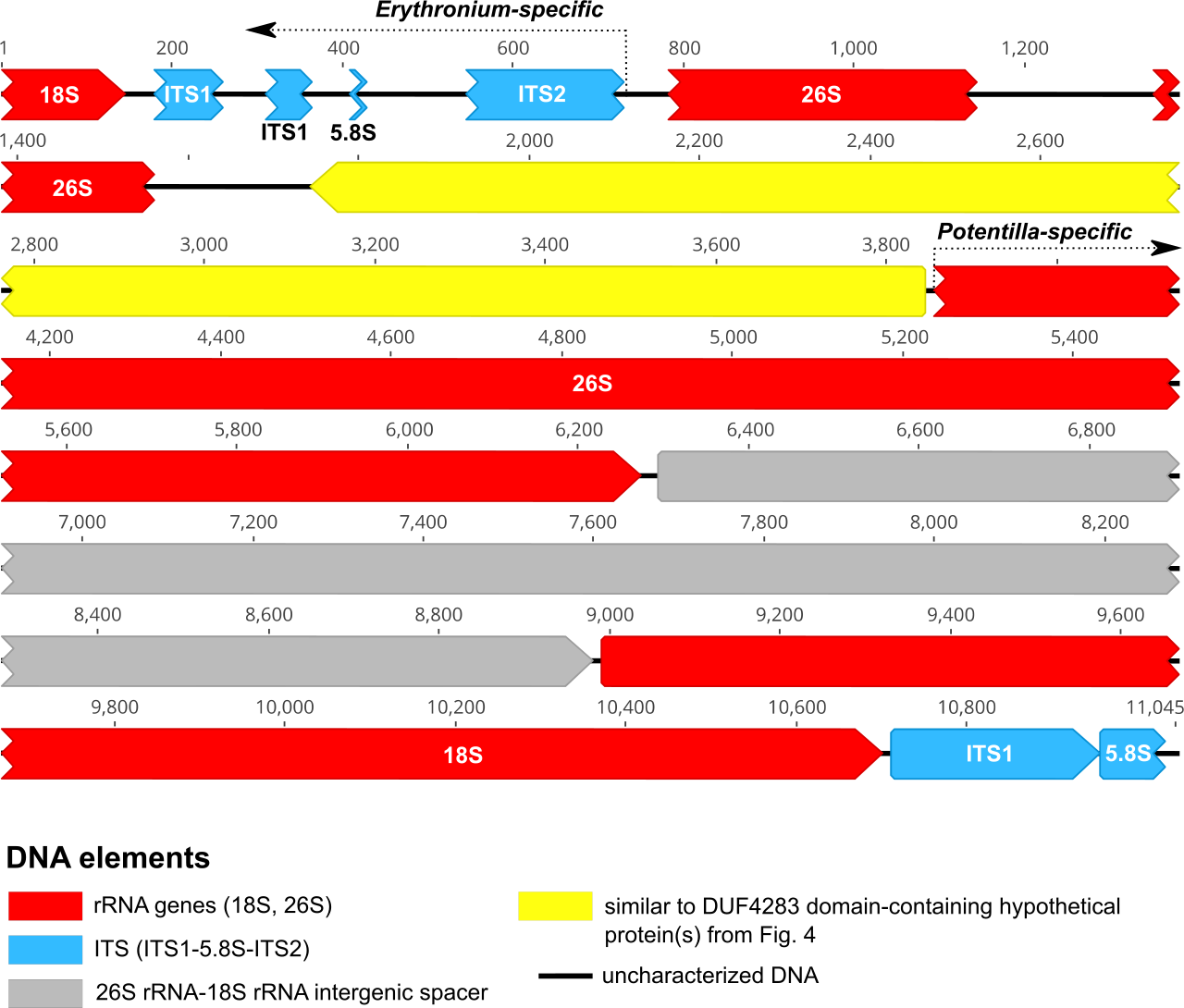

Fig S6. Schematic representation of the chimeric *E. dens-canis* WGS Nanopore read “202d7c58-6078-4612-8e8a-bf45fa7b7b04” containing native and *Potentilla*-specific rDNA fragments.

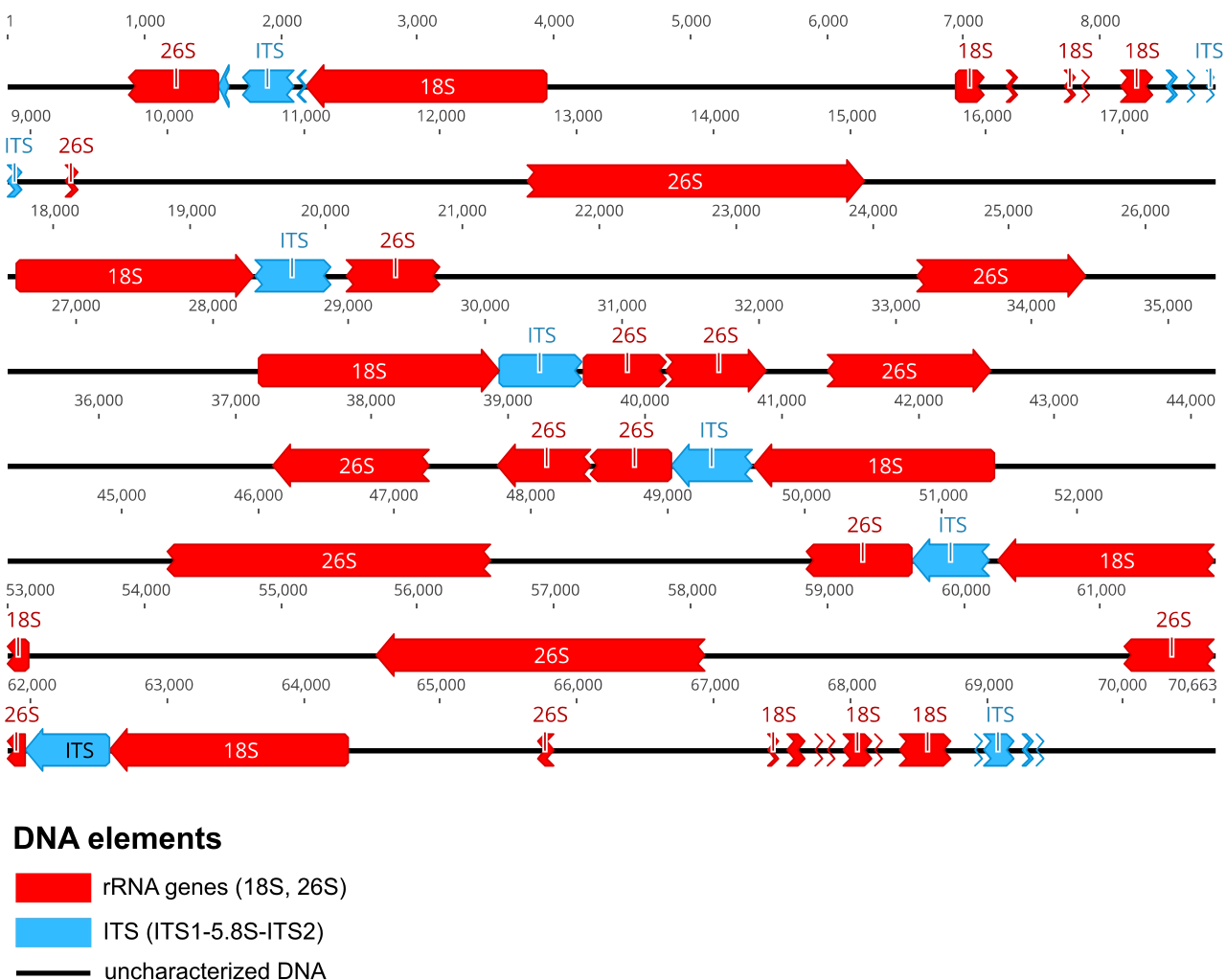

Fig S7. Organization of the main 45S rDNA units (18S, 26S, ITS) within the longest Nanopore WGS read of *E. dens-canis* containing rDNA of exclusively *Potentilla*-origin.

Table S1. List of samples used in the study including their geographic origin, voucher information, GenBank accession numbers (of newly generated sequences), type of plant material utilized and the experiments/techniques these were involved in.

| **Samples** | **Geographic origin** | **Voucher specimen or sample collector / supplier** | **GenBank accession numbers** | **Biological material** | **Genetic material** | **Experiments** | | | | | | | | |
| --- | --- | --- | --- | --- | --- | --- | --- | --- | --- | --- | --- | --- | --- | --- |
|  |  |  |  |  |  | **Cloning and/or Sanger sequencing of *Potentilla*-specific ITS/ITS2** | **Illumina WGS (50 Gb)** | **Illumina WGS (5 Gb)** | **Nanopore WGS** | **BAC library** | **SQ-PCR (based on *Potentilla/ Erythronium* ITS2)** | **Southern blot** | **FISH** | **PCR screen for *Potentilla*-specific ITS2** |
| *Erythronium dens-canis* L. | Făgetul Clujului Forest, Feleacu, Cluj County, Romania,  N 46.708730 E 23.586949 | CL 668976 | MW357751 (BAC clone 1J24)  MW357750 (BAC clone 18I01)  MW400604 (*Potentilla*-specific ITS) ^‡^  MW400603 (*Potentilla*-specific ETS) ^‡^ | bulb | DNA |  | **✓** |  | **✓** |  | **✓** | **✓** |  | **✓** |
|  |  |  |  | young roots | RNA, cDNA |  |  |  |  |  | **✓** |  |  | **✓** |
|  |  |  |  | root tips of seedlings (germinated in culture) |  |  |  |  |  |  |  |  | **✓** |  |
|  |  |  |  | fresh leaves | DNA (from nuclei) |  |  |  |  | **✓** |  |  |  |  |
| *E. dens-canis* ^*^ | Vadu Crișului, Bihor County, Romania | CL 664526 | MW400605, MW400606 (*Potentilla*-specific ITS clones) | dried leaves | DNA | **✓** |  |  |  |  |  |  |  |  |
| *Erythronium caucasicum* Woronow | Stavropol, Stavropol Krai, Russia | Sergey Banketov | MW400602 (*Potentilla*-specific ITS) ^§^  MW400601 (*Potentilla*-specific ETS) ^§^ | bulb | DNA |  |  | **✓** |  |  |  |  |  | **✓** |
| *E. caucasicum* ^†^ | Stavropol, Stavropol Krai, Russia | CL 664522 | MW930739 (*Potentilla*-specific ITS2) | dried leaves | DNA | **✓** |  |  |  |  | **✓** |  |  | **✓** |
| *E. dens-canis* ^†^ | Kladnik, Drugovo Municipality, North Macedonia | NHMR 2176 | MW930742 (*Potentilla*-specific ITS2) | dried leaves | DNA | **✓** |  |  |  |  | **✓** |  |  | **✓** |
| *E. dens-canis* ^†^ | Soboli, Primorje-Gorski Kotar County, Croatia | NHMR 2054 | MW930740 (*Potentilla*-specific ITS2) | dried leaves | DNA | **✓** |  |  |  |  | **✓** |  |  | **✓** |
| *E. dens-canis* ^†^ | Santiago de Compostela, A Coruña Province, Spain | SANT 68296 | MW930741 (*Potentilla*-specific ITS2) | dried leaves | DNA | **✓** |  |  |  |  | **✓** |  |  | **✓** |
| *E. dens-canis* ^†^ | Chemillieu, Savoie Department, France | CL 664533 | MW930743 (*Potentilla*-specific ITS2) | dried leaves | DNA | **✓** |  |  |  |  | **✓** |  |  | **✓** |
| *Erythronium sibiricum* subsp. *altaicum* Rukšāns | Zmeinogorsk, Altai Krai, Russia | Andrey Dedov |  | bulb | DNA |  |  |  |  |  |  | **✓** |  | **✓** |
|  |  |  |  | root tips of bulbs |  |  |  |  |  |  |  |  | **✓** |  |
| *Erythronium sibiricum* subsp. *sulevii* Rukšāns | Altayskiy village, near the border with the Altay Republic, locus classicus, Russia | Janis Rukšāns, www.rarebulbs.lv |  | bulb | DNA |  |  |  |  |  |  |  |  | **✓** |
| *Erythronium umbilicatum* C.R.Parks & Hardin | unknown | Janis Rukšāns, www.rarebulbs.lv |  | bulb | DNA |  |  |  |  |  |  |  |  | **✓** |
| *Erythronium howelii* S.Watson | Siskiyou Mts., Josephine County, OR, USA | Janis Rukšāns, www.rarebulbs.lv |  | bulb | DNA |  |  |  |  |  |  |  |  | **✓** |
| *Erythronium grandiflorum* Pursh | Wenatchee Mts., Kittitas County, WA, USA | Janis Rukšāns, www.rarebulbs.lv |  | bulb | DNA |  |  |  |  |  |  |  |  | **✓** |
| *Erythronium multiscapideum* (Kellogg) A.Nelson & P.B.Kenn. | Klamath Mts., Trinity County, CA, USA | Janis Rukšāns, www.rarebulbs.lv |  | bulb | DNA |  |  |  |  |  |  |  |  | **✓** |
| *Erythronium oregonum* Applegate | Siskiyou Mts., Josephine County, OR, USA | Janis Rukšāns, www.rarebulbs.lv |  | bulb | DNA |  |  |  |  |  |  |  |  | **✓** |
| *Erythronium tuolumnense* Applegate | Tuolumne County, NE of Columbia, above S Fork Stanislaus River, CA, USA | Janis Rukšāns, www.rarebulbs.lv |  | bulb | DNA |  |  |  |  |  |  |  |  | **✓** |
| *Potentilla reptans* L. | Cluj-Napoca, Romania, N 46.786362 E 23.596416 | CL 668975 | MW357749 (ITS) | dried leaves |  | **✓** |  |  |  |  |  | **✓** |  | **✓** |

^*^ Sample included in previous phylogenetic study of Eurasian *Erythronium* (7).

^†^ Sample included in previous phylogeographic study of *E. dens-canis* (9).

^‡^ *DE NOVO* Illumina contig consensus sequence.

^§^ reference-guided Illumina contig consensus sequence.

Table S2. Transposable element-like fragments in BAC clone 1J24 of *E. dens-canis* as identified by CENSOR.

| **Nucleotide range** | **Fragment length (bp)** | **Name of similar TE element from Repbase** | **Species of origin (of similar TE element)** | **Class of TE element (as specified by the CENSOR output)** | **Repbase Reports version** | **Reference** |
| --- | --- | --- | --- | --- | --- | --- |
| 5 - 250 | 246 | EnSpm-1_FV | *Fragaria vesca* | DNA/EnSpm/CACTA | 2013, 13(2) | (10) |
| 1,232 - 1,762 | 531 | EnSpm-1N1_FV | *Fragaria vesca* | DNA/EnSpm/CACTA | 2013, 13(2) | (10) |

Table S3. Nucleotide sequences of native and *Potentilla*-specific ITS1/ITS2 fragments of *E. dens-canis* that served as guides in mapping of the Nanopore and Illumina reads.

| *Erythronium* ITS1 (229 bp) | TACCCGACCGAAAGACTGTGAACTTGTAAACGGATGTCACCAGGGTTGTCGGGCAAGCTCGACCTCCCTGGACCCCCGTCGCCC CCTTTCGGAGCGACCTTGTGTTGCACGGATGGGGGTGGTACGGGATAACGAAACCCCGGCGCTGCATGCGCCAAGGAACATATG TGACCGGATGGACGTGCGCCTCTGCCCCTGTGGTGAGGCAACGACCGCTGAACATTATCAT |
| --- | --- |
| *Potentilla*-type ITS1 (229 bp) | AGAACGACCCGAGAACGTGTTTCAACGCTTGGAGACGGGGGGCCTCGCGGCTCCTCGCCTCCTCATCCCGGGAAGCGAAGCCT CGCGTGTCGTGCTTCGGCGCTTCCCGCTTGGCCGACCTCTCCGGGCGTACTGAACATCGGCGTGAATTGCGCCAAGGAACTTGA ATGAAGGAGCGTCACCCCGCCGTCCCCGGAGACGGTGACTGCGCGGGTGGTTCGTCGTCTTC |
| *Erythronium* ITS2 (205 bp) | GCCTCGCGTCGCTCTATGCTCCTGACCCTTCGGGGCGGTGGTGTTGATGCGGAAATTGGCCCCCCGTGCCTTGTGTGCGGTGG GCTAAAGAGAGGGCTGCCAGCCAGGTGGGGCACGGCAAGTGGTGGACATAGCGCCAGCAGGATGCCGTGGCCCCCCTGGGTG GAAGGACCCAAGTGACCCGGATAAGGTGACTCGCACTCCT |
| *Potentilla*-type ITS2 (205 bp) | ACGTCGTTGCCCCTCCCAACCCCTCTGGGAGTTGGGTGGGACGGATGATGGCCTCCCGTGCGCTCCGTCGCGCGGTTGGCATA AATAACAAGTCCTCGGCGACCCAGCGCCGCGACAATCGGTGGTTGTCAAACCTCGGTGTCCCGTCGCGCGCGCGTCGTCCCGG GCTTTTCCCATCTAATGCGCGTCGCTTCGTCGGCGCTTT |

Table S4. Summary statistics of the Nanopore dataset including the result of its screening with the reference sequences from Table S3.

| **Dataset type** | **# reads / contigs** | **Mean read / contig length** | **N50 read / contig length** | **Length of the longest read / contig** | **# reads / contigs mapping to the reference sequences below:** | | | **Length of the longest mapped sequence** |
| --- | --- | --- | --- | --- | --- | --- | --- | --- |
| Raw Nanopore dataset | 12,669,040 (reads) | 5,386 | 9,909 | 151,515 | Potentilla ITS1 (229 bp) | 230 | 70,664 | |
|  |  |  |  |  | Potentilla ITS2 (205 bp) | 192 | 70,664 | |
|  |  |  |  |  | Erythronium ITS1 (229 bp) | 3,480 | 79,543 | |
|  |  |  |  |  | Erythronium ITS2 (205 bp) | 5031 | 78,569 | |
|  |  |  |  |  | Pot. ITS1 & Ery. ITS1 | 0 | – | |
|  |  |  |  |  | Pot. ITS1 & Ery. ITS2 | 1 | 11,046 | |
|  |  |  |  |  | Pot. ITS2 & Ery. ITS1 | 0 | – | |
|  |  |  |  |  | Pot. ITS2 & Ery. ITS2 | 0 | – | |
| Pilon-corrected CANU assembly of Nanopore reads | 64,998 (contigs) | 5,695 | 12,299 | 1,385,370 | Potentilla ITS1 (229 bp) | 4 | 20,860 | |
|  |  |  |  |  | Potentilla ITS2 (205 bp) | 4 | 20,860 | |
|  |  |  |  |  | Erythronium ITS1 (229 bp) | 4 | 143,869 | |
|  |  |  |  |  | Erythronium ITS2 (205 bp) | 4 | 143,869 | |
|  |  |  |  |  | Pot. ITS1 & Ery. ITS1 | 0 | – | |
|  |  |  |  |  | Pot. ITS1 & Ery. ITS2 | 0 | – | |
|  |  |  |  |  | Pot. ITS2 & Ery. ITS1 | 0 | – | |
|  |  |  |  |  | Pot. ITS2 & Ery. ITS2 | 0 | – | |

Table S5. Numbers of polymorphic nucleotide sites in native and alien 18S and 26S rRNA genes within BAC clones 18I01 and 1J24 of *E. dens-canis*.

| **Type of gene** | **# of sequences in the alignment** | **# of polymorphic nucleotide sites** |
| --- | --- | --- |
| Native 18S | 12 complete | 2 (/1813) |
| Alien 18S | 8 complete | 34 (/1807) |
| Native 26S | 10 complete, 2 incomplete | 7 (/3407) |
| Alien 26S | 2 complete, 6 partial 5’-end sequences | 53 (/3349) |
| Alien 26S | 2 complete, 6 partial 3’-end sequences | 82 (/3349) |

Table S6. List of the PCR primers used in the study.

| **Primer name** | **Target region** | **Sequence (5′–3′)** | **Product size (bp, *not* including primers)** | **Annealing temperature (°C)** | **Application** | **Reference** |
| --- | --- | --- | --- | --- | --- | --- |
| ITS5 | ITS1-5.8S-ITS2 (*Erythronium* and/or *Potentilla*-type) | GGAAGTAAAAGTCGTAACAAGG | 694-695 | 53 | Standard PCR amplifications | (11) |
| ITS4 |  | TCCTCCGCTTATTGATATGC | 694-695 | 53 |  | (11) |
| Ery_ITS2_F2 | *Erythronium* ITS2 | CTCGCGTCGCTCTATGCT | 187 | 57 | BAC library and gDNA screening, SQ-PCR | This study. |
| Ery_ITS2_R1 |  | CGAGGCGACGACACAATCCCTT | 187 | 57 |  | This study. |
| Pot_ITS2_F1 | *Potentilla*-type ITS2 | CGTCACACGTCGTAGCCC | 175 | 57 | BAC library and gDNA screening,  SQ-PCR | This study. |
| Pot_ITS2_R2 |  | AAAGCGCCGACGAATCGA | 175 | 57 |  | This study. |
| Pot_ITS2_F1_v2^*^ | *Potentilla*-type ITS2 | CGTCACACGTCGTTGCCC | – | – | sequencing | This study. |
| Pot_ITS2_R2_v2^†^ |  | AAAGCGCCGACGAAGCGA | – | – | sequencing | This study. |
| Ery_ETS_F3 | *Erythronium* ETS | GTTGGCTGTTGGATGTGCTT | 821 | 59 | FISH probe amplification | This study. |
| Ery_ETS_R3 |  | CGCATGAAAACCGTCCTAGT | 821 | 59 | FISH probe amplification | This study. |
| P1 | *Nicotiana* 26S | GAATTCACCCAAGTGTTGGGAT | 220 | 55 | FISH probe amplification | (12) |
| – |  | AGAGGCGTTCAGTCATAATC | 220 | 55 | FISH probe amplification | (12) |

^*^ Improved (corrected) version of Pot_ITS2_F1

^†^ Improved (corrected) version of Pot_ITS2_R2

Table S7. Relative fold difference (RFD) between native and alien ITS2 in *Erythronium* gDNA/cDNA based on a SQ-PCR approach.

| **Species** | **Country origin of sample** | **Type of template (gDNA / cDNA)** | **Monocot ITS2 / dicot ITS2 RFD** |
| --- | --- | --- | --- |
| *E. dens-canis* | Romania | gDNA | 553 |
| *E. dens-canis* | Romania | cDNA | 2958 |
| *E. caucasicum* | Russia (Caucasus) | gDNA | 553 |
| *E. dens-canis* | North Macedonia | gDNA | 541 |
| *E. dens-canis* | Croatia | gDNA | 259 |
| *E. dens-canis* | Spain | gDNA | 195 |
| *E. dens-canis* | France | gDNA | 88 |

Table S8. GenBank accession numbers and voucher information of publicly downloaded ITS and ETS sequences of the *Potentilla* phylogeny.

| **Taxa** | **Voucher specimen** | **ITS** | **ETS** |
| --- | --- | --- | --- |
| *Alchemilla cryptantha* Steud. ex A. Rich | T. Eriksson 914 (S) | FJ356153 | FJ422344 |
| *Argentina lignosa* (Willd. in D.F.K. Schltdl.) Soják | M. Töpel MA132 (GB) | FJ356171 | FJ422369 |
| *Chamaecallis perpusilloides* (W.W. Sm.) Smedmark | Feng 52 (HIB) | KP875287 | KP875280 |
| *Chamaerhodos mongholica* Bunge | E. Rosenius 1028 (S) | FJ356155 | FJ422349 |
| *Dasiphora glabra* (G. Lodd.) Soják | Feng 120 (HIB) | KP875289 | KP875277 |
| *Drymocallis rupestris* (L.) Soják | M. Lundberg 6 (S) | FJ356163 | FJ422359 |
| *Fragaria viridis* Weston | M. Lundber 16 (S) | FJ356166 | FJ422364 |
| *Horkelia bolanderi* A. Gray | Eriksson s.n. (SBT) | FN430789 | FN421401 |
| *Ivesia kingii* S. Watson | J. L. Reveal et al. #4782 (GB) | FN430787 | FN421377 |
| *Potentilla adscharica* Sommier & Levier ex R.Keller | E:00409739 | KT985664 | KT985793 |
| *Potentilla alba* L. | MA 122 (GB) | FN430774 | FN421355 |
| *Potentilla alchemilloides* Lapeyr. | A. & A.-L. Anderberg 26 (S) | FJ356168 | FJ422367 |
| *Potentilla anatolica* Peșmen | E:00409724 | KT985669 | KT985798 |
| *Potentilla arctica* Lehm. | ARC4 | MW042148 | - |
| *Potentilla argaea* Boiss. | E:00409678 | KT985673 | KT985802 |
| *Potentilla argentea* L. | MA 143 (GB) | FN430808 | FN421387 |
| *Potentilla astragalifolia* Bunge | RSA:307559 | KT985676 | KT985805 |
| *Potentilla aucheriana* Th.Wolf ex Bornm. | E:00409752 | KT985677 | KT985806 |
| *Potentilla aurea* L. | CM:359702 | KT985678 | KT985807 |
| *Potentilla biflora* Willd. ex Schltdl. | Feng 102 (HIB) | KP875301 | KP875270 |
| *Potentilla brevifolia* Nutt. | RM579088 | KT985681 | KT985810 |
| *Potentilla cappadocica* Boiss. | E:00409794 | KT985683 | KT985812 |
| *Potentilla carduchorum* Soják | E:00081576 | KT985685 | – |
| *Potentilla caucasica* Juz. | GB:Antonelli A. #346 | FN430802 | FN421378 |
| *Potentilla caulescens* L. | MA 133 (GB) | FN430819 | FN421379 |
| *Potentilla chinensis* Ser. | Feng 110 (HIB) | KP875298 | KP875266 |
| *Potentilla chrysantha* Trevir. | GB:Topel M. MA142 | FN430803 | FN421385 |
| *Potentilla clandestina* Soják | Feng 25 (HIB) | KP875308 | KP875274 |
| *Potentilla collina* Wibel | Gray (A) | KT985689 | KT985817 |
| *Potentilla conferta* Bunge | Feng 127 (HIB) | KP875296 | KP875264 |
| *Potentilla coriandrifolia* D. Don | Feng 133 (HIB) | KP875302 | KP875269 |
| *Potentilla crantzii* (Crantz) Fritsch | S:Eriksson T. TE 703 | FN555609 | - |
| *Potentilla desertorum* Bunge | RSA:376320 | KT985696 | KT985824 |
| *Potentilla discolor* Bunge | Feng 118 (HIB) | KP875299 | KP875262 |
| *Potentilla dombeyi* Nestl. | Romoleroux 4579 QCA | HM453948 | - |
| *Potentilla elegans* Cham. & Schltdl. | GB:Eriksen B. 1440 1 | FN430779 | FN421358 |
| *Potentilla elvendensis* Boiss. | E:00201630 | KT985701 | KT985829 |
| *Potentilla erecta* (L.) Raeusch. | MA 124 (GB) | FN430780 | – |
| *Potentilla fedtschenkoana* Siegfr. ex Th.Wolf | RSA:376316 | KT985704 | KT985832 |
| *Potentilla flabellata* Regel & Schmalh. | RSA:376317 | KT985706 | KT985834 |
| *Potentilla flabellifolia* Hook. ex Torr. & A.Gray | GB:Topel M. MA164 | FN430810 | FN421392 |
| *Potentilla fragarioides* L. | Cult. in Hortus Bergianus | FN555610 | – |
| *Potentilla fragiformis* D.F.K.Schltdl. | GB:Eriksen B. 540-1-05 | FN430790 | FN421386 |
| *Potentilla geranioides* Willd. | E:00409804 | KT985709 | KT985837 |
| *Potentilla gorodkovii* Jurtz. | GB:Eriksen B. 930-05-1 | FN430800 | FN421380 |
| *Potentilla grandiflora* L. | GB:Topel M. MA149 | FN430806 | FN421400 |
| *Potentilla griffithii* Hook. f. | Feng 44 (HIB) | KP875293 | KP875261 |
| *Potentilla hirta* L. | RSA:460710 | KT985712 | KT985839 |
| *Potentilla hispanica* Zimmeter | RSA:532516 | KT985713 | KT985840 |
| *Potentilla hololeuca* Boiss. ex Lehm. | RSA:376303 | KT985714 | KT985841 |
| *Potentilla humifusa* Nutt. | E:00409788 | KT985716 | KT985843 |
| *Potentilla indica* (Andrews) Wolf | Feng 138 (HIB) | KP875300 | KP875268 |
| *Potentilla intermedia* L. | 69624HIM | MG236232 | - |
| *Potentilla kleiniana* Wight & Arn. | Feng 139 (HIB) | KP875294 | KP875263 |
| *Potentilla kotschyana* Fenzl | E:00409661 | KT985720 | - |
| *Potentilla kurdica* Boiss. & Hohen. | Kordestan Natural Resource Research Center Herbarium 8060 | AB894153 | - |
| *Potentilla matsumurae* Th.Wolf | CM:263168 | KT985727 | - |
| *Potentilla maura* Th.Wolf | E:0063802 | KT985728 | KT985854 |
| *Potentilla megalantha* Takeda | CM:382796 | KT985729 | KT985855 |
| *Potentilla meyeri* Boiss. | E:00409447 | KT985767 | KT985856 |
| *Potentilla montenegrina* Pant. | S:Eriksson T. BGE#3 | FN430782 | FN421361 |
| *Potentilla multifida* (Tausch) Wolf | Feng 124 (HIB) | KP875295 | KP875265 |
| *Potentilla nepalensis* Hook. | Topel M. MA 163 | FN430821 | FN421390 |
| *Potentilla nervosa* Juz. | Arnold | KT985733 | KT985859 |
| *Potentilla neumaniana* Rchb. | S:Eriksson T. BT#1 | FN666607 | FN421370 |
| *Potentilla nevadensis* Boiss. | E:00663775 | KT985734 | KT985860 |
| *Potentilla nivea* L. | isolate="NIV1" | MW042167 | - |
| *Potentilla pannosa* Boiss. & Hausskn. ex Boiss. | TUH:64940 | AB894155 | - |
| *Potentilla patula* Waldst. & Kit. | E:00500251 | KT985743 | KT985869 |
| *Potentilla pectinisecta* Rydb. | RM763737 | KT985744 | KT985870 |
| *Potentilla pedersenii* (Rydb.) Rydb. | GB:Eriksen B. 05-24 | FN430799 | FN421404 |
| *Potentilla pensylvanica* L. | POPECO25-290609 | MF543804 | - |
| *Potentilla persica* Boiss. & Hausskn. | TUH:24898 | AB894156 | - |
| *Potentilla petraea* D.F.K.Schltdl. | TUH:36632 | AB894157 | - |
| *Potentilla pulchella* R.Br. | E:00663796 | KT985747 | KT985873 |
| *Potentilla purpurea* (Royle) Hook. f. | Feng 64 (HIB) | KP875307 | KP875275 |
| *Potentilla pyrenaica* Ramond ex DC. | E:00128293 | KT985749 | - |
| *Potentilla radiata* Lehm. | TUH:17184 | AB894159 | - |
| *Potentilla reptans* L. | MA 131 (GB) | FN430815 | FN421368 |
| *Potentilla ruprechtii* Boiss. | E:00409744 | KT985753 | KT985878 |
| *Potentilla sischanensis* Bunge ex Lehm. | Feng 112 (HIB) | KP875297 | KP875267 |
| *Potentilla stolonifera* Lehm. ex Ledeb | BE 1382: 1 (GB) | FN430814 | FN421363 |
| *Potentilla suavis* Soják | Feng 37 (HIB) | KP875305 | KP875276 |
| *Potentilla subgorodkovii* Jurtzev | RM521873 | KT985761 | KT985885 |
| *Potentilla subvahliana* Jurtzev | GB:Eriksen B. 931-3-05 | FN430783 | FN421364 |
| *Potentilla supina* L. | Q567 | MH711447 | - |
| *Potentilla tabernaemontani* Asch. | S:Eriksson T. SG#1 | FN555608 | FN421365 |
| *Potentilla tanacetifolia* Willd. ex D.F.K.Schltdl. | S:Eriksson T. ex. Leipzig-98 | FN430797 | FN421366 |
| *Potentilla tenuis* (Hand.-Mazz.) Soják | Feng 26 (HIB) | KP875306 | KP875273 |
| *Potentilla tetrandra* (Hook. f.) Bunge | Feng 89 (HIB) | KP875303 | KP875271 |
| *Potentilla thuringiaca* Bernh. ex Link | GB:Topel M. MA119 | FN430777 | FN421406 |
| *Potentilla uniflora* Ledeb. | GB:Eriksen B. 271-4-05 | FN430785 | FN421367 |
| *Potentilla verna* L. | CCDB-18317-G06 | MG236614 | - |
| *Rosa majalis* Herrm. | T. Eriksson 641 (GH, S) | U90801 | FJ422371 |
| *Sanguisorba officinalis* L. | T. Eriksson s.n. (GH) (for ITS); T. Eriksson 804 (S) (for ETS) | U90797 | FJ422372 |
| *Sibbaldia parviflora* Willd. | M. Lundberg 4 (S) | FJ356174 | FJ422374 |
| *Sibbaldianthe sericea* Grubov | Feng 122 (HIB) | KP875312 | KP875285 |

Table S9. GenBank accession numbers of publicly available ITS sequences downloaded for the *Erythronium* phylogeny.

| **Taxa** | **Voucher specimen** | **ITS** |
| --- | --- | --- |
| *Erythronium dens-canis* (Spain, Santiago de Compostela) | Romai 68296 (SANT) | KP684250  KP684251  KP684252 |
| *E. dens-canis* (Romania, Vadu Crișului) | Bartha 664526 (CL) | KP684253  KP684254  KP684255 |
| *Erythronium caucasicum* | Volkova 664523 (CL) | KP684256  KP684257  KP684258 |
| *Erythronium sibiricum* (typical form) 1 | Stepanov 664955 (CL) | KP684259 |
| *E. sibiricum* (typical form) 2 | collected by Andrew Pyak | KP684260 |
| *E. sibiricum* (typical form) 3 | Konovalova s.n. (MHA) | KP684261 |
| *E. sibiricum* (typical form) 4 | Rukšans 664954 (CL) | KP684262 |
| *E. sibiricum* subsp. *altaicum* 1 | Alexander 664953 (CL) | KP684263 |
| *E. sibiricum* subsp. *altaicum* 2 | Rukšans 664952 (CL) | KP684264 |
| *E. sibiricum* subsp. *sulevii* 1 | Sviridova 664951 (CL) | KP684265 |
| *E. sibiricum* subsp. *sulevii* 2 | Rukšans 664950 (CL) | KP684266 |
| *Erythronium sajanense* 1 | Stepanov 664958 (CL) | KP684267 |
| *E. sajanense* 2 | Stepanov s.n. (KRSU) | KP684268 |
| *E. sajanense* 3 | Stepanov 664956 (CL) | KP684269 |
| *E. sajanense* 4 | Stepanov 664957 (CL) | KP684270 |
| *Erythronium japonicum* 1 | Rukšans 664949 (CL) | KP684271 |
| *E. japonicum* 2 | Allen 8702 (University of Victoria) | AF485283 |
| *E. japonicum* 3 | Chase 780 (K), Kew 1979-5130 | EU912083 |
| *Erythronium albidum* | Ebinger 28288 | AF485301 |
| *Erythronium americanum* | Allen 9915 | AF485304 |
| *Erythronium helenae* | RBGE (living coll.): 19861504 | AF485293 |
| *Erythronium multiscapoideum* | Allen 1240 | AF485291 |
| *Amana edulis* | K:DNA:MWC2397 | HE656027 |
