## Supplementary material for "Intact rDNA arrays of *Potentilla*-origin detected in *Erythronium* nucleus suggest recent eudicot-to-monocot horizontal transfer": Dataset S1

**Dataset S1.** Raw sequence of Nanopore read 202d7c58-6078-4612-8e8a-bf45fa7b7b04 containing parts of both native and *Potentilla*-type rDNA.

>202d7c58-6078-4612-8e8a-bf45fa7b7b04

GCATGGTCAGGTGAAATGTTCGGATCGCGGCGGCCAGCGGTCCGCCCGCCGTGACGTCACGAAGAAGTCCATTGAACACCATCATTTAGGAAGGAGAAATGCGTAACAAGAGTTTCCGTGGTGAACACTACTTGAAGGATCATTATCGATACCAGCCCAAAGGCATGACTTCCATGGCGGGATATCACAGGTTATGGCTAAACTCGACCTCCACTGGACCCCGTCGCCCCCTTCCGGGCGACCTTGTGTTCTGGATGGGGTGGCGAATGCAGGATACTTGACCCGTGTGCCTGCCGAAACCTCATGCGGCCCGGATGGACGTGCGCCTCTTGCCCTGTGGTGAAACCGCGGCAGCGAACATTATCATCGACTAAACCGCTGAATGTACAAGCTCTCTGCCCTTTCAGTAAACGCTGCGAAATGAAATACGCAACGGTGTGGAGGAGTCAGCTAAATACTAAGTGCTGACATAAATTCGCATGCTTAGAGCCTTCGGCGACCCAGCGCACTGCAATCACGCCTGTGTGGCTCTATCGACCCTTGAGTGGTGTTGATGTGCGAAATTGGCCCCCATGCCTTGTGTGCGGTGGGCTAAAGGGGAAGTTTGCCATTTAATTAGAACCTGACAAAGTGGTGGACATAGCGCCCGGCGAGGACACGTGGCCCCCTGGTGGAAGGACCCAAGTGACCCGGATAAGGTGGCTGCACTCCCCATGCAAGGAGATTGTGTGCCACCATAACCTGAAACCAATTCCAGAAGGCAGGGACACCATGAGTTCGCCCATATCAATAAGCGGAGGAGAAAAACTTACAAGGGATTCCCTAAGTACTTGGCGGCGGGCAGGGATCGGCCAGCTTGAGAATCGGGGGGCCTCGCCCCTCAGTGTAATCTGAAGCGTCCTCGGCGGACCGGGCAGAGTCCGGCTGGAAGCGCGCGTGGAGGTAGAGCCCGTCAGGCCGGACCTTGCCGCACCACGAGCGAGCGGCCAATCAGGTTGGTAGATACAGCCCAAATCAGGCGGTAAACTCCGTCCAAGGCTAAATCTTGGCGAGACAGTAGCGAACAAAGTACAGCGAGAAAGATGAAAAGGACTTTGAAAAGAAGTCAAGAGTGCTTAGAGAATTAGGGAGAAACGGATGGGGAAGCGTGCGCCCTCGGCGAGATCTTTGAGCAAGCGTCAGGCCAGAATCCGCGCTTCAGCTTTCGAGAGTCTGCTGGTGGATGCAGTACTGCTTGTGATCGCCGGATGATCCCTTGCGTGGAATCGTCCCGGTCTGGCAAAGAAGCCCGGCCTTGGCGCCGTTAAGCGTGCCTAGTGAGCCCACTGCCCGGCGCCCCAGCACAACAGGCGGCTCCCCATCGACCCGTCTTGAAACAGAGATAAAAATTCGACATGCGTGCGAGCGTGGCGTGAAGCCCGGAAGGCGAAAAGGAAGGCTGATGGGCGGGATCTTCCCCACCTACGGGGTGCGCCGCCAAGCCAGGCCCCCGTCTCATGAAGGGTTCGAGGGCAGGGCTTAATGCAGCCGGGACCCGAAGGGGCTGGTGAACCATACCTTTTGTCCAATTTAATAACTTTGAAGTGATACAATGTGCTCTCACAAATGTGGATGCCAAAATGCAAAGAAAAAAGTCACAAATACGAGTCCTAAGAGGCAAGCTCTCAAAAATTCGCTTGGCGCAATATTTTGTTCCCTCCAGGCAAAGAACTGGTCATCCAAAAGCCTTTCTTTAATCCTCCAAAAACAAAGTGCGCACAGCTCCAACCGATATCAGCCCGTCTACAGTGGGTTCCTCTTCATCTGCACATGATATCTGAAGCCCTGCCATGAATGATATTCCTGTTTGCTTGTCTTTGTTGCTCTTAGGTAAAAATTAAAGTGCTCACAGTACACTCAGCGGCAGTGCTAAGAATCTTCTTTGGAAGATGCAGTGAAGTACGGTGGCATCTCAATTGTTACGAGGTTGTGTACTTCTTTGTTTTCCTCGCTGACGAAGTGAATAACCAACCTCATCGATGCTTCCGCTTTCTTGCATCATCATTTAACGCTCTCAAGTAACATCACATCAGCCCGCTTTCATCCACGAAACAGCCTGTTAGAGTTACAACATTTCAGAAGTAGGATGCATAGAAGGTATGCATTTGGGTACAGTCAGTTGTAGCATTTAGTGGTTGAAATCGGCCTCAGCTGGATATACTTTTATCTATGGTGCATTCTCTACAGAAATACCACGAAGAAGTGTAGTCTTGACTTCAAGCCATTGTAAAGCCGTGTACGATTATTCGTTAAATTTGTCACTTTATACCTCAAAAATCGTTCATGAGGAGTTATGTCATCCATACATTTGCGGAAAAGAACATCAGTTGGGACCTTCAAAGCAATTGCACCTTGACTAACCATCTGGCTGCTATCTGTAAACCTTTATGAAGTGGGTTAGGGCATCGATCTCAACAAATCCGAGGATGTGCGGAGCTTGAGTGTTGCTAGTCTCGGGATCGAGCCTCAACAAATCGAGAGCAAAGGAGTACCAATGGTGAAAAGAGATTGTCGGAGGCGCAAGCTCGGGTTTGTGTTACAGTAGCGTAGCCACTTGTTGCGGAGGTCTGCCTTCCTTCCCGAGGCTACTTTGCCAAGTGCAGACAAGAAGTAAGGTACCTTATCCAATAATCCCATGGTTCCTTGAAATGCTTGTGTGAGGTCTTGTGTTCATCGGCTGGGTCAGGAATTCTTCTAGGAGTCGAACTCCCTTTGCTGTGGCAGAGGAATCCCACGCCCATCTGTGATACCTCAATTGCTTTGCAAAACCAGGCCTTACCACAGTGAAGCGTAATCAAGGTGTGAGCTAAATTAGCTTTAACTTTCTCCTCATAGTCAATGATGCTTATGTGAATCGATGGCACTCACTTAAGTATATGTATACAAGAACCTCATGATCAAATTACGGTGAAGGAATAGGTAGCAGCGGATTTCTTGAAGCAATAGCATAAGATCTAGGTTCTTCAATACGGAAGGCTGCTTTCTGCAGCTTGCGTGTGCTCTCCCTCTGTTTGATCTGAGCTTTGATAGCGTTTAGATAGCCGGATGTTCAGTAAGGCTGGCGGTGTCGAGCTCGGATGCGAGTTGGTCTTAGATGTAGCTTTCTTGATCGGTAGAGAAATTGAGTGAATTCTCAGTGTCTCTACATTCTGGCTGTAAATATTCGAACTCAGTTTGTAACCTTGAGTGGTTAAATCGAGGTTCAAGTTTGTAACCTTGATTGTTCCAGTAGAAACACCCTTGACTGAGATGGAGTCTGTACATCGGGATTGGGGATATATCATTTCAAATTTGAAGAAGTTGGTTGCATACGGAATACGTGCTCCTATACGCAAGCTGAACCGAGGCACTCATGCTTCCATCAGTGTAAAGAATCGGCGCCATGAGTAGTCGGAGTTGAAGCTCTCTGGCGGGAACATCATGACGTTCCCTCCCTCACGCTTGAACCAGCCCGAGGAAGGCATGCCCGAATCTTCTCCCTCCATGATCGGGCATTTGTGCCATTCATTGAGGAGTATCTTTGGCCTTAGCATTCCGTCTCTAGTCCACAGGCTCTGTGAGTGGTGCAATCTCTCCACCAAAAACATTCTGGACCAGAGAGTACATCACCTTGTGAACCCGTCCTTCGGGTTAAAGTTGGCATCTCACTTTAGTAGATCCGCATTCCCAGTGAGACCATGTGCCTCCCAGAGGAGCTTGCCCTTCCTTCTGTCCAAAGCAGTAAATAAGCCACGTAACTTCTCCCATAGGCGATCTCCCCCTCGGCTCAGGCGGCGCAGGTGTGAACCTCACATTCAGGTGCCTTCTTGGTCATCCGGAGGAGGCCTAATGGCGCTCCATCGGAGGGAGGAAGGAGGAGGAGGAGGGCGGGGCGGCCAGAGAGCTCACGGTGGAGGCCCGCGGCCGATGTGGCGTGCAAATGGTGCATCCTGACTTGGTGGCGAAAAATGTCGAACGTCTAATGGCGGTTCCCTCCGAAGTTTCCCTCGAGATGGCCGGAGCCGTGAGCGAGTTCATCGGAGTGGCTTAATGAGTTAAAGCCATGTTTTCGTGCCTCGACCTATTCTCTTTAAATAAGTGGGGCCGGCGCATTTCTTTGTTGAGCAACCATGGAATCAGGAGCTTAGTGTTGACCATTTTAGTAAGCGAAGCTAGCGTGCGGGATGAGCGAGCGTTCTGTGGTGCCATTCGCGCTAACCTAGAACCTGAGGTGTTGGTCGATTAAGGCAGCGGGACGGTGGTCATGCGTCAAAATCACAGCTAAGAGGTGTGCCAACAACTCTGCCGAATCAAACTGGCCCGAAAATGGATGGCGCTTAAGCGCGTGACCGGCCCATGGGGCAAGTGCAGGCCCGACCAGTGGGGCCAGTCATGCAGAAACACAGGGCGTGAGCAGCGGAGCGGCCCGTCAGTCTGCGAGATCTTAGTGGTAGTAGCAAATATTCAAATGAGAACTTTGAAGAGGCAAAGAGGAAAAGGTTCATGGGCACGGCACTTGCACATGGGTTAGTCGATCCTAAGGAAGCTGAGTAACCGTCCGAGTAAGCGCGTCGCGAACTTGAAAAGGAATCGGGTTAAATTCCTGAAGCAGGGACATTGGTGATTGGCAATAACGTTCGAGAGTCAGGAGAGCGTCGGCCCGGGAGCCTCGGAAGAGTTATCTTTTGTTTAACGGCACGCCACCACTGAAATGAAAGCTCGGTAGGTAAGATCCCGGCCATTTGGAAGAGCACCACTGCGTCATGGTGTTGGTCGCCCGGTGACCCGCGAAATCCGGAATGAGGTGCCATTCACGCCCGGTCGTACTCATAACGCATCGAGGTCTCCAGGTGAGCGGCCTGCGTGGAGTGGAATAATGTAGGCAAGGAAGTGTGAAATGGATCCGTAACCGGGGAAAAAGGATTAGCTGCGAGAAGTGCAGGCACGGGGGTCCCAGTCCCGAACCCGTCAGTTTGTCGTGACTGCTCGGTGCTTCGCGGCGGGCGAATGCATCAGCGTCTTGCTGAAGACAGTAGAGCGCTCTGGGGGCTTTCCCAGGCGTTCAACGGTCGACTGAACATGCGGACGAAGTCGCATTTAATTAAAACAAAGCATTGCGATGGTCCTGCTTCGGATATTAGCGCAATGTGATTTCGCGATTACTCAGATGTCAAGTGAAATTAGCTAAAGCGGAGTAAGCGGCGGGAAGTAACTATGACTCTCTTAAGGTAGCAGAATGCCTCAAATCGTCTAATTAGTGGCCGTGCTGGAATCAGTTAACAGATTCCCCACATCCTGTCTGCCTATCGAGCGAAACCAAGCCAAGGGAGCGGGCTTGGCGGAATCAGCGGGGAAAGAAGACCTGTTGGCTGGCTCTAGTCAGGCTGTGAAATGACCCGAGAGGGTGTAGTATAAGTGGAGCCTTCGGGCGTGAAATACCACTACTTTTAACGTTATTTACTTATTCCGTGAATCGGAGGCGAGAGCCTTTGCCCTCTTTTGGACAAAGCCCGCTTAGTAGTTTGATCCATGAGAAGACATTATTGGTGAGAGTTTAGGTAGGCGGCTTTATCTGTTAAAAGATAACGCAGTGTCCTGAATGAGCTCGCGAGAGCGAAATCTCGTGTGGAACAAAAGGAGTAAAGCTGTGGTTGATTACAGTTTAAGTGAATCTGACAGTAAACTCTCGAGGTCTTCAAAGACCTTTGGAATTTAAAGCTAGAGGTGTCGGAAAAGTTACCACAGGGGATAAGCGTATGGCGACCAAGCGTTCATAAGCGACATTGCTTTTTGATCCTTGATGTCGGCTCTTCCTATCATTGTGAAAGCGAATTCACCAAGTGTTGGTTCTTCACACAATGAGAGCGTGTTTGGGTTTGAGCCGTGTGAGCGAGGTTGTGGTTCTCACGTGATGGCAGTGCAATAGTAATTCAACCTAGTACGAGAGGCAGTTGATTAGCACGGTGGTCATCGCGCTTGGTTGAAAGCGATAATGGCAAACTCTCCGTCATGACTGGTCTGGCTGACGCCTCTAAGTCGAGAATCCGGGCTAGAAGCGATGCATCACGCTGTGTTCATTTGCGACCCGCAGTAAATTTCCAGGCTGAAAACCACGTCGTTAGTCGAAGCCCTACTTAGGACAAGTGGCGCCTTAGTACAATTTCTAGGCGGCGGGTAGAATCGCTATGACGGCACCATCAGGCGGGGTATTGTAAGTGGCGAGTGGCCTTCTTTGCCACGATCCCTTTGAATTAAACCTGATGAAAGCACGTTAATCCCTCCCTTCATCTTCATTTTTCCTCCTCCTCCCAGCGAGGCTAGCCTCTAACACTCTATGCTTTGAAGTCGCGGTCTTGAAGCGCTCAGCCGTGATACCAGTTCACTGTGGCCTTTAGGCTACGAACCGCCTAATCGCGAGATCGATAAGCGAGGCTGGCCTCCAACACTTAGCCAAACTTACTTTCGAGGAGGTAAAACTTTCGTGGGAGTGTCTTGTCATGATGCTTTGAGTGAAAAATTCAGAAAATAGAAACTTGTGATTTTGGCCCAAAAACCGTGTGCTATAGCTATGTCAGCCAATCGGCATTTGGACGTAACTCATAAACCACTTGTGATTGCAGGATTCGATGAACATGAAAGAACTATATCATCATGATGCAGTGGGAAAAATATGGTGAAAACAAAGACTTGCGTTTTTGGCCCAAAACGCATGTCTTTATAACCCACAAAAATCGAAAGCGAAATCGGCATTTCGGACCGTAACTCATAAATAGCATGATTTTGAGTTAAAATCCGGTTAAAAAGAAGGGCTTATATATCATCATGTCTGAGTGGGAAAAATATGAGGTGAAAATAAAGACTTGCGTTTTATTTAACCGTGTCTTTATAACCCACAAAATCGAGCGAAATCGGCATTTGAGGCGTAACTCATAAACCGCATGATTTTGAGTTCGCGTCGGTTAAAGAAGGCTTATATATCATCATGTCTGAGTGGGAAAAAATGATTGAAAATAAAGACTTGCGTTTTTGGCCCAAAGCGTTGTCTTACTCAACAACCGAGCACAAATGACATTTCAGGTGCAATTTTTTTTTTTATAGTTTCTCCAGCTCTAATGATGATTATGAACATTGTTTAGTGTGAATCTAAAGTTTCTAATATTTTTGAGGAAAATTTATGTTTTAGGTTTTGTGGAAAATAGAGTGAACCACAGGTACCAAATGGGCATGCCAACATTTGTATATATATATATATAAGAGGGCAGTGTCTTGAGTATACGAGAGGCTAAAGTATGGATGCCGGGGTTTGAGGGTTGCCTAGGTGCAAGGCATGGATCTGAGGGCAGGTGAAACAGGAGTAGCCTCCCAGGGAGGCCTATGGATGCCGAGGGTTTGAGGGGTTGCCTGAGGTGCAGACATGATGCTTAGAGGTGCAAAAACATGTTGAGATTGAGATCGCGCAACATCTTTGAGAAGTGCCAGCGGTCTCGTAGCGTGGTGCCCAAGGAGTCTTTGTCGCGTCTGCGGCAAGAAATTTCTGGCTACAGTGGATCTGCGTGCGCGGCATGAGCGAGGCTGACAGTCTTGAGTCAAAGAGCCCACCCAAGAGGCTGCAGTGGATGCAGCGGTGGCGGCAAAGTGCGAGAGAGCTGCTTGTCTGTGGCTTATGTTGCCGAATCTTGCCGCGATCTTGTGACAAGAAAAGGAGGCTGTTCTTAGGAGTCTCCCAGCAACATGGTACGAGGGAGTGCAGCAGTCTGACAAGGTTGCGGCAGAGTTGCGCGGTGTGCGAGAATATGTTGCAGTCTGCCGATGCAGCAAGGCTGCGAGGATACGCAGTGTGCAGCATAAACTGCAGAAAAGGCTACTGATCACGACAAAGGCTAAATCGGGTTACGCAGTGTCTGTAGGCTAGAGGTGATGCGCAATCTTATGGCTGGCTGCGAGGATCTGCGATGTAGCGAGGAGGCTGCGAGGATGCGCGGTCTCTTGGCCGAGGGGCTGCAGTCTTTTTGTAGTGCCAGTCTTTGTCGCGATGCGCGACCTTCTTTGCTGAGGAGCTACCGAGGGGTTCCAAATTCATGATTTTCCACTTTTTGTGAAACATGGGTGAACTCCGTGCCTTGATGTTTACAATTCCATATAGTAGGGGTTGGTGCTTTGGGCAAGGATGTGGGAATGGTTTCGGGTGGCATGTGATTTGTTCTACCAAGTAAGGGGCAAGTTATTTCCATAGATATGGTTCTCGGGTTTGTTTAAGCTGGATACTCAGCGCATTGTGCTCTAATGTTCATTCTCGGTGAGTGAGTGAGCGTGGTGTCCAACAAGTCTGCTGGTTCTGTGGGGGTTTGCCCTCCGTGGATCTGCAAACCTTTGCCTCGAAAGTTGAGCAGAGAGTCGGTTTGCTCGCAAGGCCTATTGGGTGGGTGCTTTTCTATGTTTTCCATCTTTGTGAGTGCAGACCCAGCTCACAGCCAAAGCCAGTTTAGATTGTCAAACACACATTAGTGGGCGTGGTCGGTTTAATGTTCTTTATGATGTGATGTAGCAGTGTGAGTTGTGAATCGGTTGTGTGGGTTGTAGGGCTCCGTGCTTCGGCCTTCGAACCGTCCGCCCACACACCACGTCGATATTGTAACCAAGACTTTCTTGTGCTATCTGGAGATTCTGTATTTTCCTAAGCTTAAAGGAGCAGTGTCGCTCGCTACCCTTTCCTTCCCACAGCTGCGCTGCGAGAAGATGTGTATGGCGCTTTGTGCCCCTCGCACTCCGAGAAACGCGCCGTAAAGCGCGCCGCGTTCGCGGCAGTTCTCAATTTGCTACTTGATGGTTCGAGATCTGAACAATGGTGACAGTGTCGGCTTACCTTCGATGTCAAATTCAAAGCGTTTTGTAAAACTGAACGCTATGCATTGTCGTTAAGCTACTTGGTCTCGGCCTTGATGTATCAGCGTCGGCAAAGAATGCTACGAGATTTGATCACGCGATGCGGTCATATGCTTGTCTGAAAAGATTAAGCCATGCGTGTGTAAAGTATATTCATTAATTGTGAAAGCTACGAATAGCTCATTCATCGGTCTCGTTTGTTTGATGGTACCTACTACTCGGATAACCGTAGTAATTCTAGAGCTAATCCGTGCAGCAAATCCCGACTTACGGAAGGGATCATTTATTAGATAAGGTCATAGCGGGCTCTTGCCCGATTTGTATTAATTCATGACCTTAGCCGGATCGCCTGGCCTTAGTGCGGCGACGCATCATTCATACGCCTATCGCTTCGATGGTAGGATGGTGGCTGCCATGGTGGTGGCGGGTGGCGGAGGAGTGGGTTGATTCGGAGAGGGCACGAGAAAGCAATACCACGTCCAAGGAAAGCTGGTAACAATTACCCCAATCGTACGCGGGGAAGTAAGCTGACAATAAAATAACAATACAGACTCTCACTAAATTAGTAATTGGAATCAAGTACAATCTAAATCCTTATGAGAGATCATTAGAGGAGCAAATCCCGAGTGCGTGACATGATAATTCAATCGATAGCGTATATTAAGTTATTAGTTAAAAACAAGTAGTTGGACCTTGGAGTTGCGGTCGACCCGGTCGCCTATGGGTAATGCACCCGGTCGGCTGCTGCGTCCATTCCTCTGGCGATAAACCGGCGCCAATTGGCAGATGCATTACCTCGATGCTGATACTTTGAAGAAATTAAAGTGCTCAAGCCAAGCTAGCTCTGGATACATTAGCGTGGGATAACATCATATAGGATTTCCGGTCCTATTACATTGGCCTTGGGATGAGTAATCGTGCGGGGGCAGAGTGAGGCATTAGCTATTTCATGGTCGAGGTGAAATTCTTGGATTTATGAAGAGCGAACAGCTGCCAGCATTTGCCAGAGATGTTTTATTGTCGGCGAAAGTTGGGGGCTCAAGCGAATCGGATAAGTCTGATCTCAACCACCAGCGATACCGACGAGGGTCGTGGATCATTCACTAGGACTCATGGCGCTACTATGAGAAATCAAAGTTTTAGGTTCGGGAGTAGCAGTGTAAGTAAACTTGAGGTGGCGGAAGAGGCCTGCAGTGGAGCACACGGCTTAATTTGACTGTGCGACTTACCGAGTCCGGACATAGTAAGGATTGTCAGTGAGAGCTCTTTCATGATTCTATAGGTGAGCAGGTGCATGGCAGTTCTTGGGTGGTGGAGCGATTTATGCAGTTAGACTGGGTGGGCGAGACCTCTGACTGCTAACCCCGCTATACGGAGGTCTCCCTCATGACGGCTTCTTAGGCTATGGCCGCTTAGGCCAAGGAAGTTTGAGGCAATACTTGGTCCGTGATGCCCTTAGATGTTACAGACCTGCGCTCTGTGATGTCATTCAACGAAGTCTATAGCCTTGGCCGGCGGGCGGTGTCTTTAACGAAATTTCATCGTGATGAGGGTAGATCAAATTCGATTGTTGGTCTTCAACGAGGAATTTCATCAGCGTAGTCATGGCTGGCGTTAACTCGTCCGCCCTTTGTACACACCGCCGTCGCTCCTACCGATTGAATGGTCGGTGAAGTTTTCGGATCAGCGGCCGGCGTCGGCCAGGTTCCATTGTCTGCGGCGTCGCGAGAAGTCCACTGAACCTTATCATTTAGAGGAAGGAGAAGTCGTAACGAGGTTCCAGTAGGTGAACTGGAAGGATCGATATCGAAACTGCCTAGCAAAGCGAGCCGAGAAGCGTCATTTCAGCGCTTGGAGGCAGGGGGCCTCGCGGCTCTCGCCTCCTCATCCGAGACGAAGCCTCGCGTGTCGTGCCCGGCGCTTCACGCTTAAGCGACCTCTCCCGGGCGTGCGGCCTCGGCGTGGTGCGCCAAGGAACTTGAATGAAGGAGCGTCACCGCGTCCCCGGAGGCTGATGGCTGTATGAGTGGTTCGTCGTCTTGATATGTCTAAGCGACTCTCGGCAACGGATATCTCGGCTCGCATCGATGAAGCGTGGCAGAAATGCGATACTTGGTGTGGTGCGAGAATAGCAATACGTAACT
