## Supplementary material for "Intact rDNA arrays of *Potentilla*-origin detected in *Erythronium* nucleus suggest recent eudicot-to-monocot horizontal transfer": Dataset S2

**Dataset S2.** Raw sequence of the longest Nanopore read (e16c7cdb-900b-4c49-8019-e4f837b9606f) containing rDNA of *Potentilla*-origin.

>e16c7cdb-900b-4c49-8019-e4f837b9606f

CGGTATACTTCGTTCAATTTACGTATTGCTAAAGGAAGACCTAAGCCGCCTTCCAGACAAACGGAAAGCCTCTTGACATACCCGGCTCAAATGGCAAGAAAACATCAGCACCAATCCGATTCCAGTTCGCGGCGCTATTCTACAGTCTGAGATACGATTGTTAAAGACCCCGGACCCGGCCAAATCGAGAAGCACGAGATCAGACACTAGCTTTTCAGGCATTCCTTCCTTGGTCTGACTACGAGATAGTGAAGACCCTGGATCACTCCTTATCGGATTAGGAGCCGGAGGCACCTTATTGAGACACTTTCTTATGTATTTGAGGTAAGCATACATTTAATCTAGGTGTTTGGGTTTCAATTGGCGAATTCATTAACTTAATCTTCGCGTTAGTACCACTGACCGCCTCTCTCCTTCGTTTATACGGGCTAAGCTCCCATGATCGAACAGAGGCAAGTGTGCTGTTGTTGCCAAAGCGACTTTACTAATCGGACCTTGGAGAGTGAGCAGGTCAAAAGAGATTCATCTGGCAGAAAATTGTATGTATAATTATCTCAGCCGCGAAACATCAAGGGTGCCAACGTGATGTTCAGCGAGGCAGTAGAACATTGACATTTCTTGTTTATTTGCCCTCTTTATATATGTGAATGTATAGCTTTAAAGCCAATAGTCGGCGTGTGACTCATGAAGGCTTATCTTGTATAGGCTGACTAGCTGGTGGCAACTTTCGGGACTTCTTGGTTAATAAAAGGCATAGTTCACCATCTTCCGGGTCGACGAGTATGCTCACACTCGAACCCTTCTCGGAAGATCAAGATCAAATTGTTGATTGCACCGAAAAGGATCCCGCACATTAAGCTTCCTTGCAGCCCCTTACGGAGCAATACGTTGACTACTTCTGTGTCAGGACTCAGCAGTGCGTGTTTGAGCGGGCGAATGAGGAAGCCCACGAGGCCGTTACGAGCACGCGAATCAAAAACTACGCCCGAGGCGGCGCGTCACCGCTTCCATGTCCAGCGTCGGCAGCGTCTCGCGGGCATATCGCGGCCAGACTTTAGCGCGCGCGATCATAGCAGTCCCACGCCACAGTGATGGCGGGCCGTATGTGGCCGTTCCACATCCCAGGCAAGAGTAGCATCGCCCGGCCCCCATCAGCTTCCCTCCAGCAGTTCAAGCACTCTTTGACTCTCTTTTCAAGTCCTTTTCATCTTTCTACGCGGTATGACCCCTTTCTCCAGGTCTCTCGCCCGTATTGCGCCTTGGGGCGAATTTGCCGCCGGTGGGGCTGCATTTAACAACCCCGACTACGCCAAGCGAGCGCCTCGTAGTGCGGCAGAGTCCAGGCACATGAGTCTCTCACCTACCCATGGCGCCCCCTTCCGGGGGACTTGTTCCCGATCCGCCCGGCCAGGCCTTCTCAGACTTCTGAATTAGGACCATGCGGCCTTGATTCTAAAGCTGCATTTATTCGAGTTAAGCTACGCCGTTACTAAGGAATCCTTATGAATTTCTACTGCTCATTATTGATATACTTAAATTCAGCGAGGTAACCCGCTTTGACACAGGGTCGCGTTGAAAGCCTTGAAGCGAAGCCAGCCGCGCATTAGATGGGAAAGCTTGGAGACTGACAGCGACAGGACGAGTTTGACAACCCATAGGTTAATATTGGGTCGCAGCTTGAAAGTTGTTGTTGTTGTTGTTGTTGGTATTGTTATTATTGGCGTTGTTGTTGTTGTTATTGTCAGCGCGGCGGAGGCGCCGCAGGAGCCCATCGTGAAGTCACCCAACTCCGGAGGTTGGAGGGCAACGGCGTGTGACGCCGGCGAGACGTGCCCTCGGCTGTGGCCGAGCGCGACTTGCGTTCAAAGCTTCGATGGTTCACGGGATTACGCAATTCACACCAGAAATTATCGCATTTACTCGTTCTTCATCGTCACAGAGCCGAGATATCCGTTGCGAGGCGTTTGACATATTGAAAGCTGACGAACCACCGGCACGATCACCGTCTCTAAAGGGGACGGCGGGTGTATCCTATTCAAGTTCCTTAGCGCGATTCACCTTGCGATGTTGATCGCCGGAGAAATTGACCAAGCGGGGCAGCGGCCCGACGAGCAACCGGATGAGCGAGGAGCGCGAGGCCCCGTCTCAAGCGTTGAAACCGTTCTCGGAGTCGTTACGCTGGCGGTTCGACAATGATCCTTCGTGAGTTCCACCTACGGAAACCTTATTATAACTTCTCTTCCTCTAAGCCAGTAAGGTTGATGGGCCTGGCGTCGCCGGGCAATGAAACCAAGCCACGTCGGCAGCGATCCGAAAACTTCATAGGACCATTCAATAGGTAGAGCGACAGGCGGTGTGTACAAAGGCGGGGCGTAGTCAGCGCAGCGATGACTCGCGCTTACTAGGAATTCTCGTTGAAGACCAACAAGTACAATGATCTATCCCCATCCATGATGATTTCAAGATTGCAGGGCTGTCGACCAAGGCTCTATAGACTCGTTGAATGCACTTCGATTGTAGCGCGTGCGGCCGAATATCTAAGGGCATCCACGGACTGTGTTGCCTCAAACTTCCTGGCCTAGCGGCGCGTCCCTCTAAGAAGGTGGCCATGGAGGAGACCTCCGCATAGCTAGTTAGCAGTGAGGTCTCGTTCGTTAGCAGGTGTGGACAATCGCTCCACCTGACTAAAACGGCCATGCACCACCACCCATAGAATAATGAAAAGAGCTCTGGATGCAGCGAGTGTACATATGCGGACACAGTAAAGTTTCCACAGTATTGGAGTCGAATTAAGCCATAATTCCACTCCTGATAGTGCCCTTCCGTCAATTCCTTTAAGTTTCGGCCTTGCGACCATACTCCCCGGAACCCAAAAACTTTGATTTCTCATAAGGTGCCGTGATCCTAAAGTAACATCCATGCGATCTGGTGGCCCTTCGTTTTATAGTTGAGACTAGGGCCGGTATCCCCATTCGTCTTGAGCCCCCAACTTCGTTCTTGATTAATGAAAACATCCTTGGCAAATGCTTTATGATTATTCGTCTTTCCTCGTCAAGAATTTCACCCTCTGACTATGAAATCGAATGCCCCCGGTACGTCTGTTAATCATTACTCAGATCCGAGAAGGCCAACGAATAGGACCGAAATCATGATGTTTATCCCATGCTAATGTATCAGGCGTAGGCTTGCTTTGAGCACTCTAATTTCTTAAGTAACCTGGAGGCGCAGCCCGGCAATTAAGGCCGGGAGCGTATCGCCCATTAGAAGACGGCCCAAGCCAGGGTGCCACAGCCAGGCGGACAGTCGGCCCAACCCAAGGTCCAACTCGGCTTTCATGCAACAACTTAAATATACGCTATTGGGCTGGAATTACCTTGATCTTGGCTTTGGACTTGCCCTCCGTGGATCTCGTTAAAGGTTTAGATTGTACTCATTCGATTACGAGACTCATAGAGCCCGGTATTATTGTTATTGTCACTACCTCCACGAGTGTCGGGATTGGAGTGATTTGCGCGCTTTCTTTTGCCTTCCTTAGATGTAGTAGCCGTTTCTCGAGAACTCCTCTCCGGGTCGAACCCTAATTCTCGAGTCACCGTCACCACCATGGTAGGCCACTATCCTACCATCGAAGAAGTTATTAGAGGCGAGATATTTGAATGATCGTCGCGGCCACTGAAATTTGTGCGATCCGTGAGTTATCATGAGTATCAATACAGCAGGCGAGCCCGCGTTATTACCCCCATCTATAAATGCATCCCTTCCGAGTAAGAGGATTTGTTGCACGTATTAGCTCTAGAATTTTCACGGTTATCCGGTAAGTAGGTACCATCAAACAAACTATGGCGGTTAATGAGCCATTCGCGGTTTCTGATGCTGAATTAGTTCATACTTACACATGTAACTTTAATCTTCAGAACAAGCATATGACTACGCGGGGATCAACCGGTAAGCATTCCTTTGCCGACCGCATGCCATGGCTGGAGGCCCTTGGCCATGACAGCGATCGCCATGTCTAGCGACAAAGCGCTGAATTTGGACATCGAAGGTGTGAGACACCCAAACCCACCAGATATTCCGTATCGAACCATCAAGTAGCAGCGAGAACCCAATGCAACGTGCGCGCCTTTGGCGTGCTCTCTGAGAACACAAGCCTTCTCGATCTCCCTCGCGGCGCAAGCTCGGGAAGAAATAGTAGCGGCAGCCGGCGCGTTCCTTTAGCGTGAGTAGCAAATACGGGAATCTCCAGGCTCGCTGCAAAGTCTTGCTGCGTCAGCGTGGTGTGTGGGCGGACGGTTCGATGCGAAGCCGCGAACCCTTGACCCACACAACCGATTCACAACTCTGCAGCTGTTACATGTGGGAACCAGCCCGACATTAAGCGTGCTTGAGCTAGCAGCGAGCGGAAACCAGCGATAAGCGATTTTATGTAGCCAAGCGCACCTTGACATGAAAACATGAAAAGCACCCACCCAATAGGCCTTGCAGGCATCAAAACCGACTCTGTTCCAACTTTTGACCGATGTCGTTAAGATCCACGGAGGGCAAGCCCCTACAGGGGCAGGCGAGACTTCGTTATTGGACTGCAACTCACTGACTCACTCACGAGAATGAACATTAGAGCCTGATCGCCTTGATATCGATTAAACAAACACAAGAACCATATCTATAAAATAGCACCTTACTTAAGTAGAACAAACTCACGTGCAGTAGAAACCATTCCACATCCCGTTCCAGACTACTGACCCCCTACTATATGGAATTGTGGCATGCCCATTTCAGCACCCAGGAGTTCACCCATGTTTCACAAAGTGAAATCATGAATTTGGAACCCCTCGCTGACCCCTCGCAGCTGAGGAGTTTGCGCGCGGCGGCACCTGTGAGACCTCGCGCGGCCACCTCATGACCTGCTGCGCGGCGCGGCATCCCGCGGCCTGCTGCGCCGCCGCGGCACATCCTCGCGGCCTTCTTTGCTGTCCAGCGACACCTCGGCGGCCATCACGCACCCATGTGACACGGCGCAGCTGGCTCGCACCGCGGCTGGCCCCTGTGACCATCTTTGCGCCTGTGACATCCTAGCGATGTGCTTTGGGCATGTGTGACACCTCGCTGACCATTACACTGCCAGCAACATCGCAGCCTTGGCTGCTGCATGCCGCGTGACCCCCTCGCTGACCATCTGCGCCCTCGCTGACCCCTCGCGGCCTTGCTGCGCATGTGACACCTCGGCGACGTCTCTTTGCGTGCGTAGCGACCTCGCGTACTTCGCGCCTGCGGCATCCGCTTTCGGCGGCCCCTCGCGGCCGCCGGCTACCGCCGCCGCGGCCCTGGCGTACTGCGCGCAGCGGCATCCACTCGGCGACCAGCTCGCGGCCTTGCTGCGCCGGCGAGCACCAGCCGTAAGCGTACGGCGCTGCGCTAGCGGCAACCCCCTCGGCGGCGTGTGCCAGCGCGTCTCGGCATCCCGCCAGCGGCCGTACTTTGCGCCTCGGCATTACCCCTTGCGCTAAAGCGACCTCAGGCCTCGGCATCCGTGCCACTGGCCTCTGCATGCATTAGGCTTGAGGTAGCCTTAAAAGGGAGGCTGAAGGTAGCAGATGCAGAAACTGAGGATGCTGTCTGGTATGCTTGCCCAGTCTGCTTTGTCTTGCAATAAGATCTTGTGACATCTTCTTAGAAAAGAAGCTCTTGTGTGCTGCCAACGTCTTCTGTCGCAATCTTTTTAGCAGTAGCGGCTATGAAATGCTGTGATGCGTGCGTATTTGCGAACTGCGGCGATAACGATCGAAGCTGAAAAGCTGCCGAAAATCAAGTCTGTGATCGGCGACAGGCTGCGAGGCGCTACGTCGCGACATGAAAGCGCGTCTTCCACCGCTGATCGCGGCGAAGAAGTGCTGAAGCACATGAAGCTGAAAAAAGCTGCGTGCAGCAAATTGCGACGGTCGCCCGATGTGCAAATCATGCGAATGTAAGCGTGCCAGTAAGTGAAGAGGAGATGCAGCGATGTGTGCCGAAATGCCCGTGTGCGAAGGCAGCGCTGCGTCAATTGCAGCGATAACGTAGTGAAAAAAATCTTCGCCCGATCTTTAGCGACAGGCTACGGATGCAGCGGTCTGGCAGAAGTGCGAAAGAAATCATGCGTCCTGCTGAAGATCGGCAGCGAATTCTGCGGCGAATGTTCAGCTCTCTTGCGACGCTGGCTGTGACGCACGTGAACATGGGTGAACTCTGTGCCAAATGAGCATGCCGCGGTGTTCAAATGAGAAGGCAACAACAGATGTGAAATGAACTGCGTATGTAATTTGTTCTACCAATGGTACAATTATTTCCATGAATAGTTCTGAGAGTTGTTGCTGAATATCGGCATTATGCTCTGGCTGTTCATTCTCGATGGCTCGGCGTGTCAACCTTTGGTGCACGCGTCGGATCTGCGACGCGCCTACGAATTGAGCAGTCGTTTTGCTCGCGAAAACTGTTGAGCCAGGTGCTTTCTATGTTTTCCATGGCGGCGATATGTAACTCTTCGCATAAATCGTTCGTGCTCGGCTTCTGGCTGATGAGGAGTGTCGGTTTGTGTGTGCGTCTTGATATGTTTCGCGTGAGTTGTGTGTGATTGGCAAGCTCCGTGCTGACATGTCGCCCACACCGTGGATGGCAAGCGGGCCTCTTCGCTGCCGCAATTTCTGCTGTTACCTGCGCTTGAAACAGTGCCGCTTCGCTATTCCTTCGAGCTTCGCGCTGTGAAGGCGCTCGTGTGCTTCAACCTTCATGTCGGAAACCTTGCCCGTCGCGGCAGCGCCGCATGCTGGTACCGCAATGGTCGGGCCTGAGGCGATCCGATGTCTCCTTTTTCTTCCGTGTCAAATTCGCGTTTGTCATGGGTATATGCGGCCCCGTGGATTACAGCGCAAATCGAAGCAGCGATGTGTGGCGCGTCGCGAGGCCCGATCTGCCGATAATCGTAGCTGTCTCAAGATTAAGCCGTAAATATTGTGGTATGAACTAGTGAAGCTGTGCTTCGCGAATGGCTCGATGCATCGATTATAATTTATTTGATAATTGCCACTACCGAGATAGCATGGCAATTCTAGAGCTGTCGTCTTGACAATCGCGACACCTGAAGAGACTGCATTTGTCAAATAAAAGGTCAGCGGTATGATGATTCATGATCTTCGGCAGATGCTGCGACATATCTTGGCGGCTGCCTCATTCGCTGCCTATCACCGATGTTGGGAATGGTGGCTTACCATGAAGTATTGACGATGACCGAGCGGATTGATCAGAGAAGAGCGAAGCGGCTCCGGAAAGCGGCGGTAGCAATCTGGCTGCGGAAGCTGATGACGATAAATGGCAATGCACAGCTGTGAGTCAGTAGTGGAGGTGAATAAATCCTTAGCGAGGTCCGATGGAGGCGAAATGCGTTGCCCGGCGACGTGACAATTCGACTCCGTGCGTATGTTGGGATTGTTGCCGATTAAGCCGTGAACAGGCAGCAGGTGAGTGCGACGATCGCCTATGTGGCTGCGATGCGTCCTGCGGCGATGCATCTGTGCTGTGGCGAGTGTCGCGTCGGTAAATTGCTCAACTGGCTCGCTAGATATTGGCGTAAGGATCTTTTTGTCATAGAGATTTCGGTCTATTGTTGGCGCTGAGACTCAGTGTGCGAGAGACGATCGAGAACGACAAAAATGGCCGTAGTCGGTGAAGTGCTAGTTTATGAAAACGAAAGCTATGGCTGCAGGGTCGGTGGGGCCGAGAGCGATCGATCCGTCCCCTGTCTGACCATAAGCGATCTTGAACAGGAAATCGGCGGATGTTACGCAGAACTTTCGTAGCTTTATGAAAATCAAGTTTTGGAACTAGAAGCAGGCTAACAAAGGCGGAAGGGCACCACCGAAGGCCCTGGCGACTTAATTTCGGCGGGGAAGCTTACCGGTCGGTATGACAAGCGGCGGCTCTTCATTACAGGTGGTGATACGTAACGTGCGGAGCAGTGAAGCTGTTTAATACGAGTTGTCGTTGCCCAACGGACCTCCGACGCTTTACTAACTATCGCGATGCTCCTCGCGGCGACTTCTTTAGGGGAGCATGGCGCTGGAAACAGGAAGACGGCGACGATCTGTGTTTCGCCGCTGATGTTACGGGCACGCGCGCTGATAGCGTTCGCAGTCGCGGCGGCGACGTGTCTTTAATTTCGTAAGTGATGAGAATGAATCATTGCAATTGTTGGTCTTCAACGAGGAATTCCTAATGGCAGTCCTCGACTGGCGTTATTTGCGTCCACGCCCTTTAGCTACCTGCCGCCACGTCGCTCTGCCGATGTGTTAGATAATTTTCGGACTCGGCGGCGACAACTTGCGATTACATTGGCTCTGCTGACGCTGATATCAGCCGGAATACTATCTGGAAGGAAGAAATCATAACAATTTCCGTAAAATGGCTTCTTGGAAGAGATCATTAATAGAAACACGCCTGATAAACTTGACGAGCTTGTGTTTCAAGCAGCTTGAGGCGAGGGCCTGCTCCTGCCTCCTCATGCGAGGGCGAAGCCTCGTCGTGCCAGCACTCACTGCCTCTCGGGCGTCTTCGCGTGGTGCGCAGGCCCGATGAGCTGCCGTCCCGAGCTGACTGGCTAAGTAATTAGATGCTGATATGTCTAAGCGACTCGCAAGCAGCGATTCTGTCTCCGACTCTGTCGATGGCGGCTGGCGAAAGCTCGACCTGCCCGATCCCCGTGAACCAGCATTTGAACAGTGCAAGCCGTGGCGAAATGCCGCAATCACGTCGTTACCTCGACCCCTCTGGGAAGTTAGAGTGGGAGCCGAGATGATATTCCGTCTCTTCCGTCGCCTGGTGGCATAAATAACGAGTCTCGGCGACCGGCCTTGTCCATTGTCAAACGCCGGTGTCCGTCGCGCCGCATCTTGGCTTTTCCATCTAATCTTGGCGATAAACCAAATCAACAGCTTTGTAGCGACCCAGTCGGCCAATTTACCAGCAATTAAGCATATCAATAAAGCGGAGAGAAACTTACAAGGATTCCCGCAACGGCGGCGGCAGGCGACCCGGAAATCTAATAAGCAATCAGTGAAGCGTCACCGGCGGCGGTAAAACAATCCTGGAAGAAGGCTTTGCGAAAGTGAAATGCGGACCTAATGCGCTGCAGCCTTCTTCGAAATCATTGTTTGGGAATGCGACTTATCAGCGACGTCCGATCCAAGCCGAGCGGCGATATGGCAATGCCGCGAAAAGATGAAAGAGACACGAAAAGAAGTCGAAGTGCTTAGTGTGAGGAGGAAGCCAGAGCGGCGATGCGCCCGGTCGGATGAAACGATGCAAGCCCGATAAACCCTTTCGACTTTCGAAGAAGCGTGTTCGATCAGTGCGGCAAAAGCAGGCTGTTGATATGCGCTGACGTCGGCGATCATAAACACTTGTATGCATCTCAGCGTGCTTCAGCATCTATGCTCTATGCAACTGTGTCGCCGTCTTGGCTACGGACAGAAGTACGACTCAATGCGGCTCAACAGCTGATCGCTCCATGGCGCAGGGGCTGTGTGCGGGATCCACCGAGTGGCTTAGGCCTTGATCTTCGAGAGGAACCGGTGTGTGCTGTCGGTTCCAGATGTGCCTTTTATTAATTCAGAAGCGAAGGAATACCGCAGAAAATGTTTATCTGGGCATAAACGCTCATCTGCGTTGACTATTGGCTTTAAGCTATACATTTGACATATATAAAGAGTGGCAAATAAACAATGTCATCGTTATCTTCAGCCTCTTGCGTGCGCAACACGGAGAAACAGGCGCACCGACCAGATATTGCTACATACGAGGCTGATGATGAATCTTTTGACACGTTCACTCAAACCGGTGTAAAAGTCGCTTTGGCAACTATTTATCGACACTACTGATAACGTGTTAAATGTCGTGGGAACTGCTTGACAAGCGAAGGAGGTATCTGTGAAGATGTGTGGTGTAGTGAGCGAAACACTGGGTAAATGTGTGCTTACCTCGGCAGTTACCATTAAAGCTGTTCTCGATAGGTGCCTCTGGCCCAATCTGACTGGAAGGTGATCGATGGTCTTCGTCATGCAGCGACCGTATACGTGGAGAATGCACAGCTAGTGTCGGTCATGCTTCCTGTTTCTGGCGATCGGAATCTTCTTCTTGCATCACTTTATCTGTGGTGCGCCTGCGAAGGAGCTCAGATGTGCGCAATGCTCTTCTTCTGTGCGTATGAAGAAACACTCCATTTGTCCCGATCTTGCTGCTAGTGCTCCTTGGAAAATTTGTTGCGTGTGCGGTGTGCTTCCGGTTCCAGCAGTTGCTGGTTTATACTTTATACTTCCTTTGTTACACGGAGTAATATAAACATCTATAAGCTACTTTAACGTAAATGATGTTAGGTATTATCCCTTCCCTTGCTTAAGCTAAATGAGCCATGAAAGTGAAATCGTCTCCTCGATAAGGCTGCTGGGTACTGCGTAATTGGGTGTTGCTGACTGGTCTCTGCAATTTCGGCTTCCGTAGGTTGCAAGTTGCCTTCATCGATAATTCGCCATTTCACCTTACGCATAAAAACGTAAAGATGCTGCGTCTTTGAAAGAAGGCGAACAAAGAACCTTGTGATGATGGTGGAGGCATTAACTTCAGTCTTTGCATCTCTACTCGGCAGGAGCGGCTGAGTGGGGTGTGAATCTTGCTGCACGGTCTGTATGGATGCATGTTTGCTTACTTTTTGTCCCTTAATTTTATCCCAGAACGTCTGTGACGCCTTGAATAAGCCCCTCTGGGGGTACATGTGTCTTCTTTGTGTCTACCACCTGCTATTATAGTTGGGTGTTGGGTGTCTCAGTGGTGAGAATACATTGGTTGTCGATTGTAATTGCATTTTCATTTGGAAGAACATGATTTGTATCACTGAATGATGCTTTTGGTCTGATTGCACACTTCTTCAAATCTGCAGCTTTGATGTGTAATAGGAGATGTTGCATCACCTTGGAAACATTACCTGCTTTATCCCATGATGGGTGGCATAACGAGGATCGATATTATTGAAGGATGGCGCAACCTCAGCGTTAGTGATTACTTTCAAGGGTGGCCAGTTGCAACTGCTTTGTTGCCAAACCTCCTTGTAGTTTGCCTTGATACATTTTGGTTAGGTATTCTTCTTCTCCTCTACAAAGATAAATACTGCGACAAAAGGCGTCGTGTGTCCGTGCATCTTCGCGAAGGAGGCATTAGGCAGGATTACCTTGGCGATCGGTTGCGGTATCCTTCCGGCTTCACCAATTTTGGATGAACAATTTACGTAGAATTGGTTTTATCAGTCTATCTCCACGCTGGCCGTGGATGCGGTTTTCATAATATTCGCTGTTTCGGATCGGCTTGATAATCGCGCAAAAGGTTCCCGATCGGTCACGAAGGATGGAATTCACATAAGCTCCAGGTTGCAAACACAGAGCCATGACCATTTGTAAACAATTGGTGATTCCGCCTTCTCCCAGGATTGCTTTGCCACTTGAAGGTGGTATCTCCGCTTTATGGAAATCTTCTTATGGTTCTTCGGAGGCGAATAACATGTCCTTCTCAGTTATGAATGGTCATTACTCTTCTAGAGTCTAGGGCATGTGACATTTTGTGGATATGCCCAAAGTCCGTGATGATACCTCGTATCGTCTCAGAGTTTCAGCGTACCCTCGTGAATGCATGATCGAAGGAGAATGCAAAAATTAGCCTTTGCCTTATCAAAATGGAAACTTGGCTTATATGTAATGCGAGCATACCTTGATAGTGTCATGATGACAGATCGAGAGATCTCAAATCGAGTAGCGGTGATCGTCTCTCGGTTTGTGATTGTTGCATACGTTAGGCTGCGTTGTAACAGAGCTATCTCTGCTCTCCCGGATTCATCCATGGATGCTTTTGAGATCAGGCCGAGAGTTGTGGCTCGAGACCCCAGAGTTCTGCCCGGCCCGGCTTGCGTCGCGAGTTGCTCACGCGTGGTAGTACTGCTTGGATTAGAAGCCTGCGGCGTACGGAGTTACTCCATCCAGGTTAACTCTTTGGGTGCCCATCTTTGAATCATGAAGGCATTTCTTGCAGCGAGTAAAGGGGAAGGCATGGCAGTGTGAAAACTTGACGTAAATCCTTGTGCAGCTGATTTGCCGCCCCATCCTTAGTGAAATTTTGGGAGTCCCAGGATGTGAAGGACTCTTTGATCATGGCAGTTCGGATCCAGTTTCTTGGAGGAAGGCATATCTGACCGCCGAGATCTCGCACATGCGGGCGGTCTGTACTGCTGTAACGAAAACCTGCCTTGACCTACCGTAGGCGGAGATCTGCTGTCAGGAGAGAGTCCTTGTTAGATCACGCCTCTGCCCGGGGCTTGATCGCCCGATCACTGGCACCTGCAGATCCGCCCAGGCTTGCCAGTTAGCAAGATCACAGACTCGCCCCGGCCCGTAGACCCCTCAGAATCGCCTCCACGGATCACCGTCTGCGAGTTTGTGCGTCGCGGCCGCGGGTGGAGGCCGATCCATGAGTCAAAAACGCCCCGAGATGATCCAAGGTAGCTTAGCACACGGATCCGACCTTGTGGGCTGCGGATTCTCTTTGGGCGTGGGTGGAGCCGGAGCCTCCGCCCGGAGGAGGAGGAGGGAGGAGGGGAGAGGCGAAGCCGGAGAACTTCACGGTGGAGGCCCGCGGCAATCTTGACGTGCAAATCGTTAAATCGACTTGGGTATAGGGGCGAAGACTGGCCCGAGCGAGTCAAGTATGGTGGTTCCTCGAAGTTTCCTCGGGATAAGTGGAGCCCAGTGAGCGAGTTCTATCGGGTAAAGCCAATGATTAAGAGGCCTTAGAGGTGCCTAAACCTATTCTCATTTTAAATAGGTAGGGCCAGTAGCATTTGCTTTGTTGAACCGCCATGGGACGAGAGCTCAAGTGGGCCATTTTGGTAGCGGTGGCGATGCGGGATGAGCAGGCCGGGTTACGGTGCTGACTGCGCGCCTAGAACCCACAAGGGTGTTGGTCGATTAAGGCAGGCGGAAGGTGGTCATGGAAGTGAAATCCAGCAAAGTGTGTGACTCACTGCCGAATCAGCTAGCCCCGGAAATGGATGGCGCAGCGCGCGACCTACACCGGCGGGTGAGGCAAGTGCAGAGCCCCATGATAGTGGGAGGCGGCCCAGTCATTGCCGAAACCTAGGCGTGAGCCGGCGGGCCGTCAGTGCGAGATATCTTAGTGGTAGTAGCAAATATTCAAATGAGGAACTTTGAAGGTGAAGAGGGGAAAGGTTCCATGTAGACGGCACGCCGCAGAGTTCGTAGTCCTAAAAGCTAGGTAACCCGTCGTGCGCGCGACTCGTGCGAACGAGGAATCGGGTTAAAATTCCTAGGCAGGACTATTGGTGGTTGGCCGAGAAAACGCGTTAGAATCGGAGGCGTCGGCGGGGCCTCGGGAAAGTTATCTTTTAAACATTTATGACACCTTACCACAGGCGGCTCGGTAAGGCGGGAGTCGGCCCGGGCGGAAGAGCACCGCACGTCGCGTGGTGTCAGGTGCACCCCGGCCCGGCCCTTGAAATCCGGAGCGAGGTGCCATTACGCCAGTCGTACTCACCTGCTGCTCAGTCTCCAAGGTGAGCGACCTCTGGTCGTGGAGCAGTATGGCAAGGGAAGTCGCGAAATGGATCAGTAACCCGGCCAAAAGGATTGGCTGCGAAATTTGGGCAGGGGTCCTAGTCGAACCCGTCGCTGTCATTGGCCCCACTCAGTACATACTGCGGCGAGGCGGGTGATCGCGCTGATGTCAGCCGAGACAGTGAGCGAGCTCTTTAGAGCTTTCCCCGGGCGTTGTGATCGACCAGGCCCGAGTGCGGACAAGGGAATCCATTTAATTAAAACAAAGCATTGCGATGGTCCCTGCGGATGTTAACGCAATGTGATTTCTGCCGATTGCTACGAATGTAAGTGAAGAGGAAATTCAGCCAAGCGCGGAGTAAGCGGCGAAGATAACTATGACTCTCTTAGATGGCCAAGTATATAAATCATCTAATTAGTGACGCGCATGAATAGGTGCGAGATTCCCCACTGTCCCTGTCTACCTCCGTGAAACCCATTGAGGAGCCGGGCTTGACGGAGATGTTGGGGAAAAGAAGACCTGTTAGAGCTTGACTCTAGTCAGAGCTTTGTGAAATGACTTGAGAGGTGTAGTATAAGTGGGAGCCTTGGGTAGTGAAATACCACTACTTTTAACGTTATTTCTTATTCCGTGAATCGGAGGCGAGGCCTTGCCCTCTTTTTGGACAAAAGCCATGCATTTGGCGCTGATGCAGAAGCGAAGACATTATCGGTGGGGGAGTTTAGTGGGGCGGCACATGCATTAAAAGATGCAGGATGTCTAAAAGATAGAATCCCGGCGAGAACGAAATCTCGTGTATGGAACAAAGGGTAAGCTCGTTTGATTCACGATTTCAGTCGCCCCCGAGGAATCGGCGAAGTAGCGTGGCCTATCGATCCTTTAGACCTTTGGAATTTAAAGCTAGAGTGTCGAGAAAAGTTACCTGAGATAAGCGGCTTGTGGCATTAGCGTTCATACCAGCGGCAATTGCTTTTTGATCCTTCGATGTCGGCTCTTCCTATCGTTGTAGTAGAATTCACCAAGTGTTGGATTGTTCACCCACCCGATAGGGACGTGGTACAGGTTTGGGCAGGTCGTGAGGCAGTTAGTTTTACCCTACGTGACGGTGTCGCAATGGTCAATTCAACCTGAGTACGAGAGGCGTTGATTCGCACAATTGTCCATCAGCTTGGTTGAAGCGATGAAAGCCCAGCAGCTCTACCGTGTCTTGAATTATGGCCCGAAACGCCTCTAAGTGAGAATCCGGGCTAGAAGTAACGCGTACGCCCACATCCAGATTTACCTGACCACGGTAAAGGGGCCCTAGGCCCAAAGGCACGTGTCGTTGAGTCGAGGCCCTCGCCGGGGACAAGGCTGAGGGCCGCCTGGTCTGATTTCTACGGCGGCGGGTAGAGATCCTTTGCAGACTGACTTAAATCGTGACGGGGCTATTATGAAGTGGCGAGTGGCCCTTCTTTGATCCACGAGATTCGACCTGCGTCGCCCGATTAATCCCTCGCCCATCCTTCCATTTTCCTCCTCCTCCCGAGCGGGACTAGCCTCCCAACACTTGGCCAACTTTGAAGTCGGCGGCGGTGCTTAGAGCGGCTTAGCCATATTTTCAGGGTTAACCCGACCGCGCGGCCCAGCTTTGGAAGCTCTCTGAACCCTTTGTCGCGGATCGATAAGCGGTGGCCTCAGCCATAGCCAAACACTTTCGGAGGTAAAACTTTAGGTAGAGTGTCTTATTGTCATGATAGCCAGTGGGGAAAATTTAAGAGAAAATAGAAACTTGTGATTTTGGCCCAAAAAAACGTGTGCTATAGCCCACATAATCAAGCGAAATCGGCATTTGAGACGTAACTCATAAACCGCACATGATTTTTTGAGTTCAAACCGGTTAATAGAGCTTATATCATCATGTCTGAGTGGGAAAATATGGTGAAAACAAAGACTTGCGTTTTTGGCCTTTAAAACCGTGTGCTATAACCCACAAATCAAGCCGAAATCGGCATTTGGATGGTAACTCATAAACCGCACATGATTTTGGTTCCGTCGAGTTAAAGAAGGCTTATATCATCATGTGCGAGTGGGAAAAAATGAGGTGAAAATAAGGGCTTGGTTTTTATTTAAAACACGTGTGCTACCTGAGCTGAGAGAAATCGACATTTTCCAATTTTAAATTTTTTTTTTTATTTTTTTTTTTTATCGTTTCTCAACTAATGATGATTATGAACATTTGTTTAGTGTGAATCTAAAGTTTCTAATATTTTGAGGAAAATTTATGTTTTTTCGATTTTTGTAAGAAAATAAGTGAGACCGTGCCAAATGGGCATGCCAACGTTTGTATATATATATATATATATAGAGACCGGTGTGTGCTTGAGGTAGCCTTAAGGGGAGGCTAAAGTATGGATGCCGAGGGGTTTGAGGTTGCACAGGTCTGAGAGCATGGATCAGGGAGACAGTGGAAGCTTGAGGTGTATAGGAGCTAAAGTATGGATCAAGGGAGTTTGAGGGTTGCCTAGGTCACAGGCATGGTTTGCCCGAGGTGCAAAGCATAGGCCTTGAGGATGCCGAGATCTTGTATAGCGTCACTACTTAGAAGTGCATGATGCGCGAGCGTAGTTTGCCAGTCTTTATGTGATCTTGTAATGGTCTTGAGAAATTTCTTAGTGGATGTCTTGGTCTCTTGTGACATGGTATGAAGTGCGGCGGAACAGTCAAGGCTGAGGGGCAAGTGGATGCCGCGGGTGGGCGTAGTGAGAGGCTACGCGTCCCCTTGTCGTGCCGAGTCTTTGTGATCACCTTGTGAGGCTGAGAGCTGCCGAGTCTGTAACATAGTGTGAGGGCGGCCGCAGTGCGAGCCAAGGTTGCGGCAGATTCTTATGAAAGTGTCTTGAGAATGTTTGCCCAGTCTTTGCCTTGATGCCCGAGCGAGGAGCTGCGAGGATGCGCTGGTCTGCTGACATAGGTGCGGTGCCAGCAGTCTCGTGCGAGCCAAAGGTGCGTCGGGTTGCGCGGTGTGCGGCATGGGTGTAGAGTCTTCTTGTAATCACCGCGGCAATTTGCGAGATGCGCAGTGCGCGACCCGAGGGTGCCCGAGGGGGATGCCCAGCAGTCTTGCTTGGCCCAGGGGCTGCGAGTCTTTGCGAGTCTCTTAGGAGTGCATCGCGATCACGTGACCTTGCTGAGGGGCTGCTTAGGGGAGTTCCAAATTCTGATTTTCACTTTTTGTAGAGCATGGGTGAACTCCGTGCCAAATGGGCATGCCTGTGATTCCATATAGTAGGGGTTGGTGGCATGGAGCAGGATGTGGAATGGTTTGGTGGCATATGGTTGTTCACGAGTAAGGGGCAAGTTATTTCCATAGATATGGTTCTCGGGTTTGTTTTCTTTGGATGCTCAGGCGCATTGTGCTCTAATGTTCATTCTCGGTGAGTGAGTCGGAGCGTGGCAGTGTCCAATGACGAGTCTGCTGGTTCTGTAGAGTTTTGCCTCCGTTGATCTTAACGACAACACGCCTGCGAAAAGTTGAACAAAGAGTCGGTTTTGCTCATGAGGCCTATTGGGTGAGTGCTTTTCTATGTTTTCCATGTTGGCGGTGCGGGCTTGGCTACATAAAATCGTTTGCGTTGGCAGCCATTACATTGGGTCGGGCGTGGTCGGTTTAATGTTCTTGGAGTCTGTGATATAGCAGCGTGGTGAGTTGTGAATCGGTTGTGGGTTGGCGGGCTCCGTGCCCGGCGCTCGAACCGTCGCCCACACACCACGTCGATGTGAACTTTCTTGTGCTATACGGAGATTCTGTGTTGCACCTACGCTTAAAGGAGCAGTCTTTGCTCGCTACCCTTTCCTTCCCAGGAAGCTTCAGCTGTGGGAGGACATCGTAGTAAGCGCTGTGCCTCGCATCGAGCAACGCCGTAAAGCGCGCCGGTGCGGCCAGTTCTCAGGTTGCTACTTGATGGTTCGGATCGGAGACATTCGGTGGGCAGGTGTCTCGGCACCTTTGATGTCAAATTCAGCGTTTTGTCGCTGGGCATATGCGGCTTGATTACGTGACTGCGGAGTCTCCAGTAGCGAGTGCGTAATCATTCGGCAAAAGGAATACTTCACGGTTGATCATTGCCCCGGTAGTCATATGCTTGTCTCAAAGATTAAGCCATGCATGTGTAAGTATGAACTAATTGTTGTAAGCTGCGAATAGCTCATTAAATCGATTATAGTTTGTTTGATGGTACCTACTATAGGATAGCATTAGTAATTCTAGAGCTAATACGTCTGACAAATCACGACTTACGGAAGGGATGCATTTATTGGATAAAGATTGTGAGCTCTGCCGTTGTGATGATTCATGATAACTCGGCGGATACATGCAGCCTTAGTGCGCGCGCATCGTTCAAATATACGCCTCTCTGACTTTCGATGGTAGGATAGTAGCTACCATGGTGGTAGTGACGGAGAATTAGGGTTCGATTCGGAGGGGCACAGAAGCGGCTACCACATCCAAGGCGTGGTAGCGCCGTTACCCAATCACGACGGGGAAGTGTGTGATAAATAACAATGGGCTCTATGATGCAGTAATTGGAATAGTACAATCTAAATCCTTAACGAGGATCCATTGGAGGGCAAGTACGGTGCAGTGACCCTTTGACAATTCTAAACTCCAATAGCGTATATTTAAGTTGTTGCGAGTTAAAAAAGCTCGTAGTTGGACCTTGGGATTCAGGTAATGGTCAGCATGGTGTCTGCAGGTGGCTCGTCCTCTACGGCGATACGCTCCACAGCCTTAATTGGTAGGGTCGTGCCTCAGTCTTCATTACTTTGAAGAAATTAGAGTGCTCAAGCAAGCCTACGCTGCGAGATACGTTAGCGTGGGATAACGTCGCCAGGATTCGGTCTGATACTGCCTTCGGGATCGGAGTAATGATTAACGGGGCCATTCAGAGGCATTCGTATTTCATAGTCGAAGGTGAAATTCTTGGGTTACTGAAGGCGAACAAGCTATGAAAAGCATTTTGCCAAGGATGTTTTCATTAATCCAAGAGCGAAAGTTGGGGGCTCGAAGACGATCGAGGATACCGTCCTAGTCTCGACCATAAGCAGTGGGCGGGGATCGGCGGATGTTACTTTTAGGACTCATGGCACATATAGAAATCAAAGGAAGTTTTGGGTTCGGGGAAGTATGGTCGCAAGGCTGAAACCCAGGTGTGAAGGAGCACCACCGGGAGTGGAGCACAAGCAATTTGACTCAACACGGGGAAACTTGAGTCTAAGACATAGTAAGGATTGTGGGATGAGAGCTCTTTCATGATTCTATGGGTGGTGGTGCATGGCCGTTCTTAGTTGGTGGGCGATTTGTCCCGGTTAATTAGTTAGCGAGCGAGACCTCGACACCTTCTAACTAAATTTATCACGGAGAGTCTCCTCGCGGCCGGCTTCTTCAGAGGACTATGGCCGCTTAGGCGAGAGTTTGAGGCAATAGCAGTGTGATGCCCTTAGATGTTACAGGCCGCCGCCTTGTACCACTTTGATGTGTTAGCAGGTCTATAGCCTTAGGTAAACGGGCCGGGTAATCTTTGAATTTCGCGTGATGGGGATAGATCATTACAATTGTTGGTCTTCAGGAAATTCCTAAAGCGCGAGATCATGTTTCGCGTTGACTCACGTCCACTGCCCTTTGTACACCGCCCGTCGCTCCTCTGGATTGAATGGTCGGTGAAGTTTTCGGATCGCGGCGGCGTGGCCGGTTGACATCTGCGACGTCGCGAGAAGTCCTTGAACCTTATCATTTAGAGGAAGGAAGTCGCTAAGGTTTTCGTGAGTGAACTGCGGAAGGATCATTATCGAAACTGCCCCAGGCGAAGGCGTGTTTCAGCGCTTGGAGGCAGGGGGCCTCGCGGCTCCTCGCCTCCTCATCCCGAGAGGCGAGGCCTCGCGTGTCGTGCTTCGGCGCTTCGCTTGGCCAGCCTCTCGGGCGTGCCCGAACATCGGCGTGGGTGCGCCAGGAACTTGAATGAAGGGCGTCGCCCGCGTCCCGGAGGCGGTGGTGCCTTGGAGTGGTTAAATCGTCTTGATATGTCTAAGCGACTCTCGGCGGATATATCTTGGCTCTCGCATCGATGAAACGTGGCGAAATCGATACTTGGTGTAGTTACGGAATCCAGTGAACCGTGGGTTTTGAGCGCAGTTGCGCAAAGCCATTAGGCGAGGCACGTGCACTGGCGTCACACGTCGTTGCCCCTCCCAACCCCTCTGGGAGTTGGGTGGGACGGATATTGGCCTCCGTGCGCTCGTCGCCGGTTAGCATAAATAACAAGTCCTCGCGACCCAGCGCCGTATGATCGGTGGTTGTCAAACCTCGGTGTCCGTGTAGCGTCATCGGGCTTTTCCCATCTGTCTGGCGTCAGTCGGTAACTTTCAGCCAGCGACCCGAATTGACCAGGGTTACCGTGAATTTAAGCATATCAATAAGCGGAAGAAACACAAGGATTCCCTTAATGTAGTAGTAAGCAAGAACATTGACTTGAGAATCGCGTCGCAAAGGCGTCCGGAGTCGCTGAAGCGCCCTCGGCGGCGGATCGGAACAAAGTCCTGGCGCCGCGAGGAGTGAGCCGTTGTGCTGACCTATCGCTGCAAGTGGCCCAACCAGTGGCTCAGGTTGTTTGGGAATCTGAAGCCCCAATCAGGCGGTAAATTCCGTCCAAGGCTAAATACGGGCGGGGGCCGATAGCAAACAAGTCACAGCGAGAAAGATGAGAAAAGGACTTTGAAAAAGAGAGTGAAAGAGTGCTTGGAAATTATGCGGGAGGGCGGATGGGGGGCAGCGTGCGCCCGGTCGGATGTGCACAGACAGTCCTGGCCGATCCGCCACTCGGCTCAGGCGTGGACATTCTGGATTGCGCGCGTCCAGCAGAAATTTGATGATATGCCGCGGAGGCGTAAATCGGCAACGATCGTAGTAGTGGGCGTGGCCTTATCTCAGCGTCTTCGGCATCTGCGTGCTCCCAGTAGCGGCACATGGGCTCCCCATTGGCCCGTCTTGAAACACGGACCAAAAGTCGGCATGTGTGCGAGTCAGGCGGTTAAACCGTAAGCGTAAGGAAGCTAATGTGCGGGATCCCTTTTCGGGTGCACCGCGTAGGCCTTGATCTTGAGAAGGGTTAGTGTGAGCATGCACAGCCGGGACCGAAGATGGTGAACTATGCCTTTATTAATTTAATAACCCAGTGATACAATGTGCTCTCCACCAGATGTGGATGCCAAATGCAAAGAAAAAAAAAAAGCGGTGCAATACGGTCCTACGAAAACTAGAGGCTCTCGAAGTCTAGCTTGGCGCAATGTTGCATTTCTAGGTAGGTGAATTCATCCCTTCTTTCTTTAATCCTCGAAGCAAAAGTGCTTTGGCTCCCATTACGATATCCGTGATCATTGGTTTTCTCTTCATCTGCCGTAGTGTCGAAGCCTGCCGCCCGGAATGATGTTCCTCATTTGCTTGTCTTTGTTTGCTCTTGGAGGACTGGGTGTGAAGAGTTGCTGCAGTACATCAGGCCTTTAATTGCTAAGTCCTCTTTGGAAGATGCATTTGCTGATGCAGTGGCATCTCAATTATTACGGGTTGTGTACTCTTGTTTTTCCTCCTTTTAGTGACAAGTGAATAACCAGCCTCGCCGATAACTTCCTTTCTTGCATCATCATTAATGCTCTCCAAGGTAACATCTTGATAACCTCATCTAGAAGCAAACTGTTAGAGGTTTGCAACATTTGGTGGTTGCATAGAAGGTATGGTGGGTGATCCGTTGTAGCATTTAATTTGGTTGCCAGTCGGCTCATTGGATGCTCCAAAGCATCTATGGTAAATTGCTCTGGAAGTGTAGAAGGAGGTGTAGTCTTCGGCTTCCAAACCATTCGCGACTTTGATGCACCGTTGTTAGTTGCGTATGCTTCTCTTCTGCCAAATACTAAATTTTAGACAAGAGAGGAGTTGTGTGTCACATGCATTTTGCGGAAAGAACAATAAGTTGGGTTCCCTTCAAAAGCAATTGCTGCCAATAACCATGCATCGCCTATACATGAAAACCTTTGTGGGAAGTGGGTTAGAGGCGTCGAAATCTCAGCACAAATCCAGGATGTGCGGGAAGCGAGGTGTTTGTAGTCTCGGGATCGATCTCGACAAATCTCAGGTAAAGGAATTACCAATGAGTGAAGGATTGTCAGGACATAAGCTCGGAGTTGTAAACACAGTGCTTATCAGCCACTTGTTTATAATGGGTCTGCCTTTCTTCAGGCTACTTTGCCATTGACAAAAGGTAGATGCCCTTGTAAACCATCCATGGTTCCTTGAGTAAGCGGCGTGAGGTCTTGACATTCATGTTCAATTAGAATTCTTAGAGTCAGAACTCCCCCTTTGCTGTTGGCTTAGAAGTCCCCACACCCCATGTATGATACCTCGAATTGCTTCCCGGCTTGACCTTACCCAGTGAGCGCGCCAATCAAGAGTGTAGGCTAAATTAGCTGCACTTTCTCATGATCGGCCCGGTGCTTATGTGAATGGGTAGCGTACCTTGAAAATGCTCCTTGTGCAAAACCCTCGATCCAAATTAACTGAAGGAATGAGTGTAGGTTTTATGTGATGGCATGAATCAGGTTCTTCAATGCGAGAGGCTGCTCTGCAGCCCGTGTGCTCTCCTCTCTGTTTGATACTGACTGAGCAGCCGTTTGAGTATGGATATTCCAATGAGACGACGAGTGTCGAGCTCGGATGCGAGGTTAGTCTTGGAGTGTAGCTTTCTTGATCGGTGAGGAAATTGATATGCGATCCCTCAGTGTCTCTACATTGCAGCCAAAATATTCGAACTCAAGTTTGTAACCTTGAGTGGATTCGATCGGATTCAAGTTTGTAACCTTGATTGATTCAATAAGCGCCCTTGAGCTCGAGATGGAGTCTGTAAAATCGGGATTGGGATAGCAATGTTGCTCAAATTTGAAAGCAGTTGCATGCGAATCACTAACCCCCTGTCCCGGCCCAGACAGGGGGCACTCACGTAACTTCCATGATTCAGAATCGGGCGCCATGGTGATGAAGATTGAAGCTCTCGAGGCATGAACATCATGAGCGTTTCCACCTCCCTGCACCTTGAACCGGCGAGGAAAGGCCTTGAATCCTTCTCCTCCGTGATGTTTGTTGCCATTAAGCTAGGAGTATACATATAGCATTCCGCATCTCTAGTCACCGGCTCATGAAATTAATTGCAAATCTCTCCACCAAAACATTACGAATGAAATTACCTCACCTTTTTAAAGACAAATCGCTGCGATTCCCAAGTTGGCGTAAGCGCGGTAGATCGCATTCCCGATGAATAACATTACTCCTCCCCAGGCTTGCCCCTTCTCAATCGACTTGATCATGCGGCGTGACTTCTCCCCGCCGTGATCTCCTCAATGGGCCAGGCGCAGTGTCAGAACCTCCCGTGATCGTGCCTTCTCCTTGGCCAAAATGGAGGAAGCCTAATGGCCTCCATCGAAGGGAGGAAGGAGGAAAGGAGGAGGAGGAGGAGGGAGCTACGGTGGAGGCCACGGCGTCTTGGCGTACCAATCGTTCATCGACTTGGTATAGGGCGAAGACTAATCAAGGCAGTCTGGTAAGTTTGGTTCCCTCCGAGTTTCCTCGAGATAGGTGGGCCCAGTGAGCGAGTTCTATCGGGTAAAGCCAATGATTAGAGGCATCGAGGCTCAACGCCTCGACCTATTCTCAAACTTTAAATAGTAGGGCGAAACGCTTGGTACTTTGTTGAGCATATAGGTCAAAGAGAGCTCAAAGTGGGCCATTTTGGTAGCGGGCCCCAGCGTGCGGATGAACGGGCCGGGTTACGGTGCCCGACTCTGCTAACCTAGAACCCACGAGGTGTTGGTCGATTAAGGCAATGGGGCGGTGGTCACCAGTGAAATCCAGCCTAAGTGTGTGACTCACTGCCCGAATCAACTATTGAAAATGGATGGCGCTTAAGCGCGTGACCTACACCCCAGCCGTCGGAGGCAGTGCAGAGCCTAGATGAGTAGGAGGCCTTTGGCCAGTGTGTGCAAAACACAGGGCGTGGGCCGGGGCGGGATGGCCGTCGGTCTGGTCTTGATGGTAATGTGAAATATTCAAATGAACTTTGAAGGCCGGGGGGAAAGGTTCCATGTAGGCAGCACTTACGTAGGTTGAAACGATCACTAAAAGAGTGGGTGCCGATACGATAGCGCGACTGTGTGAACCGAAAAGGAATCGGGTTAAAATTCTGAGCCAAGGTGTGGTGGTTGGCCTTGCGTTAGGAGTCGGAGGCGTCGGCGGGGCCTCGGAAGAGTTATCTTTCTGTTAACGGCCTGCCCCACTGCAGGGCAGGCTCAGTGAGGCCAGATCGGTGGGGAAGGCACATGCAATCGCCGTGGTGTCGGTGCGCCCGGCGGCCCTTGAAAATCAGGAGAATGAGAGTGCCATTCACGCCGTGTTACTCTGCATAACACATGGTCTCAAAGGTGGCTTGACCTCTGGTGATGAGCAATCTTAGGCAAGGGAAGTGGCAAAAAATGGATCAAGTAACCCGGGGAAAGGATTGGCTCGAGAGTGCAGGCACGGGGTCCCAGTCTAGAACCCGTCACCATCAATGGCTTACTCATTGCTCCATGGCGGAGCGGAGTGCTGTATCAACGAGGCTGGTGGAGCATCACAGGCTCATTAGATCGACTGGCTAGTGCAGACAAGGGAATCATTGTTTAATTAAAACAAAGCATTGCGATGGTCCTGCGGATGTTGATGTGATTTCTGCCAGTGCTCTAGATGTCAAGTGAAGAAATTCAGCAGCGGAGTAAGCGGCGGGAGTGCTATGACTCTCTTAAGGTAGCGAATGCCTCGTCATCTAATTAGTGACGCGCATGAATGGATTAGCAGATTCCCCCTTTGTCCTGTCTACTATCAGTGAAACCCACGGCGAGGAGCGAGCTTGTGAGAATCGTGGGGGAAAGAGACCACTGACCAGGCTTGACTCTAGTCCGACTTTGTGAAATGACTTGAGAGGTGTAGTATAAGTGGGAGCCTTCGGGCGAGGTAGTACACTACTTTTAACGTTATTTTACTTATTAGTGAATCGGAGGCGGGAACCTTGCCCCTCTTTGGACAAAGCCCGCTTGGTATTTGATCCAGGCGGAAGACATTGTCGAGGTGGGGAGTTTGTTTGAGGCGGCACATACATTAAGATACTTGTGGTGTCCTAAGATGAGCTCAACGAAACCCAATCTCGTGTGGAACAGGGTAAAAGCTAGATTTTGATTCGATTTCCGATACGAATACGAACCGTGGCGTGGCTATCGATCCTTTAGACCTTTGGGAATTTAAAGCTAGAGGTGTCGAGAAAAATTTACACGGGGATAGCGGCTTGTAGCGGCCAAGCATTCATAGCGGCGTTGCTTTTTGATCCTTCGATGTCGGCTCTTCCTATCGTGTAGACGGGAATTCACCAAGTGCATTGGATTGTTCACCCACCCGATAGGAGCGTGGTTTGAGTTTAGGCGGAAGTAGTGAGGCAGGTTAGTTTACCCTCTTTTGATAGCGATTATCACGATATTAATTCAACCTGAAATACGAGAGGAACCGTTGATTAGCACAATTGGTCATCGCTTGGTTGAAAACCCGATGTGCAGCGAAACTCGTCTTGTTGGATTATGGCTTGACGCCTCTAAGTCGAGTCGGGCTGACGGCCTTGTCGCCGCCGTCGGGCCGCGGTGGGGGCCCTGGGCCCCTGAGGCCGCGTGTCGTTGGTCAAGCCCTCGGCCGGGGATGAGCTGCGAGAACGCCTTGAAATTACAATTTCTGGCGGCTGGGCAAATCCTTTGCAGACCGACTTAAATACGCGGCGGGGTATTGTAAGTGGCGGAGTAGCCTTCTTTGCCCACGATCCCTTGAGATTCGATAGCATGCCAAATTCGTCCTCCCCCTTCCCATCCTTCCATTTTCCCTCCTCTCCTCCCGGCGAGGCTATTGCCTCAACACTTGGCCAACTTTGAAGTCGGCGATTGCTTAGGCAACAGCGTGTTTTGCGAGAGTTAACCAGCGCGCGGCCTTTGCGACTTCTGAACCCTTTGTCGCGAGGGGATCGTGTGGTGGCCTCAACACTTGGCAAACTTACTTTGAGAGGTAAAACTTTAGAAGTAGGGTGTCTTATTGTCATGATAAGCGAGTGGGAAAATTTGAAGAAAATAAAACTTACATGATTTTGGCCCCCAAAAACCGTGTGCTATAGCCCATGTCAATGAAAATCGGCATTTGGTGTGCTCATAAACACACTGATTTTGAGTTCCGTCGATTAATAAAGACTTATATCATCATCTAGTGGGAAAAATATGAGGTAGAAACAAAGACTTGCATTTTGGCCCAAAAACCCCGTGTGCTATAACTACCAAAAGGAAATCGAGCGAAATCGGCATTTCGGGCGTAACTCTCGCCGCGCATGATTTTGAGTTCCGTCGGTTAAAGAGACTTATATCATCATGTCGAGTGGGAAAATATGAGGTGAGAAATAGGAGCTTGCGTTTTTGGCCCCAAAAGCCGTGTGCTATAACCCGCAATCGGCGAAATCGGCATTTTAGACGTAACTCCTCGACCTTACATGATTTTTGAGTTAAATAGAGTTAAAGAGGCTTATATATCATCGTATGCGAGTGGAGGAAAAAATGAGGTGAAAATAAAGACTTGCGTTTTTGGCCCAAAAGCGTGTCTTTATAACATGAACTCGAGAGAAATCAGCGTTTCAATTTTTCAATTTTTTTTTTTTTTTTATAGTTTCTCAACTTTAATGATGATTATGAACATTGTTTAGTGTGAATCTAAAGTTTCTAATGTTACCGAGGAAAATTTATGTTTTAGGTTTTGTGGAAAATAGGTGAACCCTAGTGCCAAATGGGCATGCCAACATTTGTATATATATATATATATATATATATAAGGGGCAGTGTGCTTCAAAGGTAGCCTTCGGGGGAGGGCTAAAGTGAATGCCGAGGGTTTGAGGGGTTGCACAGGTGAGGCATGGACTAGGGGACGAGTGGGCTTGAGGTAGCCTTCGAGGAGGCTAAAGTATGGATCTTAGGGGTTTGAGGGGTTGCTAGGTGCAAACTTGTTCTTAGGAGTGCAAAGCATGGTACATGAGGATGCGAGATGCGCGCAACATCTTTGCCAGAGAAATTTGCCATGGTCTTGACGTGGTGCCCAGTCTTTGTCGCGATGTAATGTGAGTGCGAAGCCAGTGGATCTTGTGGTCTTGCGGCATGTTGCGGGAGTGCGGTCTTGAGAGTCAAAGTGCGAGGGTTTGCCAGTGGATGCGCGGTGGCGTGAGGCTGTAGGAGATCTGCGCCCCGTCTGTATCGTGCCAGTCTTTGCCGCGATCTGTGTGAGGAGCTGCGAAGAAGTGCCAGTCTGCGGCATGGGCTGCGAGGGGCTGTAGTCTGAGCCAAGGTTGCGGCAGGTTGCCGCGGTGTCTGGAATAATTGCCAGTCTTTGCCCTTGAAGCCAGATAAAGAATTTCTCGAGGATGCCGCAGTCTTGACATGTTCTGTTTACCCAGCAGTGCGAGCCAAAGTGCGTCGGGTTACCTTGATGTGCGGCATAGGCTGCTTAAGTCTTGCCGCGATCACGCGGCAAGGGCTGCGAGGATGCCGCGGTCGCCTTGTGGCTGCGAGGGATGCCACGCGGTGCGCGTGAGAATTTCGCGAGGAGTCTTTGCTTGTGGTCTGTAGAGTCTTTGTCGCGATCTTGTGACCTTGTGCGAGGGTGCTTAGAGGATTCCAAATTCATGATTTTCACTTTTTGTGAAACATGGGTGAACTCCGTGCAAATGGGCATGCCACAATTCCATATAGTAGAAGTGGTAGCTGGAGCAGGATGTGGGAATGGTTTAATTTTGGCATGTGGTTGTTCTACCAAGTAAGGGCAAGTTATTTCCATAGATATGGTTCTCGGAGTTTGTTCAAGCTGGATATCAGGCGCATTGTGCTCTAATGTTCATTCTCGGTGAGTGAAATTCGGAGCGTGGTGTCCAACGCAGCCAGAGTCTGCTGGTTCTGTGGGGGTTTGCCCTCAGTGGATCTTAACGACACTTCCTCGAAAAGTTGAGCAGATCGGTTTTGCTCGCAAGGCCTATTGGGTGGAGTGCTTTTCTATGTTTTCCATCTTTGGGCGGTGCGGGCTTGGCTATAAAATCGTTTGCGTTATGCGGCTTCATTAGCTCGGGCGTGGTCAATTTAATATTCTTGGAGTCTTTTGATGTGACGTGTGAGTTGTGGGCTCAGTTGTGTGGAGGTTGGCCCCAGACTCCGTGCTTCGGCATCGAACAATCGCCACCACCACGTCCCGATGTGACAAGACTTTCTTGTGCTATACGGAATTCTGTATTTGCTACCTCGGCAAAGGCAGTGCGCTGCGCTACCCTTTCCCCTTCCCAGGAGGCTTGCGCTGGCAGGAGGACATCGTGTGGCGCTGTGCCCTGCATCGAGCACGCCGTAAAGCGCGCCGCGTTCTCGGCAGTTCTCGAGTTGCTACTTGATGGTTGGATACGGGAACATTCGGTGGGCTTAGGTGTCGAGCACCTTTCGATGTCCAAATTCAAGCGTTTTGTCGCTAGACATGTCGTGTAATTCATTGCGGGTCTCATTGTGGTGCGTGACGTCGACAAAGGAATGCTACGGTTGATCACGCCGGTAGTCATATGCTTGTCTCAAAGATTAAGCCATGCATGTGTAAGTGAACTAGTCGGGTGTGAAGCTGAATGGCTCATTAAATCGATTATAGTTTGTTTGATGGTACCTCTGCTCGGATAACATTAGTATTTCTAGAGCTAATCGTACAGCAAATCCACGACTTGGAAGGGATGCATTTATTAGATAAAGGTCATGCGGAGCTCTGCCGTTGTATTGATGATTCATGATAACTCGACGGATCGCACGGCCTTAGTGTTTGGCGACGCATCATTCAAATATACTTCTCGACTTTTGATGGTAGGATAGTGGCCTACTGTGGTGGTGGTGTGGAGGTGGGAGTTCGATTCCGGAGGGGCACGAAAAGCTTGACTACCACATCCAAAGGAAGAGAGAGACGTGAGGGCGCGCAAATTACCCAATCACGACCTTGCGGGAGGTAGTGACAATAAATAACAATACAAGACTCTATGAGTAAGTAATTGGAATAAGTCAAATCTAAATCCCTTAACGAGAGGATCCATTGGAGGCAAGTCTGGTGCAGTGACGCGGTAATTCCCGGCTCAATAAGCGTATATTTAAGTTATTTGCGGTTAAAAAGCTCGTAGTTGGACCTTGGGTTGGGTCGGCAGTCCGCCTATGGTGTGCACCGGTCGGCTCGTCCTTCTACGGCGATACGCTCCTGGCCTTAATTGGCGGGTCGTGCCTCGGTCTTTGTTACTTTGAAGAAATTAGAGTGCTCAAAGCAAGCCTCGCTGCGGATACATTAGCATGAGGACAACATCATAGGATTTCGGTCCTGTCTGGCCTTGAAGGATCGGAAGTAATGACAATTGCGAGGGACAGGTCGGGGCATTCGTATTTAAAAATAGTGAGAGGTGAAATTCTTGGATTTATGAAGACGAACAAGCTGAAAGCATTTGCCAAGGATGTTTTCATTAATCAGAGCGAAAGTTGGGGGCTCGAGGCGATGGGATACGAATCCTAGTCTCGACCATAAACGATGCCGACGAGGGATCGGCGGATGTGCCCCAGGACTCGCCGGCACCTTATGAGAAATCAAGGATTTTGAGTTAGGGAAATATGGTCACAAGGCAGAAACTTAAGGGTGTGGGGCATACCGGGAGTGGAGCACACGGCTTAATTTGACTCAACAGGGGAAACTTGCGGAGTCCAGGACATAGTAAGGATTGTGATGAGAGCTCTTTCATGATTCTATGGGTGGTGGTGTAACCGTTCTTAGTTGGTGGGCGTTTATCTGGTTAATTTCGTTGACGAGACCTGTTTGCTAACTAGCTATGCGGAGGTCTCCTCCGCGACTTGACTTCTTGAGACTATGGCCGCTTAGGCCAAGAGAGTTTGAGGCAATAGCAGTGCAGCCGTGCCCTTGAATGTTATTACGGGCGCACGCGCTCTCTGTGATGTATTCAACGAGTCTATAGCCTTGGCGAAGCCAGGCCGGGTAATCTTTGAAATTTCATCGTGATGGGGATAGATCATTGCAATTGTTGGTCTTCAACGAGGAATTCCTAGTAGCCTTGAGTCATCATTTCGCGGTGACTACGTCCGCCCCTTTGTACACCTGTAAATCGCTCCTCACAGGTTGAATGAGTCCGGTGGAAGTTTTCGGATCCCGCGGCTGACGTGGCGGTTATTGTTGTTTGCGGCGTCGCGAGAAGTCCACTTTGAACCTTATCATTTAGAGGAAGGAGAAGTCGCCAACAAGGTTTCCGTGGGTGAACTGCGGAAGGATCGTTGTCGAAACACGCCTGTGAGAGTGACGAAGGCGTGTTTCAGCGCTTGGAGGCGGGGCCTCGCGGCTCCTCGCCTCCTCTCCAAGACGAAGCCTCGCGTGTCGTGCTTGGCGCTCCCGCTGGCCGACCTCTCCCGGGCGTGCCCGAACATCGGCGTGAATTGCGCAGGAACTTGAATGAAGGACGTCACCGCCGTCCCGGAGACGGTATTTGCTGTGGTGGTTAAATCTCGTCTTCAATGTCGCGACTCGTGGCGGATATCTCGGCTCTCATCGATGAAGGGCGTGGCGAAATGCGATACTTGGTGTAGATTGCAGAATCCGTGAACCATCGAGTTTTTGAGCATAGAGATTGCGCCTGAAGCCATTAGGCCCGAGGCACGTCTGCTAGGCGTCACACGCCGATTGCCCACTCCCGGCCTGCAGGAGTTGGGTGGGGCGGATGGCCTCCGTGCGCTCCGTCGCGCGGTTGCCCATAAATAACAAGTCTCGGCGACCCGGCGCCGCGACAATCGGTGGTTGTCGAGCCTTGGTGTCCCGTCGCGCGCGCGTGGTCGGGCTTTTCCCATCTAATGCGCGTCGCCGTCGGGCGCTTTCAGCGTGACCAGGTGGTAGGGTTACCCGTGAATTTAAGCATGTCGATAAAGCGGAGGAAAAAAGAAGCTTGCAAGGATTCCTTAATGTGACGAGGCAAGAGCAGCCCGGCTTGAAATCTGCATCATAAAGAAGCTGTCCGGTATGGCGTCACAGAGAAACGTCCTCGGCGAGCGCGGGCCAGGGAAGCAGAAGTCACACTGGGAGGGCGCCCACGAGGAAGAGGAGTGAGAGCCCGTTGTGCCCGGACCCTTCACTTTCCACGGGCGCGCCGCGAGCCGGGTTATTTGGGAATGCAGCGATCGGGCGGTAAATTCCGTCCAAGGCTAAATACGGGCGAGACCGATAGCAAGACAAATACGTGAGAAAGATGAAAAGGACTTTGAAAGAGAGTCGAAAGAGTGCTTGAAATTGTCGGGAGAGGCGGATGAGGGCGCGATGCGCCCCGGTCGGATGTGGGCAGTCCTGTGGCTTGATCGCCCACTCGACTCGGCGTGGACCAGGTGCAGTTGCGGCCCCAATGCCGGGGCTGTTGATATGCCGCGGAAGTCGTCGGCAACGATCATGGTGGCGGCCACGCGCCGTCTCGGCGTGCTTGGCATCTGCGTGCTCTAGTAACGGCTTTGTGGGGAGTTTGTTGGCGGCACATCATTAAAAGATAACGCAGGTGTCCTAAGATGAGCTCAACAACGAAATCTAGTGTGGAACAAAGGGTGAAAGCTCGTTTGATTACGATTTCAGTCTGAATACGAACCGTGAAAAGCGTGGCCTATCGATCCTTTAGACCTTTGGAATTTTGAAGCTAGAGTGTGAAAGTTACCACGAGATAAGCGGCTTGCTTAAGCGACCAGCGTTCATAAGCGGCGTTGCTTTTGATCGCCGATGGCTCTTCCTATCATTGTAGACGAAATTCACCAAGTGTTGGATTGTTCACCTGACCAGGAACGTGGTGGAGAGTTGGAGCCGTGCGTAAGCCGAGTTGGTTTTACCCTCACGTGTGATTAGCCATGATAGTGGTGCCAATTCAACCTGGTCTGAGAAGGTAGTTGATTCGCACGATTGGTCATCGCGCGGTTGAAAGCGATGTGTGGCTGCCGTGCGGCTGGATTATGAGCGAAACGCCTCTAAGTGAGAATCCGGGCTAGAAGCGGCATGTCGCCATCGTCCGTTTGCCCGACCCGCGGTAGGGGCCCTAGGCCCCGAGCGCGTGTCGTTGGTCAAGCCCTCATGAGGACAGGCTGCGAGGCGCATGAGATCCACAATTTCCTCACAGGCGGCTGAGGTAGAATCCTTTACCCACGATCGACTTAAATACGCGACGGGTGTATAAGTGGCGAGTGGCCTTGCTTGCCACGATCCTTTGAGATCGGCCTCAAGCCAGCCGATTAAATCCTCCCCCTTCCATCCTTCCATTTTTCCTCCTCCTCCCGTGAGCTAGCCTCGACACTTGGCCAACTTTTGAAGTCGGCGGTGCTTCGGCCAGCTTGGCCATGTTTCACAGTTAACAGCCGCGCCACAGGCGCTTTTGGAGAACTGCGAACCCTTTGTCGCGGATCGATAACGAAATTTGGCCTCCAACACTTAGCCAAACTTACTTTGGGGAGGTAAAACTGCAGTGGGGAGTGTCTTATTGTCATGATAGTGGGGAAGATTTGGAGAAAATAGAAACTTGTGATTTTATTCCCAAAAACCGTGTCTTCATAGCCCTGTCAGCGTCGGCATTTGGACGTATTTCCTAAACCGCACATGATTTTGAGTTAATCGGTTAATGAGAACTTATATATCATCATGTCTAGTGGGAAAAATATGAGGTGAGAAAACAAAGACTTGCGTTTTTGGCCTGCGCGGATGTTAACGCAATGTGATTTTACGCGATTATACGAATGTCAAAGTGAAGGAATTCAGCAAAAGCGGCGGGTAAGCGGCGGGAGTAACTATGACTCTCTCTTAAAGGTAGCCAAATGCCTCGTCATCTAAGTGTGTGATGAATGGATTAACGAGATTCCTTTGTCACATCTGCTATCCAGTGACCTGACCCGAGGCGGGCTTGGCGAGGTCATTGAGAAGAAGACCTGTTGAGCTTGACTCTGGTCAGGCTTACAATGAGGGAAATGACTTGAGAGGTGTAGTATAGAGTAGGGCCTTCGGGTAGTGAAATACCACTACTTTTAACGATTATTTTACTTATTAGTGAATCGGAGCGGGGCACTACCCACTCTTTTTGGACCCCAGCCTTGGTATGTTGATCGTAGACATTGTCGAGTGGGGAGTTTGGTAAAAGTAACTTTACATCATTAAAAGATATGCAGTGTCCTAAGATGAGCTCAACGGAGAACATAAATCTCGTATGGAACAAAGGGTAAAAGCTCGTTTGATTCTGATTTCCAGTAGAATACGAACCGTGAAAAGCGTGGCCTATCGATCCTTTAGACCTTTGGAATTTAAAGCTAGAGGTGTCGAGAAAAGTTTCTGAGGATAGGCGGCTTGTAAGCGGCCAGGCGTTAAATAGGCGTTGCTTTTTGATCCTTGATGTCAAATCTTCCTATCATGTGGACGAGAATTCACCAAGTGTTGGATTGTTCACCCACGATAGGAGCGTGAGGCTGGGTTTTAGGTAGTCGTGGGAGCCAGGTTAGTTTACCCTCTTTGCGATGGCAGGTGTCGCAATAGTAATTCAACCTAGTACGAGAGGACCGTTGGTCGCCTGGTGGTCATCGCTTGGTTGAAAAGCCGAGTGGCGTGGCTTCGTCTGTTGGATTGTAGTGACAGCCTCTAAGTCGGAATCCCAGGAGCTAGCGCCTTATCGCCGCCGTCAGATTTACCCGACCAACGATGGGGCCTAAGGCCCCAAAGGTACGTGTTGTTGAGTGAAGCCCTCGCCGGGGACAGGCTGCGAGGCCGCCTTGAAGTACAATTTCTAGGGCGGCGGGTAGAATCCTTTGCAGACGACTTAAATACGCGACGGGGTATTGCCAAGTGGCGGAGTGGCCTTCTTGCCACGATCCCACTTTGGTCAGCCTGCGTCGCGGTCGTCCCTCCCCCTTCCCATCCTTCATTTTCCTCCTCCTCACACGTGGAGGCTAGCCTCCAACATGGCCAACTTTTGAAGTGGCGGTGCTTCCGGCGGCAGCGTGTTCAGGAAGTTAACCAGCGCGGCCTTTTGGAAACTCTTGAACCGCTGTCATGGGGATGGGGCCATGAGTGGCCTCAATGTGGCAAACTTACTTTACGAGAGTCAAACTTTCGTAGGAGTGTCTTATTGTCATGATAAGCGAGTGGGAAAATTTGAAGAAAATAGAAACTTGTGATTTTGGCTTAAAAACATTGTGCCTATAGCCCTGTGGCCAATCGGCATTTTAGACGCAACTCATAAACCGCACATATTTTGAGTTCCGTCCCAGTTAATAGAGACTTATATATATCATCATGTACAGTGGGAAAAATGTAGAGGTGAGAAACAAAGACTTGCGTTTTTGGCCCAAAAGCGAAATGTAACCCACGGGAATCGAGCGAAATCGGCATTTGGACGTAACTCCATAAACCGCACATGATTTTTTTGAGTTAGTCGATTAAAGAGGGCTTATATCATCATGTCTGATGGGAAAAATGAGGTGAGAAACAAAGACTTGCGTTTTTGGCCCAGCCGTGTAACCTGAATCAGGCGAAATCGGCATTTCGGACGTAACTCATAACCCGCACATGATTTTGAGTTAAAATCTCGAGTTAAAGAGGGCTTATATCATCGTATGCGAGTGGGAAAAATGAGGTGAAAATAAAGACTTACTTTTTGGCCCAAAAAAGCGAAGTGTCTTTATATAACCTGAGCTTCGAGAGAAATGGGCATTTCCAATTTTCAATTTTTTTTTTTTTATAGTTTCTCAACTTTGTAGTGATTATGAACATTTGTTTAGTGTGAATCTAAAGTTTCTAATATTTTGAGGAAAATTTTATGTTTTGATTTTTGTGGAAAATAGGTGAACCCCGTGCCAAATGGGCATGCTGACATTTGTATATATATATATATATATATATATATATATATATAAGGAACCGGTATTAGCAGGGGTAGCCTTACGAGGAGGCTAAAGTATGGATGCCGAGGGAGTTTGAGGGGTTGCCTAGGTGCGAGGCATGGATCACAGGGGACCCAGTGAGCTTGAGGTAGCCTTCGAGGAGGGCTAAAAGTATGGATGCCGAGGAGTTTGAGGGTTGCCTAGGTGCAAGGCATGGAGGCTGCCGAGGAGGTGCAAAGCATGGGTGCGAGGGATGCGAGATGCGCGCGTACTGTAGGGGAGCTGCCGCGGTCTTGTGGCGTGGTGCCAAGGTCTTTCGCGATGCTAGCGAGCGAGAATTTGCGAAAAGTACAGTGGATCTGCAGTCTTGTATTTATGGGCTCTGAGGGGCTATGGCCAGTCTGTGATCAAGGCTGAGAAATTCTTAAGTGGATGCTGTGATGGGCGACAAGGCGGCGGAGAATTTCTGCCGCGTCTGTGACATGGGTTCTTAAGTCTTTGCCAGCGATCTGCTTGTGAAGTACAAATTACCGAGGTCTTATGACATAAATTTCGAGAGGCTGCCAGCAGTGCGAGCCAGGTTGTGGTTCAGCGGTGTCTGAATGTTGCCAGTCTTGCTGATGCCAGCCAATTGCGAGGATCTGTGGTCGCTGACGTGCTCACGAAGAGAGCTGTGGTCGCGAGCCTTGAGCTGCGTCGGGTTACCCAGCGGTGTGCGAGCGCCATTTCTTTAAGTCTCTGCTGCGATCTGTGACAAAGCTGCGAGGATCTGTGATCCGCCCGACCGAAGTTTCGCGAGGGGACACTACAAGCGAGCCACTGTAAGTAGCGCGTCGAGCAACGGTAGGGGTAAAACCCTCAGCTATAGTGGGCTATAGTGTTTCCGTTTGTAGTGTCCCTCACAGCCTGCTGCACCTGCAACTCCTCGCAGCCTTGCTGCACGCGCGGCAACACCTCGGCTGGCCATGCTACCACGGTTTAACCAGCTGCAACCTTAACTCGCACCGCGGCAGCCCCTCGCAGCCATGCGCACCACCTCCCCTCGCAGCCTTGCTGGTGACGGCACCTCGGCGGCATTCTGCACGCCGCGGCTGGCCATCTGTGCTGGCTCCGCCTTGCGGCAGCCCACCGGCGACCATCTTCTTGAAACACTAAGCGATATCTGCGTCTTAGACCTCTCGCCTTCTTCTGCGGCAGCCCCTCGCCACTGACTTCACCGCCCTGCCCGACCCCTGCGCAACCATCTTCTAGCATGCCCAGCGGCATCCTTTCGGCGACCCCTCAGCGACCTTCTTCTTGCTCAGCAGCGACAACGTCGCGGCTAGCGCACCTTGGCGGCTTTTCCGGTAACATGTACACGCGGCGTATATCATGCTTTGCGCCTGGCTGACCATGCCTTGCACCCTAATGACCCCTCAAACCCCTCGGCATCCATGCTGGCCTCCTCGAAGGCCACCTCGGCTCACTTTGTCCCCCTCGGCATCCGTGCCTTGCACCTAGGCAACCCCTCAAACCCCTCGGCATCCATGCCAGCCTCCCTCGGAAGCTACCTCAAGCCTGCCCCATCCCCATATATATATATATATATATATATATATATATATACAAATGTTTGGCATGCCCATTTGGCAGGGTTCACCTATTTTCCTGAAAAAGCTAAAAAAACATAAATTTTCCTCAAAATATTAGAAACTTTAGATTCACACTAAGCAATGTTCATAATCATCATTAAAGTTGAGAAACTATAAAAAAAAAAAAATTGAAAATTGGAAATGTCGATTTCTCTCGAGTTTATGGGTTATATAGCACGGTTTTGGGTAAGCATGATCTTATTTTCACCTCATATTTTTTTCCCACTCGTACATGATGATATCTCAGCCCTCTTTTAAGCGGAACTCAAAATCATGTGCGGTTTATGATTCGTCCGAAATACCGGTTGCTCGATTTTGTAGGTTACAGAAGTTTTGGGCCAAAGCGCCGTCTTTATTTTTCACCTCATATTTTTCCCTTTGTTGTGATGATATATAAGCCCTCTTTTAAGCGGCGGAACTCAAAATCATATGCGGTTTATGAGTTACGTCCGAAATACGATTGCAACTCGTTCATGGGTTACGGTTTGGGCCGCGCCGTCTTTGTTTTCACCTCATATTTTTACTCGTACATGATGATATAAGCCCTCTATTAACTGTGGAACTCAAAATCATGTGCAGTTTATGAGTTACGTCAAAATGCCGATTTCGCTTGATTATATGGGCTATAGCACCACGGTTTTAGGCAAATCTCACAAGTTTCTATTTTCTCTTCAATTTTCCCCACTCGCTTATCATGGCAAATGAGCGCTCCAGTTTTGCCTCCTCGAAAGTAAGTTTGGCCAAGTGTTGGGCGGCCCTACGCTATCGATCCCTGTGTTAAAGGGTTGAGTTTCAAGCGCTCGCTGGCAGCGTGGTTAACTCGGAAAACATAGCTAGCGTCAGGCTGCTGGGCAAAAGTTGGCCAAGTGTTGGGTAGCCTGTGGGGAGGAGGAAAAAATGGAAGGATGGAAGGGGGGAGGGACGAATCGACTGACGAGAGTGAATCTCGGTGGATAAGTAGCGTGAGCCACTCTTGCCACTACAATACCCATCGCGTATTTAAGTCGTAAGAGGGTTACACCCGCCTTTCAGTGGTATGGTGGCGGCCTACGCGGCTTGTGTCCCGGGCTTGACCAACAACACGTGCCTTTGGGGGCCTAGGGCCCCTCTGCCGGAGTCGGCAAGCAGACCGGCGAAGTATCAATGTTAAGCTAGCCCGGATTGACTTAGAGGCGTCAGTCATAATCAGTATACGGTAGCGCGCCAGCTCAAGCGCGATGACCCGATTGTGCGAATCAACGGTTCCTCTCGTGCTAGAGTTGAATTACTGTTGCGACCTTTGTCATCGGTAGAGGTAAAACTAACTTTGTCTCATGACCCAGTCTAAACCCAGCTCACGTTCCTATTGGTGGGTGAACAATCCAACACTTAGTGAATTACCTTCTGATGATAGGAGAGCGACAGTAGAAGTCAAAAGCAGCGTCGCTATGAAGGCGCATTTGCCTGACCGGTTATCCTGTGTGTACCACGACACCTCTGGCACAAATTCCAAAGATTCAAAGGATCGATGAAATACGCTTTCTCAGTTCGTATTAAATCTTGGAAATCAGAATCAAACGAGCTTTGCCCTTTTTGTTCCCTGCGGTTATGTTCTCGTTAGGCATCTTAGGACCTAATTTATCTTTTAACGGATGTGCCTTGACCAAACTCGACAATGTCTTCCGCTTAGGATCGGCCATGGAAGCGAACTGGAGTCAAAAGAGAGTGATGCCGCCTCCGATTCACGGAATAAGTAAAATAACGTTAAAAGTAGTATTTCACTTTATAGAAAGCTCCCACTTATACTACACCTCTCAAGTCATTTCACAAAATCGGACTAGAGTCAAGCTCAGCAGTCTTCTTTCCCGATTAAGCCGTTCCCTTGGTACATGGATTTATTGGATAGTGACAGGGGCCGATTAGGAATCTCGTTAATCCGATGCATGCGCGTCACTAATTAGATGGCGCGAGGCATTTGGCTACAAGAGAGTCATAGTTACTCCCGCCGTTTACCCGGCTTGGTTGAATTTCTTCACTTTGACATTGACCTTTGGGCGAGAAATCACATTCGTTAACATCCGCGGGACCAAAACGCAAGTCTTTGTTTTACCACCTCATGTTTTCCCACTCGTACATGATGATATATAAGCCCTCTATTAAGCGGCGGAACTCAAAATCATAATGCGGTTTATGAGTTACGTCCGAAATGCCGATTTCGCTTGATTATGTGAGCTATAGCACGGTTTTTGAGCCAAAATCACAGTTCCTATTTTCTCTTCAAATTTTCCCCACTCGCTTATCATGACAATAAGACACTCCACGAGTTTTACACTCCTCGAAAGTAGTTTGGCCAAGTGTTGGAGGCGGCCTCGGTGCTCCGATCCCCTGCGACAAAGGGTTCGAGGTTTCAAAGCGCCTGCGTCTTGGCGTTGGTTGCTGCGGAAAAACACTTAATGAGCGTCGGGCTCTTGCCGACTTCAAAAGTTAGCCAAATGTTGGAGGCCCCAACCTCGCCAAGGAAGGAAAAATGGAAGGATGGAAGGGGAGGGACGAATCGGGCGACCTTGGTGAATCTCGGTGGACGTGGCGTGAGCCCACTCACGCCGCCTGCCCAATCGCGTATTTAAAATTAGCGCAAAGGATTCTACCCGCCGCTCGGTGAGAAATTGTAATTCAGGCGGCCCTCGCTGACTTGTCCCGGCGAGCTTGACCAGCGACACGTGCCTTTGGGGGCCTAGGGCTCTGCCGGAAGTCGCGAGCGGGCCAGCGGGCGCATGCGTAAGCGCGGTTCGGCTTAGAGGGCGTTGATGTAATCCGGTATGACCTTATGCGGCTTTTCAACCAGCGCGATGACTGGTGTGCGCGATCAACGAAGGTTCCTCTCGTACTAGGTTGAATTACTATTGCGACACTTTGTCATCGATGAGGATAAAACTAACTTTGTCTCAGTCTAAACCAGCTCACGTTCCCTATTGGTGGGTGAACAATCCTTCACTTAGTGAATACCTTCACAATAGTAGGAAGTAAACATCGAAGGATCCAACAACGTCGCTATAGGCGGCTTGCATTTGCCACAAGCCGGATTATCCACACAATGTAACTTTTACGACACCTCTGGCAAATTCCAAAGTCTAAAAGATCGATGGGCCACGCCTGCAGTTCGTATTAAATCTTGAAATCGAGAATCAACGAGCTTTCACCCTTTTGTTCCACCCGAGATTTATTCTTGTTGACTCATCTTAGGACACCTGCGTTATCTTTTAACGGATGTGCCTTCGACCAAACTCCCCACGGGCCGTTACGAGAACGCACTTGAGATGCGAGCTGCAACGCGAGGCAGCGCGTCTGCCCACCATGATCGCGTCAGCGGCGTCGCGGGCATATCAAGCGGCCGGGCATTGGCCGCGCCGCGATCGCAGCGGTCCGCCACAGGTCGAGTGGCGGACCGGCTTGTGGCGTTCCACATCCGACGGGCGCATCGCCGGCCCCATCCGCTTCCCTCGGCCAATTTCAAGCACTCTTTGACTCTCTTTTCAAAAGTCCTTTTCATCTTTCCTCGCGGTACTTGTTTGCTATCGGTCTCAGCCCGTGTGCCTTGGGCCGGAATTTACCGCCCCCGGTGGGACTGCATTCCCAGCAACCCGGCTCGCCGGCAGCGCCTCGTGGTGCGATAGGGTCGAACGCAGCCAGGCTCTCTCACCCTCTGTGGCGCCCCCTTCCGAGGACTTGTTCGATCGCGGCCGAGACGCTTCTCAGACTACAGGTCGGGCCTTGTGCGCCTTGATTCTCAAGGTAAAAATTTGTTCCGGTTCGCTCGCATTTACTAAGGAATCACTTGTAGGATTTCTTTTCCCTCCAGCATTGATATGCTTAAATTCAGCGGGTAACCGCACTTTGACACAGGGGTCGCGTTGAAGCGCCGGCGGCGTAGCGCATTAGATGGGAAAAGCGGGGCGGCGGCTGACGGGACGCAGGTTTGACAACCACGATTGTCAGCGGCTGATCGCCCGAGCTTATTATTTATGCCAACGTGACGGGCGCGGGAGACCATCGTCCACCCGACTCGAGGAGGGTTGGGAGGGCAACGGCGTGTGGCCTTGACGGAGCGTGCCCTCGGCCCCTAATGGCGGGCGCAACTTGCGTTCAAAAACTTCGATGGTTGCGAAGTTACGCAATTCCACACCGTATCGCATTTACGCTCACGTTCTTCGCTCGTCACGAGCGAGATGCTCCGTTCTGCGTCGTTTAGACAATGAGGCAATGAACCACCCTTGTGATCACGATCGGGGACGGCGGGGGTGGCACTCATTCCAAGTTCCTTGGCGCAATTCACCGCCCGATGTTGATGCGCCCGGAGAGGTCGGCCAAGCGGGAGCGCCCGAAGCCACGGCCGCGAGGCCGCTTCACAGGATGAGGAGGCGGAGCGCGAGGCCGTCTCAAAGCGTTGGGAAACGTTCTGGGGTGATGTGGGCGGAGTTTAAACAATGATCCTTCATGAGTTCACCTACGGAAACCTTGTCTGACTTCTCTTCACTCTAAATGATAAGGTTGATGGACTTCTCGCGCGTGCAAGCAGCCCGAACCCGGCCGATAACGCGATCGAAAACTTCACCCGGACCATTCAATGGTGGGCGGCCAGGCGGTGTGTACAAAGAGCGGGGCGTAGTCAGCGGTGACTCGCGCTTACTAGAATTCCTCGTTGAAGACCAACGGTGCAATGATCTATCCCCATCACGATGAAATTTCCAAAGATTACACGGGCACCTTCGGCCAAGGCTATAGACTCGTTGAATCACGGCCAGTGTAGAGCGCGTGCGGCGAACATCTAAGGGCATCCACCCAGGACACATTGGCTGCCTCAAGCTCCCTTAGCCTAGCTGACCATGGTCCCTCTAAGAAGGCGCGCGGAGGAGACCTCCCACAGCTAGTTAGCGGCGAGGTCTGGGTTCGTTAGCAGAGTGCCCGGTTCAAATGCTCCACCAACTAAGAGCGGCCATGCACCACCACCCATAGAATCATGAAAAAAAGAGCTCTGGTCTGTCAATCCTTACTATATCTGGACATTAAGTTTCCCGTGTTGAGTCAAATTAAGCCCTTGGAGCTCCACTCCCTGATGGTGCCCGTCAATTCCTTTAAGTTTTAAAGCCTTGTATTACATTACTCCCCGGAACCCAAAAACTTTGGTTCTCATAAGGTGCGGCGGAGTCCTAAAAGTAACATCCGCCGATCACACAGTCGGCATCGTTTCTCCGTTGAGACTAGGGTGGTATCCCCGGTCGTCTTAGACCCCCAACTTTCGTTCTTGATTAATAGAAACGTCCTTAGCGAATGCTTTCGCGATTGTTAGGTATACATAAATCAAAATTTCACCTCGACTATGAAATCTGAATGCCCCATTTGTCCTGTTAATCATTACTCGCGATCCGAAGGCTGAATAGGACCGAAATCTGAAGTGTTATCCCATGCTAATGTATCCCGGAGCGTAGGCTTGCTGGGCACTCTAATTTCTTAGTAAGCCTGCAGGAGCACCGGCCCGGCAATTAAGGCCGGGAGGCGTATCGCCCGGTAGAAGGGCAGGCTTGACGGTGCACACCATAGGCGGACGTCGGCCCAACCCAAGGTCCAACTTCTGAGCTTTTTAAGCGCAGCAACTTAAATATACAGCTATTGGGCTGGAATTACAGCAGCTGCTGGCACGGACTGCCTCGATGGATCCGCTCGTTAAAGGGATTTAGATTGTGCTCATTCCAATTACGGACTCATGAGGCCGGTATTTATTTGTATCACTACCTCCGTGTCGAGTTGGGTGAATTCGTGCTATTATTGGATCGTGATAACTGATTTCTGGCTCCTCTCCGGAATCGAACCCTAATTCTCATGCTTCCCGTCACCACCATGGTAGGCCACTATCCTGCCGCCGAAGTTGATAGGAAGGATATTTGAATGATGCGTCGCCGGCCTAAGGGCATGCGATCTAACAAACCGAGGATATATGAATCATCAGCTGTGAAAGGCCAGGCGTTATTTTATCTAATAAAATGCACATCCCTTCAGGAGGTAAAGTTTGTACACGTATTAGCTCTAGAGGAAATACTAAGATTTATCCGAGTAGTAGAGTACCATCAAACAAACTATAAACTGATTTAATGAGCCATTCGCATTCACGGTCTCAGATTAGTTCATGCCCACACATGCATGGCTTAATCTTTGAGACAAGCATATGACTGCGCGGGATCAACCGGGTAGCATTCCTTTAGACCGCCGCGCACCGCGCTGGAGGCCCCGCGGCCCAGCGAGCTGACCGGGTCGTACATGTCTAAGCGACCAAAGCTTGAATTTGGACATCGGGTGTCGAGACACCCAAGCCCACAGATGTTCCGTATCGAACCATCAAGTAGCAGCCGAGAGCCCGTAGCGTGCGCGCCTTACGGCGTGCTCGATGCGAGAACCTGGCCCTTCTCGATGTCCTCCCAGCGGCGCAAGCTCGGGAAGGAAAGGGTAGCGGCGGCACGTTCCTTTAAGGCGTAGGTAGCAATGCGGGGGTCTCAGTATAGCACAAGAAAGGAGTCTTAGCTGCGTCAGCCGGCGTGGTGTGGGCGGGCGGTTCGATGCCCGAGCACGGAGCCACTTGACCCACAGCCAAGACCTGACTCACCTTATTATTGCATCGATGGGCCAGCTGACATTAATTGACCACCTTTGGGCTTACTTGGCTTAAAACCATAAGCGATTTTATATGGCCAAGCCGCACCGCGGCATGAAGAAAACATAAACACCCACCCAATAGGCCTTGCCCAATGGCAAACTCATTCAGCGCACCGGAGGAAGTGTGGGTAAGATCCACGGAGGGCAAACCCCTCGGGAACCGAGCGAGGACTCGTTGTATATTTGGACACCACGCTCGACTCACTCCTGAGAATGAACATTAGAGCACAATGCGCCTTGATATCCGGTTAAACAAACTGAGAACCATATCTATAGATCTTGCCCATGCTTGGTAGAACAAACTCACATTACCGGCCAGAAACCATTCCCGTCCCGTTCGAGGGCTACCCCCTACTATATGGAGATTGTGGCATGCCCATTTGTACGAGTTCACCCATGTTTACAAAAGTGAAAATCGCGAATTTGGAATCCCTCGGCGGCCCCTCGCGGCGAGAACGCGCATACAGCGACGGCACCTGTGGTTCAGCGGCTGACACCTGGCGGTACTGCTGCGCCTGGCGGCGCATCCCCTCGCGGCCACTGCTGCGCGCCGTGGCATCCCCTCGCGGCCCTTCTTTGCGGCGTATGGCAAACACCTCGTGACCATCTTTGCACCTGCGCGCTGACCAGCCCGCGACCTTAGCTGCCGCCGCGGCGACCCTCGCGGTATCGCTGCGCACCGCGGCACATCCCCTCGCGGCCATGCCGGGCCCCAGCCTCGGCGACCATTACGCACCGCGGCAACCGTGCCCAGCGCCCGGGCTATGGCGGCCCCTCGCGGCCATGCTGCGTACCTCGCTGACCCCTGTGCTGCTGTTATCGCGGCTTCCTCAAAGCGACCATCTTTGCGTCATGGCAGCATCCTCTCGCGTGCGCCACATGACATCGGCAGCCCCTCGCGTATCAGCTCGCACCGCGCGGCCCCTGCCCGGCCGTACTGCGCGCATCCACTCGTGACCCCCTCGCCGGCCTTGCTGCGCCATCGCGGTGACACCTTAGCGTATATTGTGCGCCTGACCCCTCGGCCCGACATGTTGGCTCTCGGCATCCCCTTGGCGGCTCTTTGCACCTCGGCGGCCGTGCCTTGCGCTAGGCAACCCTCAAACCCCTCGGCATCAATGCCCCAGCCTCCCTCGAGAGAGGCTCTGTCAAACCTCATTTGTCCCCTCGGCGTCCATTGCCTTGCACCTAGAGCAACCCTCAAACCCCTCGGCATCATGTATAGCCTCCCTCGGAAGGCCTCTTCTTGTCCCCCTTATATATATATATATATATATATATATAAAATACGCCAGCATGCCCATTTGTGCTTGAGGTTCACCTTTTCTAAAAGCTGAAAAACATAAATTTTCCTCAAAATATTAGAAACTTTAGATTCACACTAAACAATGTTCATAATCATCATTAAAGTTGAAACTATAAAAAAAAAAAATGAAAATTGGAAATGTGATTTCTCTCGAGTTTGTAGGTTATAGCACCTGGTTTTGGGCCAAAAACGCAAGTCTTTATTTTCACCTCATATTTTTTCCCACTCGTACGCGATGATATATAAGCCCTCTTTTAAGCTGACGAACTCCAAAATCAATATGCGGTTTTATGAGATTCGTAAATCGATTTAAGCTCGATTTTGTGGGTTATAGCACACGGTTTTTGGGCCAAAAACGCAAGTCTTTATTTCACCTCATATTTTCCCTCGTACATGATATTCTCAGCCCTCTTAGCGGCGGAACTCAAAATCGATATGCGGTTTACAGATTCGATAATCTGCGATTTAAGCTCGATTTTCGTGGGTTATATACCGCGGTTTGGGCCAAAAAGCGCAAGTCTTTATTTCACCTCATATTTTCCCACTCGTACATGATATTCTATAAGCCCTCTAATGGCGACGGAACTCAAAATCATATGCGGTTTATAAGTTACGTCGAAATGCCCGATTTAAACCCAGTTATGGGCTATAACCACCCGGTTTTGAGCCAAATCACAAGTTTTATTTTTTCTCTTCAAATTTTCCCCCTTTCGCTTATCAGCAATGAACCTCCCGAAAGTTTTACCTCTAAGTAAGTTGGCAAGTGTTGGAGGCCCGACCTGCTTCTCGATTACTGCGTGAGGTTGAGTTTCAAAGCGCTCGCGCGCGTTAGTTGGCTTTCGGAAAACATGTGGCGTCGGAAAGCAATTTGCGACTTCAAAAGTTAGCCGTGTTGAAACTGTCTCAGCGAGGAGGAGGAAAAATGGAAGGATGGAAGGGGAGGGGCGAATCGAAGCTCAGCGCAGGGCCGAATCTCGGTGGATGCGTGGCGTGAGCCACTGCCCGCCACAACCCGTGTGCCCAAGTCGTCTGAAGGATTCTACCCGCCTTCGGTAGAAATTGTAATTCAAGGCGGCCCCACCGCAGCTTCTCCCCGAGGGCTTGACCAACGACACGTGCCTTTGGGGCCTAGGGCCCTCTTTGCCCAGTGTGACGGACCGTGGCCCGCGTACGTCCTTCTAGCCCGGATTCGACTTAGGGCGTTCGATGTTAATCCAGTATGGTAGCTTAGCGCCACTTGGCTTTCAACCAAGCGGCGATGACCAATTGTGCGAATCAAGCAGTTTCCTCTCGTACTAGGTTGAATTACTATTGTGACTTTGTCCTTGATGAGGTAAAACTAACTGTCTCAAGCGGTCTAAACCAGACTCACGTTCCCTATTGGTGGGTGAACAATCGCTTACTTAGTGAATTAACTACAATGATAGAAGAGCCCGACATCGAAGGATCAAAAACAACGTCGCTATGAATGGCAGTTGCCACAAGCGATTATCCTGTGGTAACTTTAAGCCTCTAGCTTTAAATTCCAAAGGTCTAAAGGATCGATAAATGCGCTTTCCGCGGTTAAATAATGCGTCTTACAAATCGAGAATCAAGCGAGCTTTTTACCCTTTTGTTCCACGGTTTACTGTTCTGCGTTGAGCACAAATCTTAGGACACTGGATTATCTTTTAACGGATGTACCGCCCAGTAAACTCCCCACAATGTCTTCGCCGGGATCAGCCGCGAAGCGGAGCTTTAGGTCAAAGAGGCGGTGCCGCCTCAGGTTCACGGAATAAGTAAAATAACGTTAAAAGTAGTGGTATTTCACTTTGCCCAAGCTCCCTTATACTACACCTCTCAGTCACATTTCACAAAAGTCGGACTAGAGAGTCAGCTCATGAGGTCTTCTTTCCCAGTTACCTTCGCTGTTTCCCTTAGGCGGTGGTTTATTTGGATAGTGAGCGGGGGCGGAGTGGAATCTCGTTAATCCATTAAATGCGCGTCACTGAGTGGATGGCCCGAGGCATTTGGCTACCTTAAGAGAGTCGCAGTTTACTCGCGTTTACCCGGCTTGGTTGAATTTCTTCACTTTGACATTGAACCCAAGTGAAATCACGACGTTAACGTCCGCGAGACCATACGCGTGCTTTGTTTTAATTAAGCAGTGGATTCCTTATCCATTGCGATTAAATCGGCTGACGAGCGGCAGAAAACCCGGCGTTCGATCAAGATCCCCAGCAGCCATGGCGACCCGCTCTACGCCATGAGCGGCTTCAACCGATTCGCCCGGCGGCCGGCGGGTTCAAGAGGCCCCGGACCCGTACCAGACCCTCGAGGCCAATCCTTTCCCGAGTTACGGATCCATTTTGCCGACTTCCTTGCCTACATTGTTCCATCGACCATTTATTCACCTTGGAACTGATGAAGTTATGAAATGCGGCCGTGGAGTGGCACTTTCGGTCCTCCAGTTTTCAAATGGCGAGCTGCCGGACACCACGCGGCGTTCGGCGATCTCTCGGCCGTGGACCTACCTCCGGTGGCCGTTTCAGTGGGCGGTGGTAAACGAAAGATAACTCTTCCGGGGCCCTTCGACGTCTCGGACTCCCTAACGTTGCCGTCAACCACCGCGTCGGTTGGGGAATTTTAACCCGATTCGCCTGGGGATTGGCGAGTCGCGCTATCGACGGGGTTACCCGGTCTCTTAGGATCGACTAACCCATGTGCAAGTGCCGTTCACATGGAACCTTTCCCCCTCTTGACCTTCAAAGTTCTCATTTGAATACTTGCTACTACCACCAAGATGCACCCGTGGCCTCCGCCGGGCTCCGCCCAGTTTTGCGGCCGACCGCCCAGCGCCCTCCCTACTCATCGGGAACTGGCACTTGCCGACGGCCGAGAGTGTAGGTCACGCTTAAGCGCCCATCCATTTAGGGCTAGTTGATTGGCGGTGAGTTGTTACACACTCGCCGTGATTTAGACTTCCCGCGGCCACCCGTCCTGTGTGCTTAATCGACCAGCACCTTTTGGCCAGGTTCTAGGTTGTGTGTGGTTTAGGCACCGTAACGGCTTCGGTTCATCTGCATCGCGATTCTGCTTACCAAAAATGGCCCACTTGGGCTACCGATTCCCATGGCGGCGGCTCAGCAGCCGGCCCGGCGCTGTCCACCTATTTAAAGTTTGAGAATAGGTGGGCATTCTGCCCCCGATGCCTCCTAATCATTGGCTTTACCCGATAGGGCTTCCGTTCACGGGCTCCCGGCTATCACGAGGGAAACCGGAGGAACCCAGGCTACTGGACCGATTAGTCTTTCGCCCCTATACCCAAGTCGGATAGGTGATTTGCACGTCGATGCTGTGCAGGCCTCCACCGATTTCCCTCCTCCTCCCTCCTCCTCCTCCTTCCTCCCTTGATGGGGGCATTAGGCCTCCTCCGGCCAGCTCAAGAGAGAAGGCACCATCCTGGGAGGTTTCACACTGCGCCCTCGGCCTCCAGTGAGAGATCGCTATGGGGAGAAGTTGGCGGCCCGGCATGATGATTCGATTGAGAAGGAAGAGGCGAAGCCTACGGGAGGAGCAATGTTATTCGTGGGAATCGGGGATCTCTCATGTGTTTGAGCGTCAACTTGGAATGAGGATGGTTCAAAAGGTGGTGGTTAATAAAGTAATGTTTTGGTGGAAGTTACGCGGTCAATTTCCGCAATTTTAGGTGGACTAAGAATCACGGAATGCTATGTATACTCCTCGATTAATGATACAATGACTCGATCATGGAGAGAAGGATTCATGCCTTTCCTCGTTGGTTCAAGGTGTAGGGAGGGTGGAAAGCACGTTCGTGATGTTCATCTTTCAGAGCTTCAACTCGGCTGACCATGGCGCTGATTCTCGAATTGATGGAAGTTCGTGAGTGCCCTGGTTTGGAGTTACGGGGGATAGGAATCGTATCTTTGCATGCAACCAACTTTCAAATTGAAAGTCGATTGCTATCCCAATCCCGATTTCACAGGACTCCATCTCTAGAATTGAAGGGCGTTTCTATTAGATCGATCAAGGAGTTACAAACTTGAACTTTGATCGAACCACTCAAGGTTACAAACTTGAGTTCGAATATTTCAGGAACCCTTGATGTAGAGACGGAGGAGCTGCACACTTGAAATTTCCCCTCACAGGTAAAGCTACACCCAAGACCAGCACCGCGTCCAAGGCACAAACAGCTACTTGGCCTCATTGGAACATCCGGTTACTCAAGCGGTTGCTTCAACGAGATCAAGCAGAGGAGAGCACACGGGTTGCGGAAGCGGCTGCGTTGAACCTAGATCTTATGCCATTCTTTGCAGAGCCTACCTATTCCTTCTGTCGATTGTTGCGAGGTTCTTATATTAAGGAGCATTTGAAGGCATGCCATCGATTCACCTGTACTTTGATCATGAGAAGCGAAAGCTAATTTAGCTACACCTTGATTATCGCTTCGCGAATTGGTAGAAGCTGGAGACGGTCGAGGTATCATACATGGGAGTGTGGGGACTCTCTAAGCCGGTGGGAGTTACGGACTCTAGAAGAATTCTAATCGGCGATGAATCTTTGAAGACCTCATTGTAACAGCAAGGAACCATGGGATTATTGACAAGGGTACCTTACTCTTTGTTCACTTGGCAAAGTAGCACGGGAAGAAAGAGCGAATCTCCATTAATACAGCAGTGTCGTTTACCGAGGTTTCTGACCCAGGCTTTGCGTCGGACAATCCTTTCACTCATTGGTAATTCCTTTGCTCGATTTGTTGAAATTTGATCCGAGACTACAAACACTCTCGAGCCCGCACATCCCTCAGTTTGTGTTGAGTTGACGTCTCTAACCCACTTCCACAAGAGTTTTCCTTCTGGGTAGATGCAGATGGTTATTGGCGAGGCAATTGCTTTAAGAGGGCCCAACTTATTATTCTTTACGCAAAATGCATGGACACACAACTCCCCTTTGTCGCAAACGTATTTTGAGGAAGAGAAGCATCACATAGCAGTTAAGCAGTACCAGTGCGAATGGTTTGGAAGCCGAACTACACCTCCTTAGGTACCCAGGCAGAAATGCTAGATGCTTTGGAGCATCCGGTTCGGTGACTTGCTGACTTGACCCAAAATGCTACAACGGATACGATGACCATACCTTCTATACAGCGACCCCGAAATGTTGCAACCTCTAACGAGTTTTTACTTCTCTAGATAGAGGGTTGTGATGTTACTCTTGAGAGCATTAATGATGATACGAAAGGAAGTTATTGATGAGTTGGTTATTCACTTGTACTAAAAGGAGGAAAAACAAGTACAACCTGTAACAGATTGAGATGCCGGCTCGTACTGCGAGATCTTCCAAGAGGACCCAGCACCGAGGCGCTTGTAGACCCGAGCAACTCTTCTGCAACTCCAGGTTCAGAGCAAACAAAGACAAGCGAGCGGAACATCATTCATGGCGGGCTTGGACACTACGGCAGATGAAGAGAACCACGAGATCGAGTTGATATCGGAATGGAGCAGGCGCCCTTTGTGAGGATTAAAGAAAGAAGCGGGGATAGATTGACCACCTAGGAAATGCAACATTGCGCTCAAGGCTAGACTTGAAGCTCTAGTTTATTAAGTGTGTTATCTTAGTTTTTTTCTTTGCATTTTGGCATCCACGTGGAAGCACATTGTATCACTTAGGGTTATTAAATTAATAAAGGCATAGTTCACCATCTTTGGTCCCGGCGAATAAGTACTCACTCGAACCCTTCTCGGAAGATCCAAGTCGTCGGTGCCGAAAGGGATCCCGCACATTAGCTTCCCTACTTCATATACGGGTTTAACCTTAGTTGACTACTTTTAATATCGAGACTCCTTGATCAGTGTTTCAGGCGGGCCGAATGAAACAGTGCGTGCGAACGCATGAATCTGCGAGAAGCCGCAGCAAAGGCGGTAGCGTCGCGCCCACCGTAATCAGCGTCGTGGCGTCTCGCATGAGCATATCAACGGCCGGAGCTTTGGCATACGCGCAATCCGCAGCAGTCGCCCGGTCGAGTGGCGGACGGCCCGCGGCCGTTCCACATCCGACAGGCTATCGCGGCCCCCATCCGCTTCCCTCAGAGCAATTTCAGCACTCTTTGACTCTCTTTTCAAAGTCCTTTTCATCTCTCGCGGTACTTGTTTGCTATCGGTCTCTCGCCCGTGTTGTGGGCGGAAATTTCTTAGCCCGATTGGTGCATTCCCAAACAACCCGACTTCATGACAATCGCCTCGTAGTGCGTGAAAGTCCAAGGCTGTAGAGCTCTCACCTACTGTGGCGCCCCCTTCCAGGGGACTTGTTCGATCCGCCATTAGGCCTTCTCCGGTTACAGGTCGGAGCGCCTTGCGGCCTTGAGATTCTCAGGCCCCAGTGATCTACGGTTAGCTCGCCGTTACTAAGGGTCTTGCTTAAGTTTCTTTTCCTCGCTTATTGATATACTTAAATTAACGGGTAACCCGCTGACACAGGGTCGCGTTAAGCCTTCGGCGAAGCGTATGCATTAGATGGGGAAAAGCCGAGGATGGCCTTGTGACGGACACCGAGGTTTGACAACCCATGGAATTGTCTTGGCCCCGGGTAGCGGAGACTTATTATTTATGTAGCCATGTGACAGCGCCACGGGAGGCCATCTCAAGGCCCACCCAACTCCCGGAGAGTTTGGGAGGGGCAACGACGTGTAGCGCCGGTAGACGTGCGCAAACCTAATGGCTTGAGCAACCCCGTTCAAAAACTCGATGGTTCACGGGATTACGCAATTCACACCCAAGTATACAGCATTTATTTCACGTTCTTCGCTCGATGCGAGCCAAGATATCCGTTGCGAGTCGTTTAGACATATTGAAGGCGGCCGAACCACCGCGCGATCCGTCTCCGGGGGCCGAGGCGGTGGCAACTCCTTCATTCAGAAGTTCCTTGGCGCAATTCACGCCGATGTTATCACGCCAGGAGGAATTGACGGCAGGAGCGCGAAGCCACGACAGCGAGCTTAAGCTTCCGGGGATGAGAGCGAGGCTGCGAGGCCCCGTCTCCAGCGTTGAAACACGTTCGGGTGGATTACCTTTGGCGGAGTTTTACGTGATGATCCTTCCAGCGAGTTCACCTACGGAAACCTTATTCACGACTTCTCCTTCCTCTGATGATAAGGTTAGTGGACTTCTGGCAGCGTCGCGGGCGGAGCCGGCCACGTCGCGCGTCAAGCTTCACCGGACCATTAATCCCGGTAGGAAGCGCCCAGGCGGTGTGTACAAGGCGGGGACGTAGTCAGGCGATGATTCGCGCTTTAGGAATTCCTGGGTGAAGACCAACAATTGTAATGATCTATCCCCATCACGGTAAAATTTCAAAGATTACAGACTGTCGGCCGAGGCTATAATTCGTTGAATACATCCAGTGTAGCCGCGTCACCGACCCAGAGGCATCTAAGGGCATCCATGGACCTGTTGGCCTCAAACTTCCTTGGCCTGGCCATAGTCCTCTAAGAAGTGGCCGCGGAGAGACCTCCGCATAGCTGAATAGCAGTGAGGTCTCGTTAGATTACTTGGAGTGCGAGACAAATCGCTCTACCAACTAAGAACGGCCATGCACCACCACCCATAGAATCGGCAGAAAGAGCTCTCGATGCCTTGATCTTACTATGTCTAGACACAGTGATTTCCGTCATTGAGTCAAATTAGCCATGGGCCTCCACTCTGATGGTGCCCTTAGTGATTCCTTTAAGTTTATTGTGCGACCATACTCCCAAGACCCAAAACTTTGATTTCTCATAAGGTCTGGCGGAGTCCTAAAAGTAACATCCGCGATCCACTGATGGCATCGTTTATAGTTGAGACTAGGGCATTGTCACATTCGTCTTGAGCCCCTAACTACGTTCTTGATTAATGAAACTCATCCTTGGCAAATGCTTCGCGGTTGTTCGTCTTTCATAAATCCAGAATTTCACCTCGACTATGAAATACGAATGCCCCCGGTGTCTGTTAATCATTACTCCGATCCGAGAAATAGCGAATAGGACCGAAATCTTTATGATGTTATCCCATGCTAATGTATCAGGCGTGAAGCACTTTGAGCACTCTAATTTCTTCAAAGTAGCAGCCACTTAGGAGGCAGCCCCGTGGTGGGCCGGGCGTATCGCCGGTAGAAGGAGACGAGCGACCGGTGCACCATGGCGAATGAGTCGACTTGACGAGTCAACCTACAGGCTTTCATTGCTCTTAAATATACGCTATTGGAGTTGGAATTACGCTTGATGTGGCACGGACTTGCCCTCCAATGGATCCTCGTTAAGGGATTTAGATTGTACTCCATTCCAATTACAGGACTCATAGAGGCCGGTATTGTTATTTATTGTCACTACCTCCCAGTATCGGGATTGGGTAATTTGCCTTTCTTGCCTTCCTTGGATGTGGTAGCCGTTTCTCTAGGGCTCCCTCTCACGAATCGAACCTAATTCTCGGAAGTCACGTCACCACCATGGTAGGCCACTATCCTACCATCGAAAGTTGATAGGGCGGATATTTGGTGCGTCGCGAAGCCACTAAAGAGTTTTGTGCGATCCGTGAGAGGATATCATGAATCATATTCTAGCGGCGAGACCACGTTGACCTTTTATCTAATAAATCCATCCCTTCCGGAAGTCAGGTTTGTTGCACGTATTAGCTCTAGAATTTTCACGGTTATCCGAGTAGTAAGTACCATCAAACAAACTATAAGCGATTTAATAGGCCATTCGCGATTTCTGATGCAGAATTAGTTCACCTTACACATGCATAGCTTAATCTTGAGACAAGCATATGACTACACAGCGAGGATCAACAGATAGCATTCCTTTGCGGCCTTACATACGTAGCTGGGAGACCCAGTACTTGAGCAGCCCGATCATTACATGTCTAGCGACAAAAGCGCTTAGATTTGGACGCGAAGGTGTCCAGGACACTATTACAGATATTCATTATCCGAACCATCAAGTAGCAACCCGAGCCCGTCGCAAGCGTGCGCGCCCTTACGGCGTGCTCGATGCGCGAGGAGCCTGGCCCTTCACGCGTGTCCTCCCAGCGCAAGCTCGGGAAGGAAAGAGATGTGGCGGCCGTTCAGCTTTAAGCGTAGGTGGCAAATCTGGGAATCTCCGTATAGCACGAAAGTCTTAGCTGCGTCAGCGTGGTGTGTGGAAGTGAGACGGTTCGATCTGCGAAGCACGGAGCGCCAGCCCTGGCCGTTACAACTCCTGCGTTGCATCGGTGAGACCCGACAACATTAAGCCAAACCCATCCCAGTATGACCGACAAAGCATAAGCGATTTTATATAGCCAAGCACTTTCCGTGACATGGAAAACATAAAGCACCGCAATAGGCCTTGCAGGCAAACCGTCTTTGTTCAACTTTTGAATGATGTCGTTAAGATCAAAAGCGGAGAGACAAGACCCCCTCACGGGGTGAGGCGGGGACTCGTTGTTGGACTGCGCTCTGACTCACTGCAGGAATAGTGTTTAGAGCACAATGCGCCTTGATATCCAGTTAAACCAGGCTTTGAGAACCATATCTATGGAAATAACTTGCCCCTTACTTGGTAGAACAAACTCACATACCCGGCCGAAACCATTCCCACATCCGTTCCAGGCTATAACCCCCTACATATGGAATTGTGGCATGCCCATTTAGCACGGAGTTCACCCATGTTTCACAAAAAGTGAAAATCCCTTGAATTTGGAACCCCCTCGGCCCCTCGCGGTAGAGGCTGCGCATGGCAACAGTACCTCGCGGACCCTCGCGCGCGCCTCGCGGCCACTGCGCCACCGCGGCATCCACCATGACCTGCTGCGCCCAGCATCCCTCGCGGCCTTCTTTGCGCATTGCAGCGACACCTGCGACCATGCGCACACCGCGGCAACGCGGCGCGGCCTTGGCTTTCCATGCGCGGCCGGCCTCGCGGCCATCTTTGCCTGGCCATCCTCGCGGCCCTTCTTGGGCATCGCGGCTGACACCTCGAGCCATTACGCACACCGCGGCAACCCGTCGCAACAGCAGCTCATGCCAGCTGCCGCCTGCCAGCCATGTTGCGCACCTCGCGGCCCCTCGCGGCCTTGCGCGTCGCGCGGCACCTCGCGGCCATCTTGCGTGCGCGATGGCCTCTCGCGGCCTTGCTTTGCCCACCGCCCAACATCCCTTGCCGGCCTCGGCCTTGGCTCGCACCACCATGACCCCTCATAGCCATCTGCGCCTGTGACATCCACTCGGCGGCCCCCTGCGGCCTTGCTGCACGCCAGCTGACACCTTAGAAGCCATTTGCATACCCATGGCTGACCTCGGCGGCATGTTAGCGCATCTGGCATCCTTGGCGGCCATGCTTTGCACCTCGCGGCCATGCCTTGCACCTGAGCAACCCCTCAAGCCCTCGCGTCCATGCTAGCCTCCTGGAAGGCTACCTCAAGCTCTGTCCCCCTCGTCATGCCTTGCACCTAAGCTGACCTCAGAAACCCCTCGGCATCCATGCACTTTAGCCTCCCTACGAAGACTACCTCAAACTACTTGTCCCCCTTATATATATATATATACTCTAGTGTTGGCATGCCGTTGGCCTGGGGTTCACCTATTTCCTGGCCAGAAAACACCTCAAAAATGGCAAGGCTGTAGATTCACATAAACAATGTTCATATCATCATAAAGTTGAAAGCGATAAAAAAAATAAAAAAATTTAAAATTGGAAATGTAAGATTTCTCTCGAGTTTGTAGGTATAGCACACGGTTTTGAGCCAAAACGCAGTCTTTATTTTCACCTCATATTTTCCCACTCGTACATGATGATATATAAGCCCTCTTTTAAGCGCGGAACTCAAAAATCATGCGGTTTATGATTCGTCAGAGCGATTGCTCGATTTTGTGGGTTATAGCACCGCGGTTTTTGGGCCAAAAACGCCAAGTCTTTGTTTTCACCTCATATTTTCCACTCATACATGATGATATCTCAGCCCTCTATTAAGCGTGGAACTCAAAAATCATGTACGGTTTATGAGATTAAACCGAAATGCAGTTGGCGATTATGGGCTATAGCACACGGTTTTAGGCAAAATGCGGTTTATTTTCTCTTAAATTTTTTCCTTATCATGACAATGAGCCTCCCACGAAAGTTTACCTCCTCAGTAAGTTTGTGAAGTGTTGGAGGTGACCTCGCTACTCGATCCTGCGTAGGGTTGAGAGTTTCCAAAAAAGCGCTCATGCCTTGGCGTTAGTTAACTCGGGAAAACATGACAAAAGCGTCGGAGCTCTTGCCGACTTCAAAAGTTGGCCAAGTGTTGAAGCTAGCCTCGCCCGAGGAGGAGGAGGAAAAATGGAAGGATGGAAGGGGAGGGACCGAATCCCGAATGACAGTGAATCTCGATGGATAAGTAGCGTGAGCCACTCTGCCACTTATTACCCGTCCGTATTTAGAGTAAATGCGCAAAGTTCTACCCGCCCGCTCATTAGAAATTGTAATTCAAGGCGGCCCTCGCGGCTTGTGTCCCGCGAGCTTGACCAGCGACACGTGCCTTTGGGGGCCTAGGGCCCCTCGCCGGGTCGCGAGCGGACGATGGCGCGTCGTCACTTAGCCCGGATTCGACTTAGAGCGTTCGGTCATAATCCGGCCTGCAGTATTAATGGCGCCGCGGCTTTTCAACCAGTAGCGATGACCAATTATGCCGAATCAGCAGTTCCTCTCGTACTAGGTTGAATTACTATTGCGGCTCTTGTCATCGGTAGGGTAAAACTAACTGTCTCACGGCGGTCTAAACCAGCTCACGTTCCCTATTGGTAGGTGAACAATCCAACCCTTGGTGAATTCTACTTCACAATGATAGGAAAATGAGCATCGAAGGATCAAAAGCAACGTCGCTATGAGCGGCTTGGGCGCCTGGCCGAGTTATCCACCATGGTAACTTTAGAAGGCACCTCTAGCTTTGAGATTCAAGGTCTAAAGGATCGATAGGCCACGCTTTCACGGTTCGTATTCGTGCGGAAATCGAATCAAACGAGCTTTTACCTTTGTTTCACCAGTTACGTTCTCGTTGATCTCATCTTAGGACATCTGGGTATCTTTTTAGCGGATGTGCCGCCCGACCAAACTCCCACTGTGATGTCTTGGCCGGATCAGCCCGGCAAGGCGGAGCTTTGGAGTCCCAAGAGCGTGCCGCCTCGATTCACGGAATAAGTAAAATAACGTTAAAAGTAGTAGTATTTCACTTTCGCTTCGAAGGCTCCACTTATACTACACCTCTCAAGTCATTTCACAAAGTCGGACTAGAGTCGAAGCTCAACGGGTCTTCTTTTCCCCGTGATTACATGACCCGTTCCCTTGGTGTGGTTTACATTTGGATAGTAGACGGGGACGGTAGAATCTCGTTAATCTATTCATGCGCGTCATAATTAGATGACGGGCATTTGGCTACCTTAAGAGTCGTTAGTTACTCCGCCCGTTCACCCGGCGCTTGGTTGAATTTCTTCGCGACATTGAGCCTTTGGCAGAAATCACATTGCGTTAACATCCGCGGGGACCATCGCAATGCTTTGTTTTAATTAAGCGATTCGGATTCCCCTTGTCCGTACGATTCAGGTATTTCATTGAGCATACGGAAAGCGAAAGAAGCGTTCGATCGGAATCCCCGGCGGAAACCGCGCGGCGACCGCTCTCGCGAAGCGGCTCGAAGCGATTCGAAGCGGTAAGCGGGTTGAGACTAGGACCCCGTGCTGACCACCGAGCCAATCCTTTTGCCGAAATTCACGGATCCATTTCACCGAGCTTCCCTTGCCCTCGTTGTTCCCGCCGACCAGAAATTTTCACCTTAGGGGAACTGATAAGTTATAGTCGACGGGCGTAAGATAGCTCGGTCTCCGGATTTTCAAGGGCCGCCGGGGGTGCGGACACCACGCGGCCGTGCGGTGCTCTTCCGGCCCGGACCTCACATTGAGCCCGTTTCCAGTGGGCGAAATTTGTTAAAGCGAGAAAGATAACTCTTCCCGAGGCCCCGCCGGCGTCTCCGGACTCCCTAACGTTCGTCAACCACCGTCCCGGTTCAGGGTACAACCCGATTCCCTTCGAGTTCGCGCGAGTCGCGCTGTCGGGCCAGGGTTACCCGGAGTCTCTTAGGATCGACTAACCCATGTGCAAGTGCGTTCACCATGGAACCTTTCCCCCTCTTCGACCTTCAAAGTTCTCATTTGAATGTTGCTACTACCAGATCGCTGACCTCCGCCGGGCTCACGCCTAGGTTTGCGGCGGCACGCGCCCTCCTGCTAAATCGGGACGCACTTGCGGCGGCGGGTGAGTCACGCGCCAGCGCCATCCATTCCGGGGCTGGATTGATTCGGCGGTGAGTTGTTACCACACTCCTTAGCGGATTTCGACTTCATGACCACCTTCCTGGCTTGTCGACCAACACCCTTTGTGGGTTCTGGGTTAGCGCGTGATTTAGGCAAGTAACCGGCTTCGGTTCATCCCGCATCATGATTACGCACAAAATGGCCCACTTGGAGCTCCTGATTCCATGGCGCAGCTCCAACCGCGGCAGCGAGTCCTACCTATTTAAAGTTTGAGAATAGGTCGGGCGTTCTTGCGATGCCTCTAATGTTAGCTTTGCGATAGAACTCGTTCCACGGGCTCCATTATCCTGAGAAACCGGAGGAGACCCGGCTACTAGACGGTTCGATTAGTCTTTCGGCCCTATACCCAGTCGAGACGAGCGATTTGCACGTCGGTATCGCTGCGGGCCTCACGAGTTCTAAGGCAGCCCTCTCCCCTCCTCCCTCCTCCTCCCCTCGATGGAGGTTCGGCCTCCCACCCACGTCTAAAGAGAAATCCACGCAGCCCCACAAGGTCGGATCGCAAGGCGCTACCTTGGATTCATCCTTGGGGCATTTGACCATGGATCAGCCTCCACGTAGTGGACGCCTGGCTGCCGAGATCGGCATCCCAGGAGGCGGCACGAGAGTCTGAGTGAGGCGAGTTCGGATCTTGCGGAATCAAGTGCTTAGGAGGCGGATCTGTGCGGGCGAGCAGAATCGCGTCAGTTCTGGGGCGTATTCAACAGGACTCCTCCACGTCTGAGGCGAGATCACTCCATGGTAAGTTTGTGACGGAGTTTTCAAGCTAAAGTAGTCACAGTCTGCTGCACGGGCGAGATCTGAGGTTGTGCTGCCTTCCTCCAAGAAATCCGGATCTGAACCGGCCATGATCAAAGAGTCCTTCACATGCAGGACTCCCAAAATTTCGACCAAGGATAGGAAGTGCAAATCTAACTGTATGCAAAGGGGCCCAGCGTCTTTGTAGTCTTCTAGCCGGCGATGTTGCGGCCTTCCCCCTTTACTGCTGTAAAGCAGGTTCATGATTAAACAAAGATGCAAGGCACCCAGCTGACGGATAGAGCAACTTTGCCATATTTGTAATTTCTTCCGCAGCAGCGCAGAACTTCTGCCTTGGAGCAACTGCGGCGCAGCGGCTACAACGGGTCCTCGAGCTCGGCCGATCGCGGTGCCCCATCCATGGATGAATCGGAAGTGCTAGAGATAGCTCTTGTTACACAAGCGCGACCAACGTATACAGCAATCACAAAACCGAGACGATACCAGCACTTTCTGATTTTGAGATCTCACGGATGCATCATCGAGGACACCATTGAGAATTATCTGCGTTACATACTCGCCAAGTGATTTGATAAGGCAAAGGCCAATTTTGCATTCTCTTTGATCATGCGATTCACAGTGCGCGAAAACTTGAGGCAATCTGGTATCATCAATGGGACTTAGAGGGCATATCCACAGTTTCTTTGTTGGCGTCACAGACTCTAGAAGGTATTGATCAGTCTATTAAGCAAGGGCAGTATTATTTGCGCTCTCGAAGAGACTTCTGCATGACATCGATAAGTGAGATACTCACTCTTCAAGTGGCAAAACAACACAGGAAAGAAGGCGGAATCACCAATTGTTTACAAATGTTGACTTCGGAGTTTGCAACCGGAGCTGCGTAAGATTCCATCCTTACCTTTGATGGGAACTCTTTTGCGCGATTTGTGGGTGATCCCAGGCAAGCGAATACTATGAGAGCAGCGCATCACGGTTGTGCGTGGAGATAGACTTGACAAAACCAATTCCACGAAATTGTTCATCAAATTGGTGAAGCGGAAGGGATGCGCTTAACAGATCTTCAGAAGGCAATCCAATCCTCTTTGCTCATATGGTCATGGACCGCCACGCTCTTTTATCGCGAGACGCATCTTTATAGAGAAGGAAGAATGCGAACCAAATGTGCGGAAGGCAACTACGAAGAATTTGTGACCATGGAGTTGCACAAGCCACCCTTGAAGTAATCACCAATGTTGGTTGCGACGTCATCGGCTAATGTCTCCCGATCTCATTATGCCACCCATCATGGACGGTGCTCACAGCCCGAGGCGAGATGCAACATCTCCTAATACACATGAGTTGCGAGACTTTGAAAGTGGCAATCTGTACCAAAAGCATCATCGGCCAGATACAAAGCTGTGCCTTCGGTTTGTGACGCAATTACACCATTAGCAGCCCAGAATTCTCCACCTGGAACATAACCATGGGCCATTACCGCAATCTGTGTGTCTGAGACGGCCAGAGGGCTTATTCAAGCGTCCTGAATCGGCCTGGAGGCAGCTCGAGAATGACAAGCAAACATGCATCCCATACAGGACAGTGCGTGACATTCACACACCCGACTGAGGTATTCCCAAGCGAGTAGAATCACAAAAGACTCGAGTTAATGCACTCCACCATCATCACAAGGTTCACTGTCTTTACCTTCTTTCAAAGATACGGCCAGATCTTTACTATACGAGAAGGTGAAATGGCGGGTGCTCCCGATGAAGGCAGCGCAGCCTACGGAAACCCAGGCTCGCGAGACGAGTTGAAGTACCAACGAATTATGCAGGCACCGAAATAGGAGATGACGGCCTCTTATGTGGCTCGTTTAGCGAAGCAAGGGAAGGGATACACCCAGCACCGGATGATGCTAGGCGAGCCGGTATAGATGTCGATACTTCTCCCGGGTAACAAGCGGAAGCACGAGAAGCATAAGCCAAGCGAATTGTTGAGCCGGAGCGAAGCCGAGCTCGCCTGTATGACAATCACCAAGGAAGACCTAGCAGCACTTGCTGGACAAGCGGAAGGCCTCTTGTTATACCGGCTCAAGAGCCGGCAAAGAGAGCATCAAAGCACCAATCCGATTCGATTCGCGGCGCCAACCGGAATCTGGTCAGTTGGCCAAAGACCCGGGCCAATAGGAAATCCGAGGAAGCCGCGAGATCCGACACTAGCTTTTGAACATTCCTTCCCACCAGAGATCTGAACACCGGATGGTGAAGACCCAGCTGGATCAACTCCTTATCGGATTGGGAGCAGGCACCTTATTGAGACCGCTCACACATGTAGTGAGATGGCATACATTTAATCTAGGTGTTTGGGTTTCAATTGGCGAATTCATTAGCAATCTTCCGCATTAGTAACCGCCTCTCTCCCCGCCCGTTATCTTGGGCTAGCTCCGCCCAATCTGGAACACCGAACCAAGTGTATACCATGTTGTTGCCATGACTTTACTAATCGGACCTTGGAGGTGAGCGAGTCAAGAGATTCATCACTTGAAATTGTATGTATATTATCTGGCCGTGACTTGCTCAAGGTGCCAACGTGATGTTTGAGGCAGTGGAGCGATGACATTTCTTGTTTATTTGCCCTCTTTATATATATCAAATGTATAGCTTTAAAGCCAATGATCGATGTGACTCATGAAGGCTTATCTTGTATAGGCTCACTTTGGTGGCAACTTTGGGCTTCTTGAATTAAAGGCATAGTTCACCATCTTCGGGTCAGGCAGAATGTGCTCACACTCGAACCCTTCTCAGGAAGATCAAGGTCGGTCGGTGCACCGAAAAGGATCCCGCACATTAGCTTCCCTTGCGCCTTACGGGTTTAATCGCGATTGGCTCGCACACATATCGGTCTCCTTGGTCCGTGTTTCAAGGCGGGCGAATGGGGAAACCCACGAGGCCGTTACAAAGAACGCGGAATGCAGAAGCACGCCGAGACAGCGCGTCTTGCCCATCGCTGATCGCCGTCGGCCGGCGTCTCCGCTGAGCATATCAACAGCCCCCGGGCTTTGGCGCCGCCGCGATCATAGCGGTCCCACGCCCCGGAAGTCGAGTGGCGGACCGGCTTGTGACGTTCCACATCGTGACTCCAGCAGCTATCATCCCTCTGAGCAATTTCAGGCACTCTTTGACTCTCTTTTCAAAGTCCTTTTCATCTCTTTCCCTCGCAGTACTTGTTTGCTATGGAGTCTCGCCGTATTTAGCCTTGGGCGGAATTTACCGCCCGATTGGGGCTGCATTCCCAAACAACCCGACTCACTTGACGGGGCGCCTCGTGGTGCGACGAGGTCCGGGCACAGCGGGGCTCTCACCCTGTGGTACTTTGCCGAGACTTGTTCGATCTTGTGAGGGCCTTCTCCGAGACTTCTGGTCGGACGCCTTGCATGCCCGGATTCTCAAGTGCAATTTTATTTCGGTTCGCTCGCCGTTACCACAAGAGTCCTTGTAAGTTTCTTTTCTCCAAGCTGTGATATGCTTAAGTTGGCCAGGTAACCCGCTGACACGGGGTCGCGTTTCAGAAAAGCGCCGGCCGGCAGCAGCATTAGATAGAGAAAAGCACGGGGCTGACAGCAACGAGGACACGAGGTTAACCAGCCACCGATTATCGCGTGGAGTGCGAGGACTTATTATTTATGGCGATGCGGCCAACGGAGCCACGGGAGGCCATGATCAAATCCACCCAACTCCCGGGGGGTTGGGAGGGCAACGGCGATGTGGCGCCAGACGTGCCCTGCGGCCTAATGGCTTGGGCGCAACTTGCGTTCAAAAAAACTTCGATGGTTCGCGGATTACGCAATTCACACCGAAGTATTCGCATTATCGTTCTTCGCCGATCGCGAGCGAGATATCCGTTGCCGAGTCGTTTAGACATATTGAGGCAGCGAACCACCCGCGCAGTCACCGTCTCCGGGGACGGCGGAGTGGCAGCTCCTTCATTCAAGTTCCTTGGCATGAGTCACGCCGATGTTCGATGCGCCGGAGAGGTCGGCCAAGCGGGAAGCCGAAGCACGACACGCGAGGCTTCGCTTCGGGATGAGGAGGCGGACGCGAGGCCCCGTCTCAAAGCGTTGAAACCATTCTCTGGGTCGTTCTACTAGGCGAGTTTAGACAGTCCTTCGGGTTCACCTCACAGGAAACCGCTATTCGACTTCTCCTTCTCTAAATGGTAAAGTTGATGGACTTCTTGTCCTTGGTGTGGGCAAGCCACGTCGCCGCGATCCGAAAACTTCACCGGACCATTCAATCGGTAGGACGGCAGGCGGTGTGTACAAGGCGGGGGCGTAGTCAACGCCGGTGATGACTCTCAGCTTACTAGGAATTCTCGTTGAGGCCCACTTGAGTGCAATGATCTATCCCCATCGCGATAGAAATTCCAAAGATTACCGGGCACCCATCGGCCAAGGCTATAGACTGTTGGATACATCGGTGTAATGCGTCATTAGAACATCTAAGGGCACTCAGGCTGTTATTGCCTCAAACTTCCTTGGCCTAAGCGGCCATAGTCCCTCCTAAGAAGGCGGCCGCGGAGGAGACCTCCCAAACAGCTGGTCAGCAGTGTTTGAGGTCTCGTTCGATTACTTGGAATTAACCGGACAAATCGCTCCACCAACTAAGAACGGCCATGCACCACCACCCATAGAATAATGAAAGGAGCTCCTCTCGATGCATCGATCCTTTATATACGGACTGGTAAGTTTCCCGTGTTGAAGTCAAATTAGCCGCGACTCCACTCCTGATGGTGCCCTTAGAAATTAATTCCTTTAAGTTTCAGCCTTGCGACCATACTCCCCAGGAACCCAAAGCTTTGATTTCTCATAAGGTGCGGCGAATTCCTAAAAGTAACATCCGCCCGATCCCTAGTCGCGTCGTTTATAGTTGAGACTAGAATGATGTCAGTCGTCTTGAGCCCTGACTTTCGTTCTTGATTAAGTAAAACATCCTTGGCAAATGCTTTCGCTTGATTGTTCGTCTTTCATAAATCGAATTTCACCCTCGACTGTAAATCCCCAGATACTTTGGCCCATCTGTTAATCATTACTCGATCCCAAAAGTAGCAGAATGGACAAAATCCTATGATGTTATCATGCTAATGTATCAGGCGTGACTTGCTTTGAGCACTCTAATTTCTTTAAGTAATACGGAAATACGACCAGCCAATTGAGGCAGTATCGCAGTGAAAGGAATGACAATGGGTGCACACATAGTGGGCCAGTGTTAACCCAAGGTCAACTCACGAGCACCATGCAACAACTTAAATATACGCTATTGGGTGGAATTACATGTTCACTTGGCACCAGACTTGCCCTCAGTGGATCCTCGTTAAAGGGATTTGGGTGTACTCATTCCAATTACCGGACTCATAGAGCCGGGTATTGTTATTTATTGCTCACTACCTCCCGTGTCGAGTTAGGTAATTTGCGCGCCTCTTTGCCTTCCTTGGATGTGGTAGCCGTTTCTCGGGCTCCCTCTCCGGAATCGAACCCTAATTCTCAATCACCCGTCACCACCATGGTAGGCCACTATCCTACCATCCAGTTGATAGGGCAGATATTTGAATGATGCGTCGCGGCCACTAAAGTGCATGCGATCCGTCGAGTTATCATGAATCATCAATCTGGCAGGCGAATCGCGTTGACCTTTTATCTAATAAATGCATCCCTTCGGAGAAGTCGGGATTTGTTGCGTATTGACTCTAGAATTACTACGGTTATCCAGTGGTAGGTACCATCAAACAAACTATAACTGATTTAATGAGCCATTCATGATTCCATGATCTGAATTAGTTCATACTTACATGCATGGCTTAATCTTTGAAGTTGCTGTGTACTTGACTCTTGAGGATCGACCGGGTAGCATTCCTTTAAGCCCAGCAGCGCATGTGTGGAGACCCAGCAGTATGACAAGCGATCGTACATGTCTAGCGACAAAGCAGCAGATTTGGACATCGAGAGGTGTCGAGACACCCAAAGCCTTACGAATGTTCCGTATCCGAACCATCAGTATGGCAGAAGGCAGTACAACGTGCGCGCCACGGCGTGCTCGATACGAGGAGCCTTGGCCCTTCTACGATGTCCTTGTGCCAGCTCGGGGAAAATATTGGCAGCCCGGCACCGTTCTTTAGCGTAGGTAACCCAAATACGGGGTCTCCGTATAGCCATGAAAGTCTTCTTCGCGTCATTGCAGCGTGGTAATGTGGGCGGGCAGTTCCGATGCCAGCCCGGAGGCGCAGCCCTGCGACCCGATTCACAACTCACCGCGTTGTTACATCGTGGGGACAAACAACATTAAACCGACCACACCCCCGACTAACGAGGCGAAACAGCGCCAAAGCGATTTTATGTAGCCAAGCCCGCACACGCCCAGCGGCCAAAACATAAGCACCCACCCAATAGGCCTTGCGAAACTCAGAAACCGACTCTCTGTTCAACTTTTCGAGAGCGAGTGTCGTTAAGATCCACCGGAGGGCCGACCTACGGGGAACCGAGGCGGGACTCGTTATTTGGACACCACGCTCGACTCACTCAGGAATGAACATTGAAACTGTCGCCTTGATATCGATTAAACAAACACGAGAACCATATACATGGAAATAACTTGCCCCTTACTTGGTAGAACAAACTCCACATGCAGTGAAACCATTCCCACATCCGTTCCGAGCTACCAACCCCCTACTATATGGAATTGTAGTGCCCATTTGGCCACGAGTTCACCCATGTTTCACAAAAGTGAAAATCATGAATTTCAGAACCCCTCGCGGCCCCTCGCGTGAGGAGTGCCTGTCGCGGCAACCTCGCGAGACCCTCGCGCGGCACCTCGCAGCCTGGCTGCGCCACCGCGGCATCCTCGCAGCCCTGCTGCATGCAGCATCCTCGTAGCGCTTCTTTGCGCGCTTCAGCGGCACCTCGGCAAACCATCTTTGCACACGCGGCAACCAGCGCGGCCTTGGCTTTCGCACCACAGCGACCCCTGCGCAGCCATCGCTGCGCCTTGCATGGCATCCCTACGCCCAACCCCTTCACCGGGCATCGCGGCGGCACCGCCGGCGGCCATTACGCACGCGGCAACCGTCGCGAGCTTGGCTCGCCGCCTTGGCCTCGCGGCCATGCTGCGCACCTAAGCAGCCTCGCGGCCTTGCTGCACGCCGTGGCAAGCACCTCGCGGCCATCTGCGTGCGCGCGACCTCTGGCCTTGCGCCTGCGGCCCATCACTCGCGGCCCCTCGCGGCCCAGCCAGCTCGCACACTTGCCCGGCCCCTCGCGGCCATGCCTACCGGCATCACTCGGCGGCCCCCTCGCAGCCTTGACTGCGGCAGCAGCGCCTTGGCGGCCACCAGCTGCGCACCGCGCGGCCCTCGCGACATGTTGCCAGCATCTCGGCATCCCACTTGGCGGCCACGTGCTTGCACCTCGCGTATGCCCCTTGCGCTAGGCAACCCCTCCAAAGCCCTCAACATCATGCTAGCCTCCTCGAAGGCTACCTCAAGCTCGCAGGGGAGGCTAGAAGTATGGATGCGGGGGTTTGAGGTTGCTAGAGTGCAAGCATCTTGCCAGTCTGTATAGTGCAAAGATCTTGAATACGCACAACATCTTGCCCGAAGAGCTGCAGCGATCTGCCCCGGCGTCTTTTACGAAAAATGTGTCGCGTGCGCGGCAACTGCCCGGCCAGCTATGAGTGGAATGCGCGATGCTTATGGCCTGCTCTTTGCGAAAATGCGGCGCAATGCAAACCAGCAATGCCAGCGCAGGTAAACGACTGAAACTGCGAAGGCTGCAGCGTCTGCCAGCGTGAATGCCTCTTTGCCGCGTCTGGCTAAGTGAAAAACTGCCGAAATGCTGACGTAGCTCTGGCGGCTGCATGTCTTGGCTCTGAGTTGCCGCGGGTATGCGAAATAGCTGCAAATCTACCCTTGACTTCATTCATCTTGAGGATCTTACGCCAGTCACCTTGGCTATGTTGCGAGCTTTACGCGATGCGAGCAAAAGCTGCGTCGAGTGCAGCGTGCGGTATATGCCGTCTTCGCGTGAAATCTTGCTGGCGGCTGCGAGGGGATCTGCGATGCGTAACAGCTGAAGATGCGCTGATAGCGGCAGCTGCTGAAGAATGTGTGAAAATGCCGAAAGATGTATCGCGACTCTTGCCGGCTGCTAACTGCCCGGCCTAACGAAAATCATGATTTTTCACTTTTGTGAGTAGGTGAACTCATTATATAGGCATCTCTGATTATATAGTAGGGGGTTGTGCTGAAGCATGAGAATGGTTCGGCTGACTGCTAATTTGTTCCTACAATGAGGCAAGTTATTTCCATAGATATGGTTCTGGAGTTTAATATCAAGCGCGTAGCATTCTCGATAGTGTCGGCGTGGTAATCCAACGGCAGTCTGCTGGTTCTCAATTTACCTGGATCGCGACAACACCTACGAAAAATTGAGCGAAATGTTTTATCCGCGAAACTATTGAAATGAATTCTTTCTATGTTTTCCCATCTTTAAAGGCGATAACTTGGTGTCAGCCTATTAGTGGCGATTTTATTCTTGGAAAATCTCTTGATATAGCGGCGTGTGGTTAACCAATCCGGCGTGGAATGAGCTCATAGCCAGGCGTCGAACCGTCCGCCACACCTGCGTGAAACGGGCCTTTCTTGTGCCTGCGAAGATTCTGTATTTGCTACCTGCGCTTAAAAAGGCAGTCTTGGCTGTATTCCTTCGAGCTGGCGCGGCCGGGATGCTCGTGTGGCGCTGTGCCTGCATCGAGCGCGCCCGTAAGCGGCGCTGCGTGCTGATTTCGAGTTATTCGATTGAGATCTGTACGATCCATTGAGCAGGTGTCTCGACCTTCGTGTCAAGTCAGCGTTGTCGCTAGTTTGTATATGGCTTGTCGTTGCGACAGAATGTGGCGTCGTAAAAGAATGCTACAGTTGATCTGCCCTCTTTATCTCAAGGTAAGCCATGCATGTAAGTATGAACTAATTGAGCTGTGCTTTGCGAATGCTCATTAAACTCGATTATGTGATGAAGCACTACTATTCCGAGATAACGTAGTAATTAATCGTGCAACAAATCGCTTGGAAGAATGCATTTATTAGATAAAGGTCAGCAGCTCACCGTTATATTGATGATTCATGTCTTCGGCAGATCGCCGACATAAATCTTTGGCGACTTGCCTATCAGCACCGATGGTAGGAGGTGGCCTACGTGATGACGAGCGAGTCGAGGACCGGTCGGAGGCTTTGAAGCGACACCACATCAGGAAAACGACCGTGTGTAAATACCCAATCTGCGGGAGGTAGTGACAATAAATAATGCGGGCTCGCAATTGCAGTATTGGAATAGTACAATCTAAATCACTGCGGAATCTGGAGAACAATCTGCCCGGCGACGGCTATTCGGCTCAATAACATATATTTAAGTTGTTCGGTCAAGCCGTGGTGTTGGCGACGATCGCCTGGCGACCGACTGCGTCCTCCTGCGGCGATGCGTCGGCGCTATTGGCGAGGTCATACCTCGGTCTTCGCCCTTGAAGAATTAGAGTGCTCAAAGCTGCAGCCTCGCTCTGAAATACATTAGTGGTTCATAGGATTCCGATCTATTGATGACGCCGGGCTCAGTGGAGCGATCGGCGTTTGTCGTATTCCTGTCGAAGGTAGTGCGTTTATGAAGGCGAACAATAGCGAAAAGCTTGCCAGGATGTTTCGTGTCAGAAGCGAGATTGAGGCACCGAGCGATCGGTCCGTCTGAGTCTCAACCATAAACACGACGTCGGCGGATCGCCACAGGACTCGCGCGACTATGGTCAAAGTTTTTAGGTTGTCACTCAAAGTAAGCAAAGGGTGACGGAAAACGCCGCAGTGAGCACAAGCTGACTTCGACGAAGCTGCAGTCGGTACTGTGAGATTGACGACTGAAGCTCTTTCATTATTCTGGGTGGTGGTCCTATGTAAGATTCTTAGTTATTGAAGCCCATTCTCTGATTAGTCACGTTAGCGGCGGACACCGACGCTAACAACTATCTTGTCTCCTCGCCCGGCCGACTTCTTAGAGGACTATGGCAGCAGAACAGAGGACTGAGGCAATAGCAGTACGTGATCTGCTGAATGCAGGCATGTAGCGCTAGCGGCCAACAGTCCTGTACTGGCCCGGCAGGCGATAATCTTTGAAATTTCATCGTGATGGTCGTCATTGCAGTGTTGGTCTTCAACGAAGAAATTCTAATGGCGTGATCATCGGCTTCAGCCGTTGACGTCACGCGCTGTACACGCGCCGTCACTCTACGGTGAATGGTCGACGAGACACGGACCGGCGGCGTGACGACCGTGTCACGGCACGTCGCAGCATCCACTTCGAACATCGTGGAGAGAGGAGAGAGAATAAATAGCAAAGGTTTGGTAGTGGCTGTGAGTCGATGTCCGAACTGCTGCGGCGACCGAGTGTTTCCGAAAAGGCGAGGGGAGTTTCGAGCCTCCTCGCTCGAGGAAGCGGCCTCGCGTGTAAGTCAACCCGGCGCTCCGCTGGCTGACCTCGTCATCTGGAAGTATCGGCGTGAGTGCGCCGAAACGAATGAGAGGAAGCGTCACCCGCCCGTCCCGAGGAGGCGGCCAGTGTTCATCTCAATATGTCAGCGACCGGCTGGCGAATATCGGCTCTGCGCTCGATGAAGAAGCATGTAGCTCGTGCGCGAGGTGTGGTGCGAATCCGTGAACCAGCCGAGAGTTTTTGAGCAAGGATTCGCGCCCGGCCGATGGCGAGAATGCTGTGCAGGCGTCCTGTGTGCCCTGCGGGAGGACGAAATGAGGCAGAGATGATGGCCTCGTGCGCTCGTGCCTTGGTGGCATAAATAACAAGTCCTCGGCGACGACGCGCTGTTGTCGGTGATTATCAGCCAAATCATCGGCGGCGGCAGCGTGTCATTTCCCATCTAATCTTGGCGTCAAGCCGTCGGCTCGCGACCCGGAGTCGGCGGACCGGGTGGCGTATCGCTGTGGGAAGAAACCCTGGTCCCTTAATGGCGGCGGCTGGCGGGCGGCTCGGCGCGAATGCGAGTGCAAGCGTTCGAGTATGATAAGTCGGCGGACGAAGAACAAATCCTGAAGAAGCGCGCAGTGAAGGACCCGTTGTGCGGACCATCGCTGCAGTGTATCGGCGATGACTGAGCTCCTTAATCGGCGACAAGCGTCGAAACTAAACTCTGAAGCGGCGATGATAGCAAACAATACTGTGAAGATGAACACGAAAAGAGTCAAGTAACTTAATGACAGGCGAATGAAAGTAGCGATGCGCCGACCGGATGTCGAAACATTTTTAACGTGCGCCACTCGACGAAGCTCGTTGATGAGCTAAGCGGCGGCCGGGCTATTAACCATGCGTGAGCGTCGTCGTATGATCATGTGGCGTACCCAGCGCCGCGTCTCCGGCGTCTTACTCTATGCTCTGTGGCGGCACAATGGCTCGACCGGCGCGTCTTTATGAGCAAAATGACATAATGTGAATCAACGGCGATAGCGCAAGGAAAGCAATGTCGCGAGGATCCCCGAATGCGCGTGGCGCCGACCTTACGAAGGCGATGTGGCGTACTGCTCGGGCCGCAAAATGGTGAACTGCCTTTGTGTTCAGAAGCCGGTGCCGCAAAGTAATTACAGATAAACCTCATGGTCACCGTTATTATTGGCAAAGCTATGCATTGACATATATAAAGAGTAACGACAGCGAAATGTGCATCGTTCTCGTCTTGGCTGTGCGTTGGCACCTTTGAAACTGAGCTGGCATTGAATTAATTATACATACAGCCGATGATGAATACTCAAAACAATTATCTCAGGCTCGATAGCAAGAAATAAACTTTAACAACAAATGCGACGATAACGTGTTCCATCATAAAGGCGAAGAGGAAGAAGCCGATTATCTTAATGCGAAGGTATGAAATCATGATGTACCTGATGTCTGCTTTTTCGGCCGTGCAATGTCTCAGTAAAGGTGCCTCGGCTCTCAATTTGACAAATGATCGATGGGTGCCTGTCATCTGTAACGTGTCTGTGGAAGAGCTCTGGCGGGTGTCACCCTACCATTTCCTTTGGCGATCAGTGCTAACTGCTTGCATCCGCATCTGTGGTGGCGCTGGCAGGGCTCCGGGCGGCTGTGCTCTTCTCTTGCGACTCTTGAAGCGTATGGTGACATCGATGTGTACTCATAACAATACGTGGC
